# High fat diet induces microbiota-dependent silencing of enteroendocrine cells

**DOI:** 10.1101/658435

**Authors:** Lihua Ye, Olaf Mueller, Jennifer Bagwell, Michel Bagnat, Rodger A. Liddle, John F. Rawls

**Author notes:** Address correspondence to: Rodger A. Liddle < > or John F. Rawls < >.

## Abstract

Enteroendocrine cells (EECs) are specialized sensory cells in the intestinal epithelium that sense and transduce nutrient information. Consumption of dietary fat contributes to metabolic disorders, but EEC adaptations to high fat feeding were unknown. Here, we established a new experimental system to directly investigate EEC activity in vivo using a zebrafish reporter of EEC calcium signaling. Our results reveal that high fat feeding alters EEC morphology and converts them into a nutrient insensitive state that is coupled to endoplasmic reticulum (ER) stress. We called this novel adaptation “EEC silencing”. Gnotobiotic studies revealed that germ-free zebrafish are resistant to high fat diet induced EEC silencing. High fat feeding altered gut microbiota composition including enrichment of *Acinetobacter* species, and we identified an *Acinetobacter* strain sufficient to induce EEC silencing. These results establish a new mechanism by which dietary fat and gut microbiota modulate EEC nutrient sensing and signaling.

## INTRODUCTION

All animals derive energy from dietary nutrient ingestion. The energy harvested through digestion and absorption of dietary nutrients in the intestine is consumed by metabolic processes or stored as fat in adipose tissues. Excessive nutrient intake leads to metabolic disorders such as obesity and type 2 diabetes. To maintain energy homeostasis the animal must constantly monitor and adjust nutrient ingestion in order to balance metabolic needs with energy storage and energy intake. To accurately assess energy intake, animals evolved robust systems to monitor nutrient intake and communicate this dynamic information to the rest of the body. However, the physiological mechanisms by which animals monitor and adapt to nutrient intake remain poorly understood.

The primary sensory cells in the gut epithelium that monitor the luminal nutrient status are enteroendocrine cells (EECs) (Furness, Rivera, Cho, Bravo, & Callaghan, 2013). These hormone-secreting cells are dispersed along the entire gastrointestinal tract but comprise only ~1% of gut epithelial cells (Sternini, Anselmi, & Rozengurt, 2008). However, collectively these cells constitute the largest, most complex endocrine network in the body. EECs synthesize and secrete hormones in response to ingested nutrients including carbohydrates, fatty acids, peptides and amino acids (Delzenne, Cani, & Neyrinck, 2007; Moran-Ramos, Tovar, & Torres, 2012). These nutrients directly stimulate EECs by triggering a cascade of membrane depolarization, action potential firing and voltage dependent calcium entry. Increase of intracellular calcium ([Ca^2+^]_i_) can trigger the fusion of hormone-containing vesicles with the cytoplasmic membrane and hormone release (Sternini et al., 2008). The apical surface of most EECs are exposed to the gut lumen allowing them to detect ingested luminal contents (Gribble & Reimann, 2016). However, some EECs are not open to the gut lumen and reside close to the basal lamina (Hofer, Asan, & Drenckhahn, 1999; Sternini et al., 2008). These different morphological types are classified as “open” or “closed” EECs respectively, and traditionally have been thought to reflect distinct developmental cell fates. However, the transition between open and closed EEC types has not been described.

Besides morphological characterization, EECs are commonly classified by the hormones they express. More than 15 different hormones have been identified in EECs which exert broad physiological effects on gut motility, satiation, food digestion, nutrient absorption, insulin sensitivity, and energy storage (Moran-Ramos et al., 2012). EECs communicate not only through circulating hormones, but also through direct paracrine and neuronal signaling to multiple systems including the intrinsic and extrinsic nervous system, pancreas, liver and adipose tissue (Bohorquez et al., 2015; Gribble & Reimann, 2016; Kaelberer et al., 2018; Latorre, Sternini, De Giorgio, & Greenwood-Van Meerveld, 2016). EECs therefore have a key role in regulating energy homeostasis and represent the first link that connects dietary nutrient status to systemic metabolic processes.

Energy homeostasis can be influenced by many environmental factors, although diet plays the most important role. Despite efforts to reduce dietary fat intake in recent decades, the percentage of energy intake from fat remains ~33% in the US (Austin, Ogden, & Hill, 2011). High levels of dietary fat have a dominant effect on energy intake and adiposity (Hu et al., 2018) and have been implicated in the high prevalence of human metabolic disorders worldwide (Ludwig, Willett, Volek, & Neuhouser, 2018; Oakes, Cooney, Camilleri, Chisholm, & Kraegen, 1997; Panchal et al., 2011). The effects of a high fat diet on peripheral tissues like pancreatic islets, liver and adipose tissue have been studied extensively (Green & Hodson, 2014; Kahn, Hull, & Utzschneider, 2006). It is also well appreciated that consumption of a high fat diet affects the microbial communities residing in the intestine, commonly refered to as the gut microbiota (David et al., 2014; Hildebrandt et al., 2009; Murphy et al., 2010; Turnbaugh, Backhed, Fulton, & Gordon, 2008; Wong et al., 2015). Gnotobiotic animal studies also demonstrated that gut microbiotas altered by high fat diet can promote adiposity and insulin resistance (Ridaura et al., 2013; Turnbaugh et al., 2008; Turnbaugh et al., 2006), but the underlying mechanisms are incompletely understood. Notably, despite the importance of EECs in nutrient monitoring and systemic metabolic regulation, it remains unknown how a high fat diet might impact EECs function and whether the gut microbiota play a role in this process.

A major problem in studying the effects of diet on EEC physiology has been the lack of *in vivo* techniques for studying these rare cells in an intact animal. Historically, *in vivo* EEC function has been studied by measuring hormone levels in blood following luminal nutrient stimulation (Goldspink, Reimann, & Gribble, 2018). However, many gastrointestinal hormones have very short half-lives and peripheral plasma hormone levels do not mirror real-time EEC function (Cuenco et al., 2017; Druce et al., 2009; Kieffer, McIntosh, & Pederson, 1995). EEC function has been measured *in vitro* via cell and organoid culture models using electrophysiological cellular recordings and fluorescence-based calcium imaging (Kaelberer et al., 2018; Kay, Boissy, Russnak, & Candido, 1986; Reimann et al., 2008). However, these *in vitro* models are not suited for modeling the effect of diet and microbiota on EEC function as they are unable to reproduce the complex *in vivo* environment that involves signals from neighboring cells like enterocytes, enteric nerves, blood vessels and immune cells. Moreover, *in vitro* culture systems are unable to mimic the dynamic and complex luminal environment that contains food and microbiota. Therefore, to fully understand the effects of diet and microbiota on EEC function, it is necessary to study EECs *in vivo*.

In this study, we utilized the zebrafish model to investigate the impact of dietary nutrients and microbiota on EEC function. The development and physiology of the zebrafish digestive tract is similar to that of mammals (Wallace, Akhter, Smith, Lorent, & Pack, 2005; Wallace & Pack, 2003). Zebrafish hatch from their protective chorions at 3 days post-fertilization (dpf) and microbial colonization of the intestinal lumen begins shortly thereafter (Rawls, Mahowald, Goodman, Trent, & Gordon, 2007). The zebrafish intestine becomes completely patent by 4 dpf and feeding and digestion begin around 5 dpf. The zebrafish intestine develops most of the same differentiated epithelial cell types as observed in mammals, including absorptive enterocytes, mucus-secreting goblet cells, and EECs (Ng et al., 2005; Wallace et al., 2005; Wallace & Pack, 2003). Digestion and absorption of dietary fat occur primarily in enterocytes within the proximal intestine of the zebrafish (Quinlivan & Farber, 2017) (yellow area in Figure 1D). These conserved aspects of intestinal epithelial anatomy and physiology are associated with a conserved transcriptional regulatory program shared between zebrafish and mammals (Lickwar et al., 2017). To monitor EEC activity in zebrafish, we used a genetically encoded calcium indicator (Gcamp6f) expressed under control of an EEC gene promoter. The excitability of EECs upon luminal stimulation could be measured using *in vivo* fluorescence-based calcium imaging. By combining this *in vivo* EEC activity assay with diet and gnotobiotic manipulations, we show here that specific members of the intestinal microbiota mediate a novel physiologic adaption of EECs to high fat diet.

**Figure 1.**
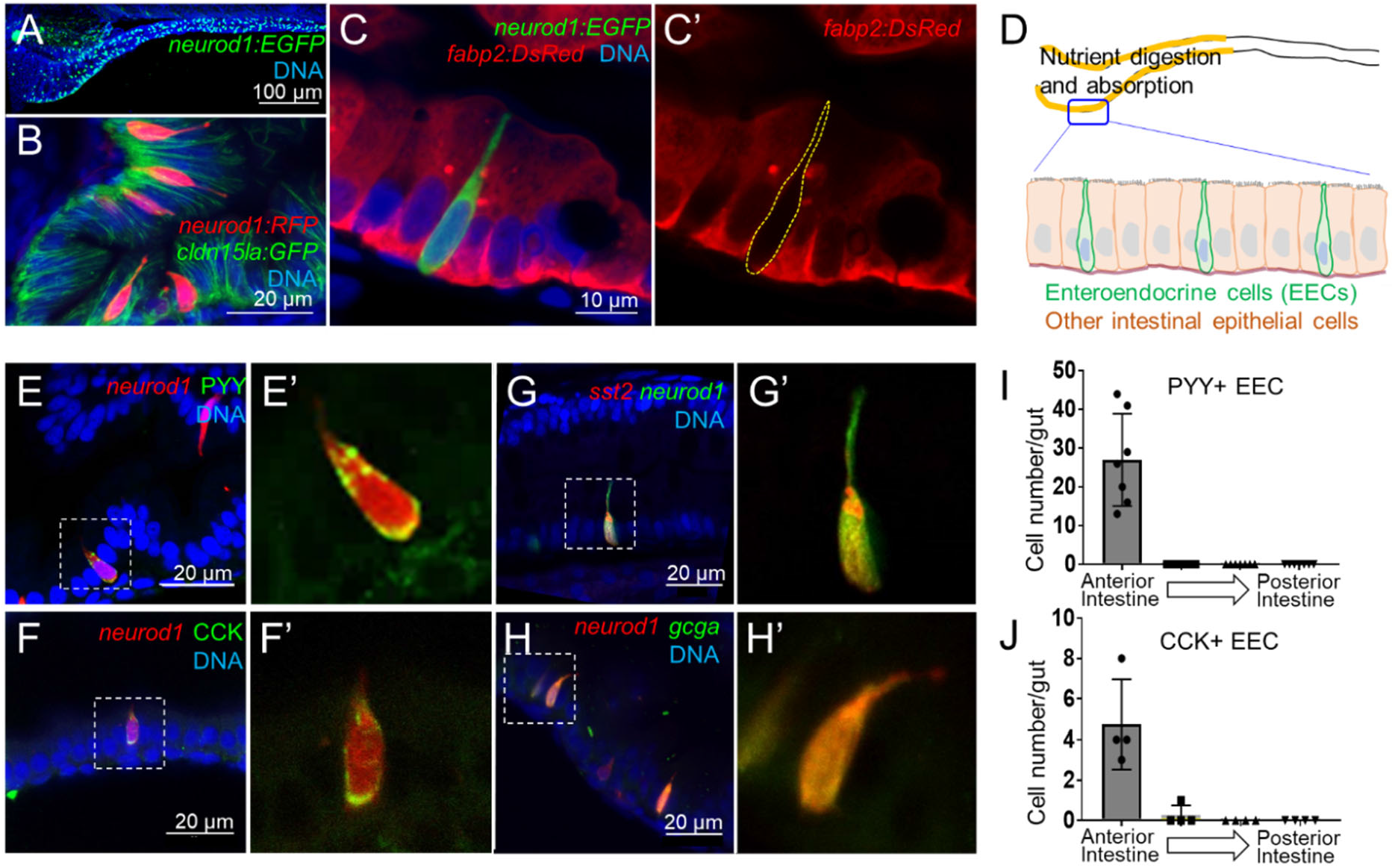
Identification of *neurod1*+ zebrafish enteroendocrine cells (EECs). (A) Confocal projection of zebrafish EECs marked by *Tg(neurod1:EGFP)* transgenic line. (B) Confocal projection of zebrafish EECs marked by *Tg(neurod1:RFP)* transgenic line. *TgBAC(cldn15la:GFP)* marks intestinal epithelial cells. (C-C’) Confocal image of zebrafish EECs marked by *Tg(neurod1:EGFP)* transgenic line. Note that *neurod1*+ EECs do not express the enterocyte marker *fabp2* which is marked by *Tg(fabp2:DsRed)*. (D) Schematic diagram of 6 dpf larval zebrafish intestine. The anterior region of the zebrafish intestine that is largely responsible for nutrient absorption is highlighted in yellow. (E-F) Confocal image of *neurod1*+ EECs stained for PYY (E, E’) and CCK (F, F’). (G-H) Confocal image of *neurod1*+ EECs express somatostatin [marked by *Tg(sst2:DsRed)* in G, G’] and proglucagon hormones [marked by *Tg(gcga:EGFP)* in H, H’]. (I-J) Quantification of PYY+ (n=7) and CCK+ (n=4) EECs in 6 dpf zebrafish larval intestines.

## RESULTS

### Establishing methods to study enteroendocrine cell function using an *in vivo* zebrafish model

We first developed an approach to identify and visualize zebrafish EECs *in vivo*. Previous mouse studies have shown that the transcription factor NeuroD1 plays an essential role to restrict intestinal progenitor cells to an EEC fate (H. J. Li, Ray, Singh, Johnston, & Leiter, 2011; Ray & Leiter, 2007), and is expressed in almost all EECs without expression in other intestinal epithelial cell lineages (H. J. Li, Kapoor, Giel-Moloney, Rindi, & Leiter, 2012; Ray, Li, Metzger, Schule, & Leiter, 2014). We used transgenic zebrafish lines expressing fluorescent proteins under control of regulatory sequences from the zebrafish *neurod1* gene, *Tg(-5kbneurod1:TagRFP)* (McGraw, Snelson, Prendergast, Suli, & Raible, 2012) and *TgBAC(neurod1:EGFP)* (Trapani, Obholzer, Mo, Brockerhoff, & Nicolson, 2009). We found that both lines labeled cells in the intestinal epithelium of 6 dpf zebrafish (Fig. 1A-B, Fig. S1A), and that these *neurod1*+ cells do not overlap with goblet cells and express the intestinal secretory cell marker 2F11 (Crosnier et al., 2005) (Fig. S1C-E). To further test whether these *neurod1*+ cells in the intestine label secretory but not absorptive cell lineages, we crossed *Tg(-5kbneurod1:TagRFP)* with the Notch reporter line *Tg(tp1:EGFP)* (Parsons et al., 2009). Activation of Notch signaling is essential to restrict intestinal progenitor cells to an absorptive cell fate (Crosnier et al., 2005; H. J. Li et al., 2012), suggesting *tp1*+ cells may represent enterocyte progenitors. In accord, we found that *neurod1*+ cells in the intestine do not overlap with *tp1*+ cells (Fig. S1B). Additionally, our results demonstrated that *neurod1*+ cells in the intestine do not overlap with the mature enterocyte marker *ifabp/fabp2* (Kanther et al., 2011)(Fig. 1C). These results suggested that, similar to mammals, *neurod1* expression in the zebrafish intestine occurs specifically in EECs. In addition, using EdU labeling at 5 dpf zebrafish larvae, we found that EECs in the intestine are post-mitotic and require about 30 hours to differentiate from proliferating progenitors (Fig. S2A-F).

Hormone expression is a defining feature of EECs, so we next evaluated the expression of four hormones in *neurod1*+ EECs in 6 dpf zebrafish larvae: pancreatic peptide YY (PYY), cholecystokinin (CCK), somatostatin (*Tg(sst2:RFP*),(Z. Li, Wen, Peng, Korzh, & Gong, 2009)) and glucagon (precursor to glucagon-like peptides GLP-1 and GLP-2; *Tg(gcga:EGFP)*, (Ye, Robertson, Hesselson, Stainier, & Anderson, 2015)) (Fig. 1E-H). We found that PYY and CCK hormones, which are important for regulating fat digestion and feeding behavior, are only expressed in EECs in the proximal intestine where dietary fats and other nutrients are digested and absorbed (Carten, Bradford, & Farber, 2011; Farber et al., 2001) (Fig. 1I-J). In contrast, somatostatin expression occurred in EECs along the whole intestine and glucagon expressing EECs were found along proximal and mid-intestine but excluded from the distal intestine (Fig. S1F-G). The regionalization of EEC hormone expression may reflect the functional difference of EECs and other epithelial cell types along the zebrafish intestine (Lickwar et al., 2017).

EECs are specialized sensory cells in the intestinal epithelium that can sense nutrient stimuli derived from the diet such as glucose, amino acids and fatty acids. Upon receptor-mediated nutrient simulation, EECs undergo membrane depolarization that results in transient increases in intracellular calcium that in turn induce release of hormones or neurotransmitters (Goldspink et al., 2018). Therefore, the transient increase in intracellular calcium concentration is an important mediator and indicator of EEC function. To investigate EEC function in zebrafish, we utilized a *neurod1:Gcamp6f* transgenic zebrafish model (Rupprecht, Prendergast, Wyart, & Friedrich, 2016), in which the calcium-dependent fluorescent protein *Gcamp6f* is expressed in EECs under control of the −5kb *neurod1* promoter (McGraw et al., 2012). Using this transgenic line, we established an *in vivo* EEC activity assay system which permitted us to investigate the temporal and spatial activity of EECs *in vivo*. Briefly, unanesthetized *Tg(neurod1:Gcamp6f)* zebrafish larvae were positioned under a microscope objective and a solution containing a stimulus was delivered onto their mouth. The stimulus was then taken up into the intestinal lumen and EEC Gcamp6f activity was recorded simultaneously (Fig. 2A; see Methods and Fig. S3 for further details). Using this EEC activity assay, we first tested if zebrafish EECs were activated by fatty acids. We found that palmitate, but not the BSA vehicle control, activated a subset of EECs (Fig. 2B-F, supplemental video 1). Similar patterns of EEC activation in the proximal intestine were induced by the fatty acids linoleate and dodecanoate; whereas, the short chain fatty acid butyrate did not induce EEC activity (Fig. 2D). The ability of EECs in the proximal intestine to respond to fatty acid stimulation is interesting because that region is the site of dietary fatty acid absorption (Carten et al., 2011). In this region EECs express CCK which regulates lipase and bile secretion and PYY which regulates food intake (Fig. 1IJ). Our results further establish that activation by fatty acids is a conserved trait in zebrafish and mammalian EECs.

**Figure 2.**
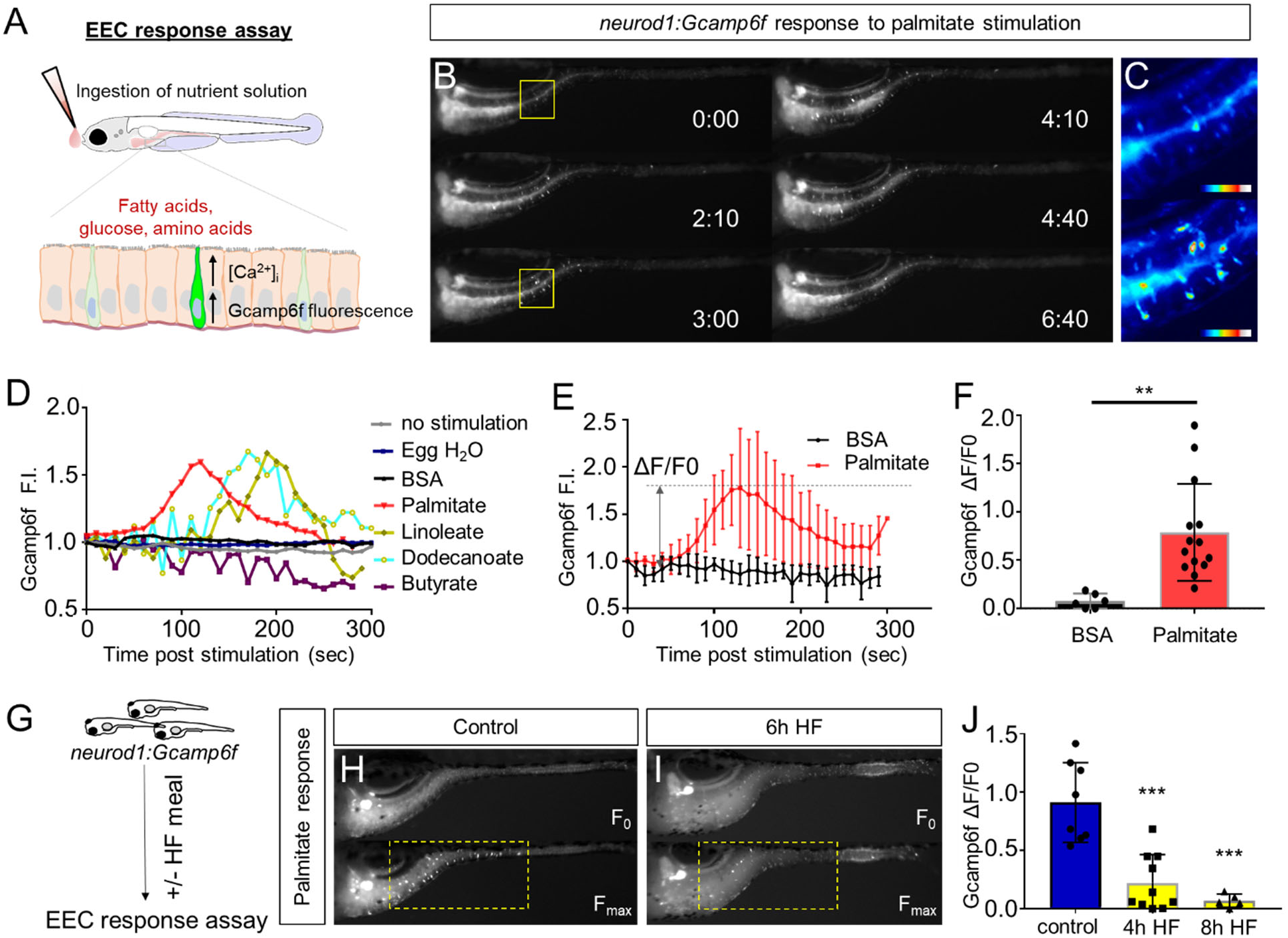
High fat feeding impairs the EEC calcium response toward palmitate stimulation. (A) Measurement of the EEC response to nutrient stimulation using *Tg(neurod1:Gcamp6f)*. (B) Time lapse image of the EEC response to BSA conjugated palmitate stimulation in *Tg(neurod1:Gcamp6f)* using the EEC response assay. Note that palmitate responsive EECs are primarily in the proximal intestine. (C) Heat map image indicating the EEC calcium response at 0 and 3 minutes post palmitate stimulation from the highlighted area in B. (D) Change in Gcamp6f relative fluorescence intensity in 5 minutes with no stimulation or stimulation with egg water, BSA vehicle, palmitate, linoleate, dodecanoate or butyrate. Note that only palmitate, linoleate and dodecanoate induced EEC calcium responses. (E, F) Change in Gcamp6f relative fluorescence intensity in BSA stimulated (n=4) and palmitate stimulated (n=5) animals. (G) Measurement of EEC calcium responses to palmitate stimulation following 4 - 8 hours of high fat (HF) meal feeding in 6 dpf *Tg(neurod1:Gcamp6f)* larvae. (H, I) Representative images of the EEC response to palmitate stimulation in control larvae (without HF meal feeding, H) and 6 hours of HF feeding (I). (J) Measurement of EEC calcium responses to palmitate stimulation in control embryos following 4 and 8 hours HF feeding. Student t-test was used in F and One-way ANOVA with post-hoc Tukey test was used in J. ** P<0.01, *** P<0.001

### High fat feeding impairs enteroendocrine cell nutrient sensing

The vast majority of previous studies on EECs in all vertebrates has focused on acute stimulation with dietary nutrients including fatty acids. In contrast, we have very little information on the adaptations that EECs undergo during the postprandial process. To address this gap in knowledge, we applied an established model for high fat meal feeding in zebrafish (Carten et al., 2011; Semova et al., 2012). In this high fat (HF) meal model, zebrafish larvae are immersed in a solution containing an emulsion of chicken egg yolk liposomes which they ingest for a designated amount of time prior to postprandial analysis using our EEC activity assay (Fig. 2G). Importantly, ingestion of a HF meal does not prevent subsequent nutrient stimuli such as fatty acids to be ingested and distributed along the length of the intestine (Fig. S4A-F). To our surprise, we found that the ability of EECs in the proximal intestine to respond to palmitate stimulation in our EEC activity assay was quickly and significantly reduced after 6 hours of HF meal feeding (Fig. 2H-J, supplemental video 2).

We next sought to test if HF feeding only impairs EEC sensitivity to fatty acids or if there are broader impacts on EEC nutrient sensitivity. First, we investigated EEC responses to glucose stimulation. Similar to fatty acids, glucose stimulation activated EECs only in the proximal intestine of the zebrafish under unfed control conditions (Fig. 3A, B, supplemental video 1). Previous mammalian cell culture studies reported that glucose-stimulated elevation of intracellular calcium concentrations and hormone secretion in EECs is dependent upon the EEC sodium dependent glucose cotransporter 1 (Sglt1), an apical membrane protein that is expressed in small intestine and renal tubles and actively transports glucose and galactose into cells (Song, Onishi, Koepsell, & Vallon, 2016). Similarly, we found that Sglt1 is expressed on the apical surface of zebrafish intestinal epithelial cells including enterocytes and EECs (Fig. 3E). In addition, co-stimulation with glucose and phlorizin, a chemical inhibitor of Sglt1, blocked the EEC activation induced by glucose (Fig. 3F-G). Consistently, the EEC response to glucose stimulation was significantly increased by the addition of NaCl in the stimulant solution which will facilate sodium gradient dependent glucose transport by Sglt1 (Fig. 3C). In addition, zebrafish EECs also responded to the other Sglt1 substrate, galactose, but not fructose (Fig. 3D). These results suggest that glucose can induce EEC activity in a Sglt1 dependent manner in the zebrafish intestine.

**Figure 3.**
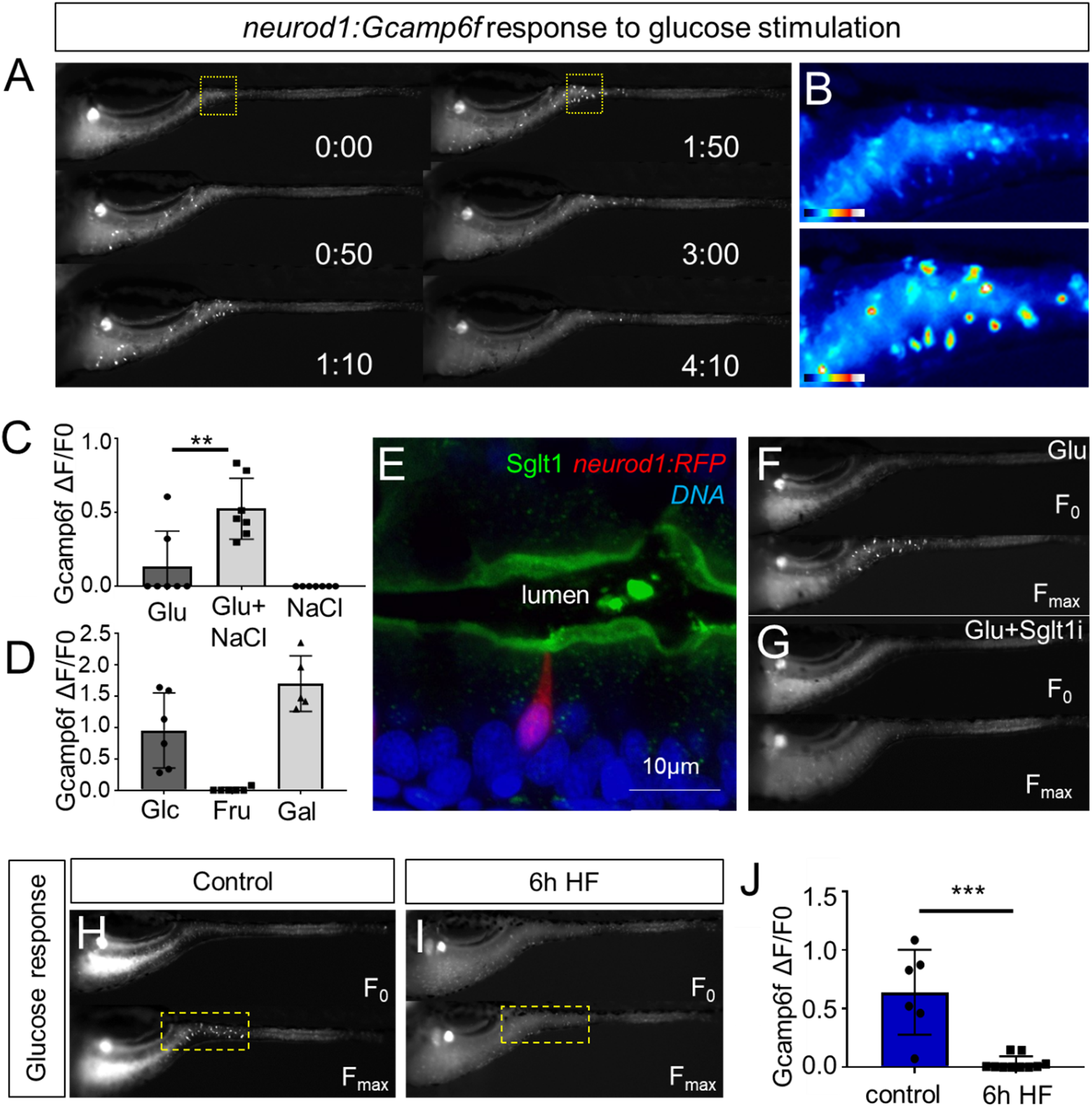
High fat feeding impairs EECs’ calcium response to glucose stimulation. (A) Time lapse images of the EEC response to glucose (500 mM, dissolved in 100 mM NaCl solution) in *Tg(neurod1:Gcamp6f)* using the EEC response assay. (B) Heat map image indicating the EEC calcium response at 0 and 110 seconds post glucose stimulation from the highlighted area in A. (C) Measurement of the EEC calcium response when stimulated with glucose (500 mM) dissolved in water or 100 mM NaCl vehicle. Note that the presence of NaCl significantly increased the glucose induced EEC calcium response. (D) Measurement of the EEC calcium response when stimulated with glucose (500mM), fructose (500mM) and galactose (500mM). All of these stimulants were dissolved in 100 mM NaCl solution. Note that only glucose and galactose induced the EEC calcium response. (E) Confocal image of 6 dpf zebrafish intestine stained with Sglt1 antibody. EECs were marked by *Tg(neurod1:RFP)*. Note that Sglt1 is located on the apical side of intestinal cells. (F, G) Representative image of the EEC calcium response in *Tg(neurod1:Gcamp6f)* when stimulated with 500 mM glucose or 500 mM glucose with the Sglt1 inhibitor (0.15mM phloridzin). Note in G that when co-stimulated with glucose and Sglt1 inhibitor, the intestine appeared to dilate but no EEC activation was observed. (H,I) Representative image of the EEC calcium response to glucose stimulation in control larvae without high fat (HF) meal feeding (H) and 6 hours HF fed larvae (I). (J) Quantification of the EEC calcium response to glucose stimulation in control and 6 hours HF fed larvae. Student t-test was used in C,J. ** P<0.01, *** P<0.001

We then examined if HF feeding impaired subsequent EEC responses to glucose, as we had observed for fatty acids (Fig.2G-J). Indeed, HF feeding significantly reduced EECs’ response to subsequent glucose stimulation (Fig. 3H-J, supplemental video 3). We extended these studies to investigate zebrafish EEC responses to amino acids. Among the twenty major amino acids we tested, we only observed significant EEC activity in response to cysteine stimulation under control conditions (Fig. S5A-B, supplemental video 1). However, in contrast to the fatty acid and glucose responses, zebrafish EECs that respond to cysteine were located primarily in the mid intestine (Fig. S5A-B) and HF meal ingestion did not significantly impair subsequent EEC responses to cysteine (Fig. S5C-E). These results collectively indicate that HF feeding impairs the function of palmitate and glucose responsive EECs in the proximal intestine, the region where fat absorption take place.

### High fat feeding induces morphological adaption in enteroendocrine cells

To further investigate how HF feeding impacts zebrafish EECs, we leveraged the transparency of the zebrafish to permit morphologic analysis of EECs. In zebrafish under control conditions, most EECs are in an open-type morphology (Fig. 1B-G) with an apical process that extends to the intestinal lumen, allowing them to directly interact with the contents of the intestinal lumen (Fig. 4A). When we examined the proximal zebrafish intestine after 6 hours of HF feeding, we discovered that most EECs had adopted a closed-type morphology that apparently lacked an apical extension and no longer had access to the lumenal contents (Fig. 4B, Fig.S6A-C). We first speculated this shift from open-type to closed-type EEC morphology may be due to cell turnover and loss of open-type EECs and replacement with newly differentiated closed-type EECs. To test this possibility, we created a new *Tg(neurod1:Gal4); Tg(UAS:Kaede)* photoconversion tracing system in which UV light can be used to convert the Kaede protein expressed in EECs from green to red emission (Fig. S7A-C). This allowed us to label all existing differentiated *neurod1*+ EECs by UV light photoconversion immediately before HF feeding (Fig. S7G), so that pre-existing EECs emit red and green Kaede fluorescence and any newly differentiated EECs emit only green Kaede fluorescence (Fig. S7D-E). However, we did not observe the presence of any green EECs following HF feeding (Fig. S7F-G). To test whether HF feeding induced EEC apoptosis, we used an *in vivo* apoptosis model in which *Tg(ubb:sec5A-tdTomato)* (Scott T. Espenschied, 2019) zebrafish were crossed with *TgBAC(neurod1:EGFP)* allowing us to determine if apoptosis occurred in EECs (Fig. S8A-B). However, we did not detect activation of apoptosis in closed-type EECs following high fat diet feeding (Fig. S8C). These results suggest that the striking change in EEC morphology during HF feeding is not due to EEC turnover but is instead due to adaptation of the existing EECs.

**Figure 4.**
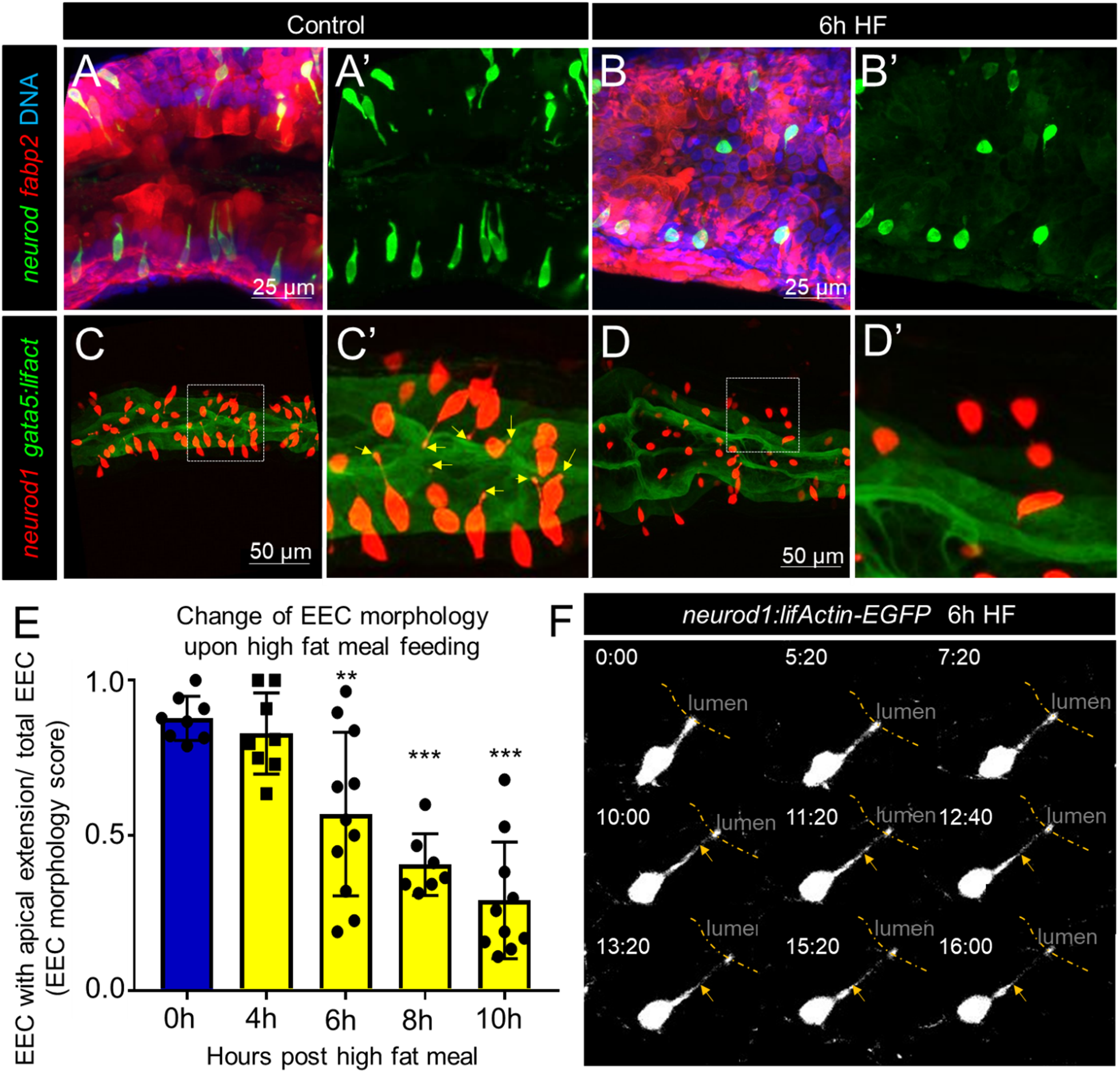
Enteroendocrine cells lose their apical extensions following high fat feeding. (A-B) Confocal projection of 6 dpf zebrafish intestine in control (A, A’) and 6 hours high fat fed larvae (B, B’). Enteroendocrine cells are marked by *Tg(neurod1:EGFP)* and the enterocytes are marked by *Tg(fabp2:DsRed)*. (C-D) Confocal projection of 6 dpf zebrafish intestine in control (C,C’) and 6 hours high fat fed larvae(D,D’). The enteroendocrine cells are marked by *Tg(neurod1:RFP)* and the apical region of intestine cells are marked by *Tg(gata5:lifActin-EGFP)*. Note that in control intestine, the enteroendocrine cells have extensions that touch the apical lumen (yellow arrow in C’). Such apical extensions in enteroendocrine cells are lost following high fat meal feeding (D, D’). (E) Quantification of EEC morphology in control and 4-10 hours high fat fed zebrafish larvae in *Tg(gata5:lifActin-EGFP)*;*Tg(neurod1:TagRFP)* double transgenic zebrafish. The EEC morphology score is defined as the ratio of the number of EECs with apical extensions over the number of total EECs. (F) Time lapse images showing loss of EEC the apical extension in 6 hours high fat fed larvae using *Tg(neurod1:lifActin-EGFP)*. One-way ANOVA with post-hoc Tukey test was used in E for statistical analysis. **P<0.01, ***P<0.001.

To analyze this adaptation of EEC morphology in greater detail, we used a new transgenic model *TgBAC(gata5:lifActin-EGFP)* together with the *Tg(-5kbneurod1:TagRFP)* line. In these animals, the apical surface of EECs and other intestinal epithelial cells can be labeled by *gata5:lifActin-EGFP* and the cytoplasmic extension of EECs to the apical lumen can be visualized and quantified through z-stack confocal imaging of the proximal intestine (Supplemental video 4). We measured the ratio of EECs with apical extensions to the total number of EECs, and defined that ratio as an “EEC morphology score”. In control embryos, most EECs are open-type and the morphology score is near 1 (Fig. 4E). We found that the EEC morphology score gradually decreased upon high fat feeding (Fig. 4E, supplemental video 5), indicating that EECs had changed from an open-type to closed-type morphology. To further analyze the dynamics of the EEC apical response, we generated a new transgenic line *Tg(-5kbneurod1:lifActin-EGFP)*(Fig. S9 A-C). Using *in vivo* confocal time-lapse imaging in *Tg(-5kbneurod1:lifActin-EGFP)* zebrafish, we confirmed that EEC apical processes undergo dynamic retraction after HF feeding (Fig. 4F), which was not observed in control animals (Fig. S9 C-D, supplemental video 6). Interestingly, EECs in the distal intestine retained their open-type morphology following HF feeding (Fig. S6 F-H), suggesting the adaptation from open- to closed-type EEC morphology is a specific response of EECs in the proximal intestine. This suggests that this EEC morphological adaption upon HF feeding is associated with impairment of EEC sensitivity to subsequent exposure to nutrients such as palmitate and glucose. We operationally define this novel EEC morphological and functional postprandial adaption to HF feeding as “EEC silencing”.

### Activation of ER stress following high fat feeding leads to EEC silencing

We next sought to identify the mechanisms underlying HF feeding-induced EEC silencing. Quantitative RT-PCR assays in dissected zebrafish digestive tracts revealed that HF feeding broadly increased expression of EEC hormones (Fig. 5A). The largest increases were *pyyb* and *ccka* (Fig. 5A), both of which are expressed by EECs in the proximal zebrafish intestine (Fig.1) and are important for the response to dietary lipid. However, HF feeding did not significantly alter expression of EEC specific transcription factors (*neurod1*, *pax6b*, *isl1*), nor the total number of EECs per animal (Fig. 5A, C). These data suggested that HF feeding challenges the existing EECs to increase hormone synthesis and secretion, perhaps in response to depletion of pre-existing hormone granules. We speculated that this increase in hormone synthesis might place an elevated demand and stress on the endoplasmic reticulum (ER), the organelle where hormone synthesis takes place. ER stress is known to activate a series of downstream cell signaling responses called the Unfolded Protein Response (UPR) (Hetz, 2012; Xu, Bailly-Maitre, & Reed, 2005). Increased misfolded protein and induction of ER stress activates ER membrane sensors Atf6, Perk and Ire1 (Hetz, 2012; Xu et al., 2005). The activated ER stress sensor Ire1 then splices mRNA encoding the transcription factor Xbp1, which in turn induces expression of target genes involved in the stress response and protein degradation, folding and processing (Yoshida, Matsui, Yamamoto, Okada, & Mori, 2001). Using quantitative RT-PCR analysis in dissected zebrafish digestive tracts, we found that HF feeding increased expression of UPR genes including chaperone proteins Gpr94 and Bip (Fig. 5B). To investigate whether ER stress is activated in EECs, we took advantage of a transgenic zebrafish *Tg(ef1α:xbp1δ-gfp)* that permits visualization of ER stress activation by expressing GFP only in cells undergoing *xbp1* splicing (J. Li et al., 2015). We crossed *Tg(ef1α:xbp1δ-gfp)* with *Tg(-5kbneurod1:TagRFP)* zebrafish and found that zebrafish larvae fed a HF meal, but not control larvae, displayed a significant induction of GFP in *neurod1*+ EECs (Fig. 5 J, K, O). Next, we tested if activation of ER stress in EECs is required for EEC silencing. Whereas HF feeding normally reduces the EEC morphology score, this did not occur in zebrafish treated with tauroursodeoxycholic acid (TUDCA), a known ER stress inhibitor (Uppala, Gani, & Ramaiah, 2017; Vang, Longley, Steer, & Low, 2014) (Fig. 5 L-N, R).

**Figure 5.**
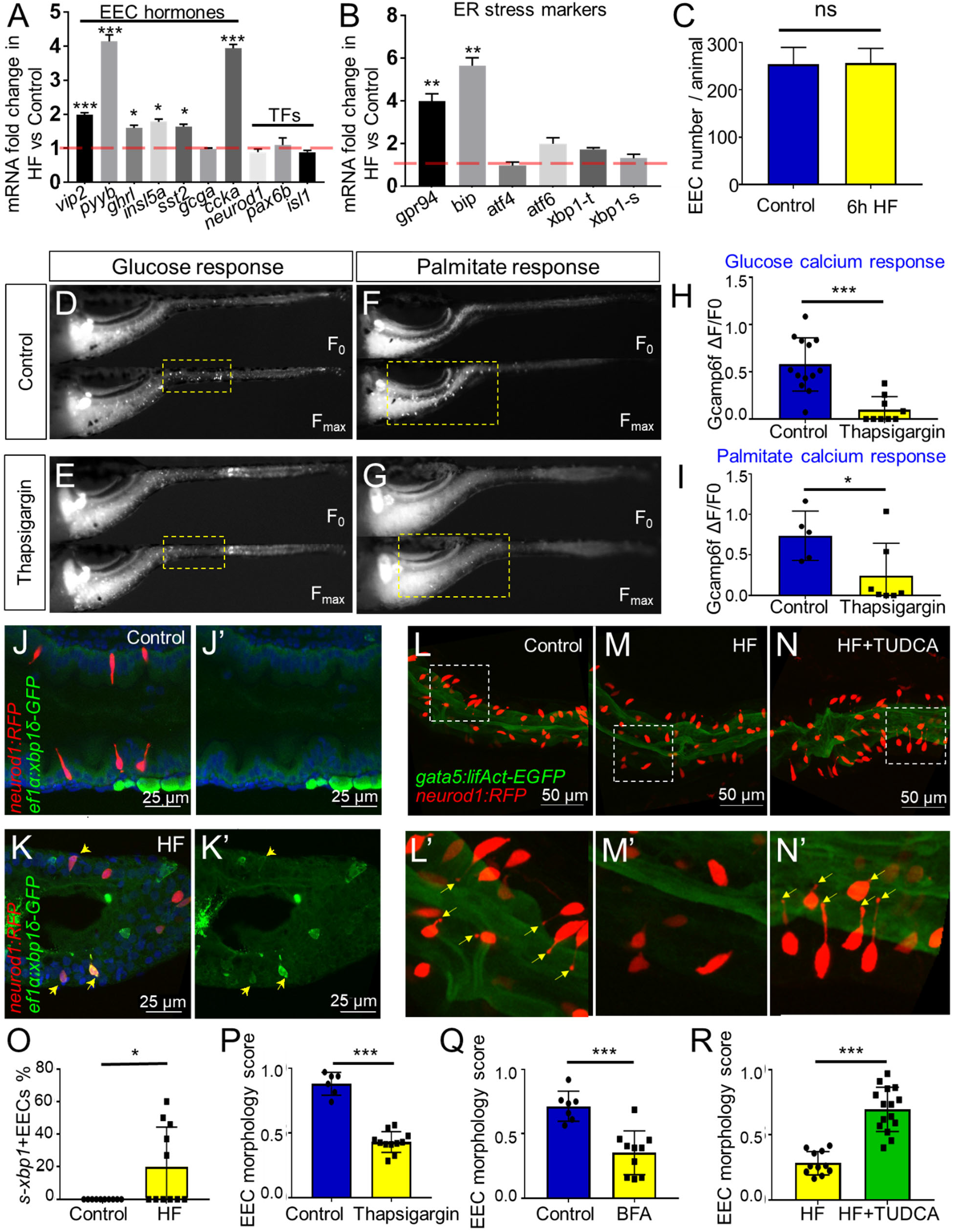
Activation of ER stress following high fat feeding leads to EEC silencing. (A-B) Quantitative real-time PCR measurement of relative mRNA levels from dissected tracts in control (n=4, each replicate from 20 pooled fish sample, 3 technique replicates) and 6 hours high fat (HF) meal larvae (n=4, each replicate from 20 pooled fish sample, 3 technique replicates). The plot indicates the fold increase of relative mRNA expression of indicated genes. (C) Quantification of total EEC number in control (n=8) and 6 hours HF fed larvae (n=6). (D-G) Representative images of the EEC calcium response to glucose or palmitate stimulation in control (D, F) and 2 hours thapsigargin (ER stress inducer, 1 µM) treated larvae (E, G). (H, I) Quantification of the EEC calcium response toward glucose (H) and palmitate (I) in control and 2 hours thapsigargin (Tg, 1 µM) treated larvae. (J-K) Confocal images of control (J, J’) and 6 hours HF fed (K, K’) larval zebrafish intestines. The EECs are marked by *Tg(neurod1:TagRFP)*, the activation of ER stress is marked by *Tg(ef1α:xbp1δ-GFP)* and DNA is stained with Hoechst 33342 (blue). Yellow arrows indicate EECs with *xbp1* activation. (L-N) Confocal projection of zebrafish intestine in control (L, L’), 10 hours HF fed (M, M’) and 10 hours HF fed treated with 0.5 mM TUDCA. EECs are marked with *Tg(neurod1:TagRFP)* and intestine apical lumen is marked with *Tg(gata5:lifActin-EGFP)*. EECs’ apical extension is indicated with yellow arrows in (L’, N’). (O) Quantification of *s-xbp1*+ EECs percentage in control and 6 hours HF fed zebrafish larvae represented in J and K. (P, Q) Quantification of EEC morphology score in control and 10 hours thapsigargin (0.75µM) or brefeldin A (BFA, 9µM) treated larvae *Tg(gata5:lifActin-EGFP)*;*Tg(neurod1:TagRFP)* double transgenic line. (R) Quantification of the EEC morphology score in zebrafish larvae following 10 hours HF feeding and 10 hours HF feeding with 0.5 mM TUDCA represented in L-N. Student t-est was performed for statistic analysis. * P<0.05, **P<0.01, *** P<0.001.

To further confirm the hypothesis that ER stress activation can lead to EEC silencing, we tested if induction of ER stress is sufficient to cause EEC silencing independent of HF feeding. We treated 6 dpf *Tg(neurod1:Gcamp6f)* zebrafish larvae with thapsigargin, a chemical compound commonly used to induce ER stress by interrupting ER calcium storage and protein folding (Samali, Fitzgerald, Deegan, & Gupta, 2010), and then performed the EEC response assay. Thapsigargin treatment reduced the EEC calcium response to both glucose and palmitate (Fig. 5D-I) and decreased the EEC morphology score, both key phenomena associated with EEC silencing (Fig. 5P). To confirm this result, we tested a second ER stress inducer brefeldin A (BFA), which inhibits anterograde ER export to Golgi and blocks protein secretion (Donaldson, Cassel, Kahn, & Klausner, 1992; Klausner, Donaldson, & Lippincott-Schwartz, 1992). Similar to thapsigargin, treatment with BFA significantly decreased the EEC morphology score (Fig. 5Q). These results support a working model wherein increased hormone synthesis and secretion following HF feeding induces ER stress in EECs which leads to EEC silencing.

### Blocking fat digestion and absorption inhibits EEC silencing following high fat feeding

We next sought to explore the physiological mechanisms within the gut lumen that may lead to EEC silencing after HF feeding. We reasoned that induction of ER stress in EECs after a HF meal is likely caused by over-stimulation with fatty acids and other nutrients derived from the meal. Fatty acids are liberated from dietary triglycerides in the gut lumen through the activity of lipases, so we predicted that lipase inhibition would block EEC silencing normally induced by HF feeding. We therefore treated zebrafish larvae with orlistat, a broad-spectrum lipase inhibitor commonly used to treat obesity (Ballinger, 2000; Hill et al., 1999). We found that treatment of *Tg(neurod1:Gcamp6f)* zebrafish with orlistat during HF feeding significantly increased the ability of EECs to subsequently respond to glucose and palmitate (Fig. 6 A-F). Next, we investigated the effect of orlistat on EEC morphology during HF feeding in *Tg(gata5:lifActin-EGFP); Tg(-5kbneurod1:TagRFP)* zebrafish. We found that following 10 hours of HF feeding, EECs in control animals had switched from an open-type to a closed-type morphology and significantly reduced the EEC morphology score (Fig. 6 G, N). By contrast, treatment with orlistat prevented HF induced EEC morphological changes (Fig. 6 H, N), suggesting lipase activity is required for EEC silencing.

**Figure 6.**
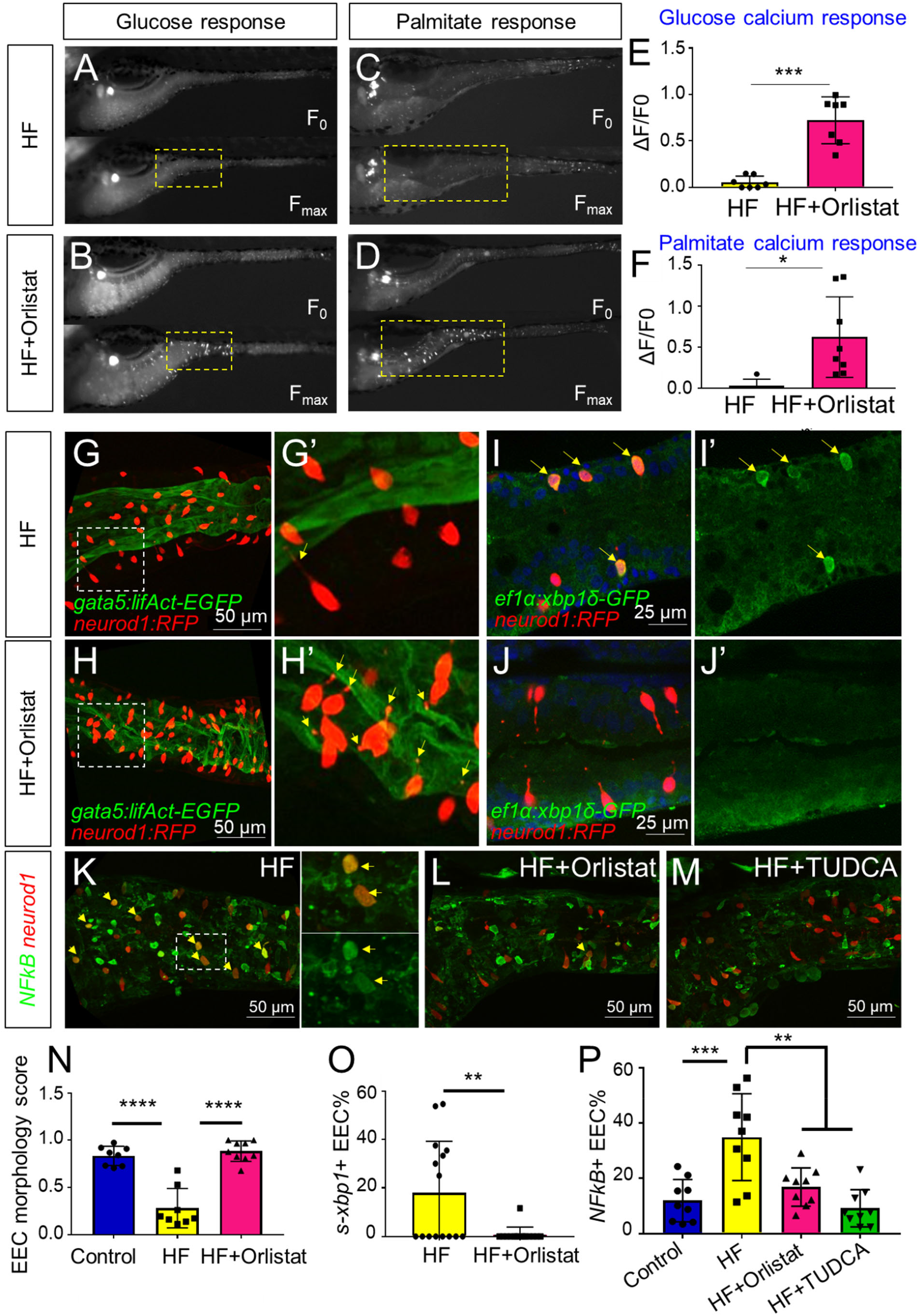
Orlistat treatment inhibits high fat feeding induced EEC silencing. (A-D) Representative image of the EEC calcium response to glucose (A, B) and palmitate (C, D) stimulation in 6 hours high fat fed and 6 hours high fat (HF) fed with 0.5 mM orlistat treated *Tg(neurod1:Gcamp6f)* zebrafish larvae. (E, F) Quantification of the EEC calcium response to glucose and palmitate stimulation in 6 hours HF fed and 6 hours HF fed with 0.5 mM orlistat treated zebrafish larvae. (G-H) Confocal projection of *Tg(neurod1:TagRFP); Tg(gata5:lifActin-EGFP)* zebrafish intestine in 10 hours HF fed larvae (G, G’) and 10 hours HF fed with 0.1 mM orlistat treated larvae (H, H’). The EECs’ apical extensions are marked with yellow arrows. (I-J) Confocal images of *Tg(neurod1:TagRFP); Tg(ef1α:xbp1δ-GFP)* zebrafish intestine in 6 hours HF fed larvae (I, I’) and 6 hours HF fed with 0.5 mM orlistat treated larvae (J, J’). (K-M) Confocal images of *Tg(neurod1:TagRFP); Tg(NFkB:EGFP)* zebrafish intestine in 10 hours HF fed larvae (K), 10 hours HF fed larvae treated with 0.1 mM orlistat (L) and 10 hours HF fed larvae treated with 0.5 mM TUDCA (M). Yellow arrows indicate *neurod1:TagRFP*+ EECs co-labeled with the *NFkB* reporter. (N) Quantification of the EEC morphology score in control, 10 hours HF fed and 10 hours HF fed with 0.1mM orlistat treated larvae represented in G and H. (O) Quantification of *s-xbp1*+ EEC percentage in 6 hours HF fed larvae and 6 hours HF fed larvae treated with 0.5 mM orlistat represented in J and K. (P) Quantification of the percentage of NF-κB+ EECs in control, 10 hour HF fed, 10 hour HF fed with 0.1 mM Orlistat and 10 hour HF fed with 0.5 mM TUDCA zebrafish larvae represented in K-M. Student t-test was performed in E, F, O and one-way ANOVA with post-hoc Tukey test was used in N, P for statistical analysis. * P<0.05, ** P<0.01, *** P<0.001, **** P<0.0001.

To investigate further how orlistat treatment inhibits EEC silencing, we analyzed the its effect on ER stress in EECs following HF feeding using *Tg(ef1α:xbp1δ-gfp)* zebrafish. We found that orlistat treatment significantly reduced the percentage of EECs that are *ef1α:xbp1δ-gfp+* following HF feeding (Fig. 6 I, J, O). We next sought to test if additional pathways are activated in EECs by HF feeding, and if those EEC responses are dependent on lipase activity or ER stress. Induction of ER stress can lead to activation of the transcription factor NF-κB through release of calcium from the ER, elevated reactive oxygen intermediates or direct Ire1 activity (Kim et al., 2015; Pahl & Baeuerle, 1997). After crossing a transgenic reporter of NF-κB activity *Tg(NFkB:EGFP)* (Kanther et al., 2011) with *Tg(-5kbneurod1:TagRFP)*, we found that HF feeding significantly increased the number of NF-κB+ EECs (Fig. 6 K, P), but that this effect could be significantly reduced by treatment with orlistat or the ER stress inhibitor TUDCA (Fig. 6 L, M, P). These results indicate that EEC silencing and associated signaling events that follow ingestion of a HF meal require lipase activity.

Lipases act on dietary triglycerides to liberate fatty acids and monoacylglycerols that are then available for stimulation of EECs (Hara, Hirasawa, Ichimura, Kimura, & Tsujimoto, 2011; Lauffer, Iakoubov, & Brubaker, 2009). To test if free fatty acids are sufficient to induce EEC silencing, we treated 6 dpf zebrafish larvae with palmitate, a major fatty acid component in our HF meal (Poureslami, Raes, Huyghebaert, Batal, & De Smet, 2012). Treatment with palmitate for 6 hours significantly reduced the ability of EECs to respond to subsequent palmitate stimulation, but did not influence the EEC morphology score, nor the EEC response toward subsequent glucose stimulation (Fig. S10). These results suggest that the fatty acid palmitate is sufficient to induce only a portion of the EEC silencing phenotype induced by a complex HF meal.

### High fat feeding induces EEC silencing in a microbiota dependent manner

Using the same HF feeding model in zebrafish, we previously showed that the gut microbiota promote intestinal absorption and metabolism of dietary fatty acids (Semova et al., 2012), and similar roles for microbiota have been established recently in mouse (Martinez-Guryn et al., 2018). We therefore predicted that the microbiota may also regulate EEC silencing after HF feeding. Using our established methods (Pham, Kanther, Semova, & Rawls, 2008), we raised *Tg(gata5:lifActin-EGFP); Tg(-5kbneurod1:TagRFP)* zebrafish larvae to 6 dpf in the absence of any microbes (germ free or GF) or colonized at 3 dpf with a complex zebrafish microbiota (ex-GF conventionalized or CV). In the absence of HF feeding, we observed no significant differences between GF and CV zebrafish in their EEC morphology score or EEC response to palmitate (Fig. 7C,D,G,I). We then performed HF feeding in these 6 dpf GF and CV zebrafish larvae. In contrast to CV HF-fed zebrafish larvae, EECs in GF zebrafish did not show a change in morphology after HF feeding (Fig. 7A, B, I) and exhibited significantly greater responses to palmitate stimulation (Fig. 7E, F, H). In accord, the ability of HF feeding to induce reporters of ER stress and NF-κB activation was significantly reduced in GF compared to CV zebrafish (Fig.7J,K). These results indicate that colonization by microbiota mediates EEC silencing in HF fed zebrafish. EECs are known to express Toll-like receptors (TLRs) (Kanwal, Wiegertjes, Veneman, Meijer, & Spaink, 2014) (Palti, 2011), which sense diverse microbe-associated molecular patterns and signal through the downstream adaptor protein Myd88 leading to activation of NF-κB and other pathways (Kawasaki & Kawai, 2014). To test if EEC silencing requires TLR signaling, we evaluated *myd88* mutant zebrafish (Burns et al., 2017). We found that EECs’ response to palmitate after HF feeding was equivalent to that of wild type fish (Fig. S11 A-B), suggesting microbiota promote EEC silencing in a Myd88-independent manner.

**Figure 7.**
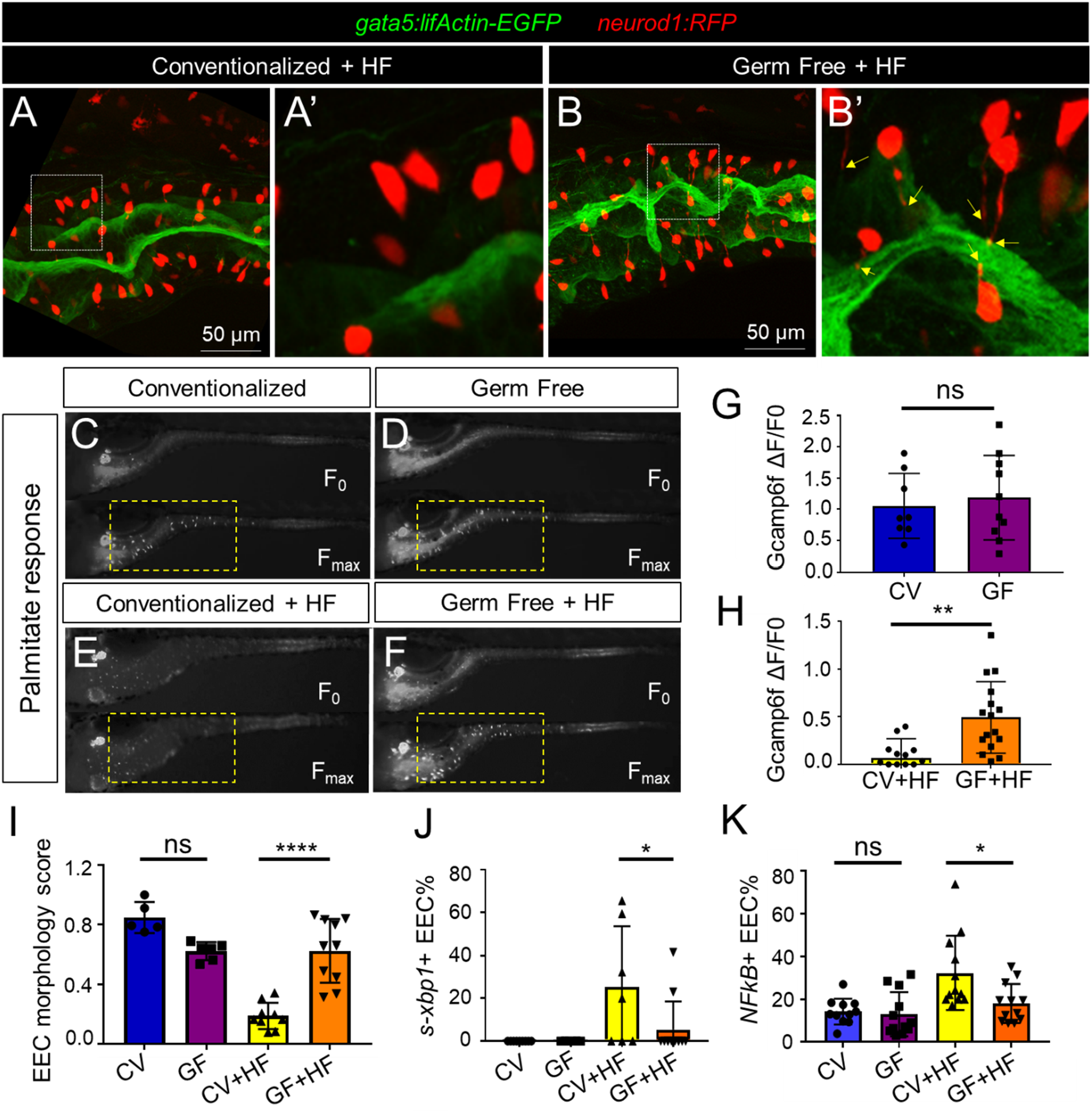
High fat feeding induced EEC silencing is microbiota dependent. (A-B) Confocal images of 6 dpf zebrafish intestines from conventionalized (CV) and germ free (GF) larvae following 10 hours high fat (HF) feeding. EECs are marked with *Tg(neurod1:TagRFP)* and the intestine apical lumen is marked with *Tg(gata5:lifActin-EGFP)*. (C-F) Representative images of the EEC calcium response toward glucose (A, B) and palmitate stimulation in CV and GF *Tg(neurod1:Gcamp6f)* larvae with or without 6 hour HF feeding. (G-H) Quantification of the EEC calcium response to palmitate stimulation represented in C-F. (I) Quantification of the EEC morphology score in CV and GF zebrafish larvae with or without 10 hours HF feeding represented in A and B. (J) Quantification of *xpb1*+ EECs (%) in CV and GF *Tg(neurod1:TagRFP); Tg(ef1α:xbp1δ-GFP)* zebrafish larvae with or without 6 hours HF feeding. (K) Quantification of NF-κB+ EECs (%) in CV and GF *Tg(neurod1:TagRFP); Tg(NFkB:EGFP)* zebrafish larvae with or without 10 hours high fat feeding. Student t-test was used in G,H and one-way ANOVA with post-hoc Tukey test was used in I-K for statistical analysis. * P<0.05, ** P<0.01, **** P<0.0001.

HF diets are known to significantly alter gut microbiota composition in human, mice and zebrafish (David et al., 2014; Hildebrandt et al., 2009; Wong et al., 2015). We therefore hypothesized that HF feeding might alter the composition of the microbiota, which in turn might promote EEC silencing. To test this possibility, we first analyzed the effects of HF feeding on intestinal microbiota density through colony forming unit (CFU) analysis in dissected intestines from CV zebrafish larvae. Strikingly, we found that intestinal microbiota abundance had increased ~20-fold following 6 hours of HF feeding (Fig. 8A). To determine if this increase in bacterial density was accompanied by alterations in bacterial community structure, we performed 16S rRNA gene sequencing. Since diet manipulations can alter microbiota composition in the zebrafish gut as well as their housing water media (Wong et al., 2015), we analyzed samples from dissected intestines of zebrafish larvae in control and HF fed groups as well as their respective housing medias (Fig. 8B). Analysis of bacterial community structure using the Weighted Unifrac method (Caporaso et al., 2010) revealed, as expected, relatively large differences between gut and media samples (PERMANOVA P<0.02 control gut vs. control media, P<0.005 HF gut vs HF media) (Fig. 8C). The addition of HF feeding had a relatively smaller but consistent effect on overall bacterial community structure in both gut and media (PERMANOVA P=0.2 control gut vs HF gut, P=0.094 control media vs HF media) (Fig. 8C). HF feeding caused a small reduction in within-sample diversity among media microbiotas as measured by Faith’s Phylogenetic Diversity (Kruskal-Wallis P=0.049), but no significant effects on gut microbiotas (P=0.29)(Faith & Baker, 2007). Taxonomic analysis of zebrafish gut and media samples revealed several bacterial taxa significantly affected by HF feeding (Table S2). Members of class Betaproteobacteria dominated the control media, but HF feeding markedly decreased their relative abundance (LDA effect size 5.45, P=0.049). Conversely, HF feeding increased the relative abundance of members of class Gammaproteobacteria (LDA effect size 5.49, P=0.049; Fig.8D) such as genera *Acinetobacter* (LDA effect size 5.13, P=0.049), *Pseudomonas* (LDA effect size 5.02, P=0.049) and *Aeromonas* (LDA effect size 4.78, P=0.049; Fig. 8E; Tables S2 and S3). HF feeding also increased the relative abundance in media of class Cytophagia from phylum Bacteroidetes (LDA effect size 4.66, P=0.049; Fig. 8D) due to increases in the genus *Flectobacillus* (LDA effect size 4.76, P=0.049; Fig. 8F; Tables S2 and S3). The increased relative abundances of *Aeromonas* sp. and *Pseudomonas* sp. in HF fed medias was not recapitulated in the gut microbiotas (Fig.8G; Table S2). However, similar to the media, HF feeding significantly increased abundance of class Cytophagia (LDA effect size 4.01, P=0.018; Fig.8D) due to enrichment of *Flectobacillus* (LDA effect size 4.01, P=0.004; Fig.8H). Additionally, HF feeding resulted in a 100-fold increase the relative abundance of *Acinetobacter* sp. in the gut (average 0.04% in control gut, 4.28% in HF gut; LDA effect size 4.31, P=0.001; Fig. 8G, Tables S2 and S4). These results establish that HF feeding has diverse effects on the bacterial communities in the zebrafish gut and media, and raise the possibility that members of these affected bacterial genera may regulate EEC silencing in response to HF feeding.

**Figure 8.**
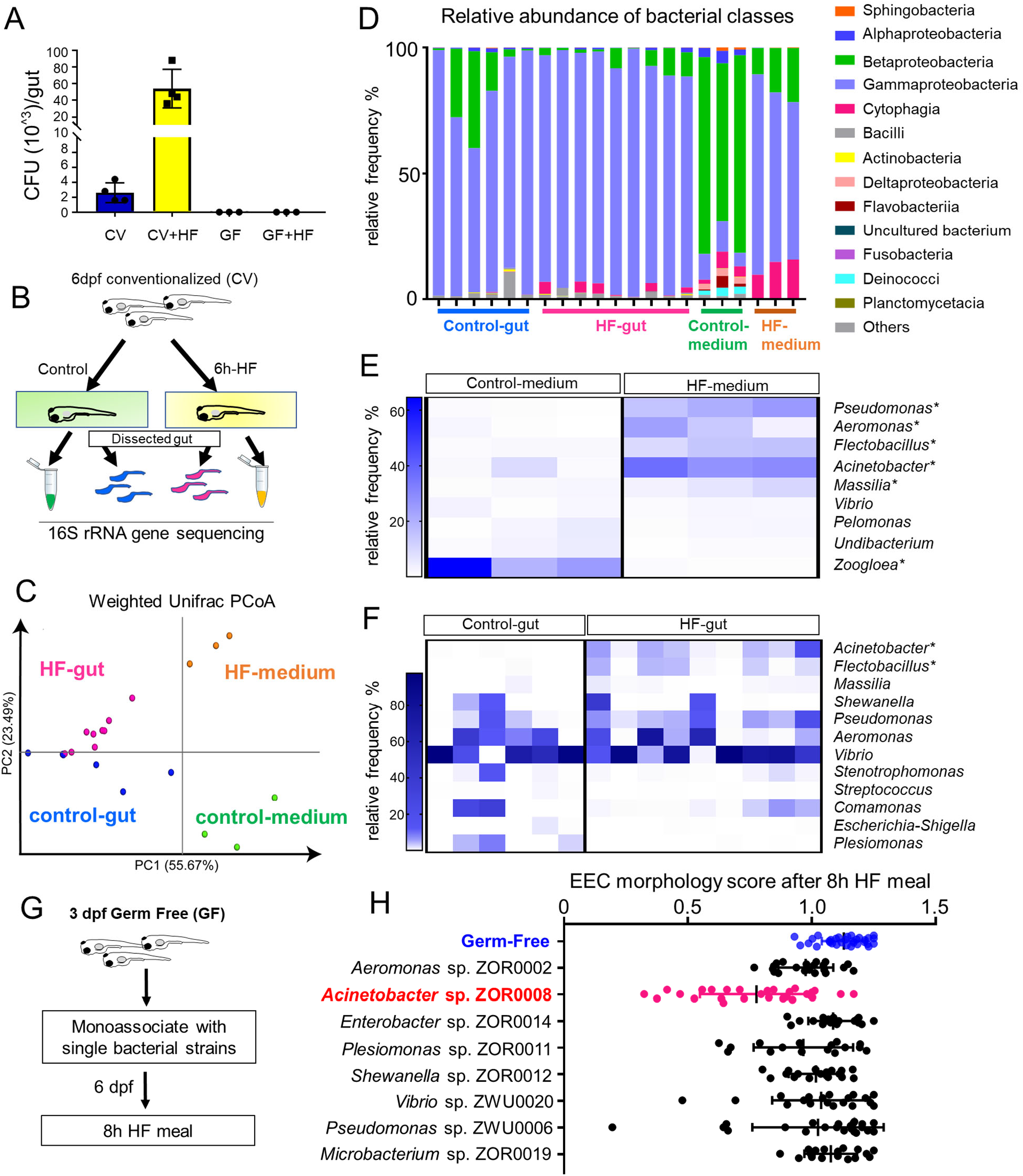
High fat feeding modified microbiota composition. (A) CFU quantification in GF and CV dissected intestine with or without 6 hours high fat (HF) feeding. (B) Experimental design of 16S rRNA gene sequencing in control larvae dissected gut and medium and 6 hour HF fed larvae dissected gut and medium. (C) PCoA plot representing the microbiome community in control and HF fed gut and media. (D) Relative abundance of bacteria composition at the class level in control and HF fed gut and medium. (E-F) Change in representative specific bacteria genus following HF feeding in gut and medium. * indicates taxa with P<0.05 by LEfSe analysis. (G) Schematic of mono-association screening to investigate the effects of specific bacteria strains on EEC morphology. 3 dpf zebrafish larvae were colonized with one of the isolated bacteria strains and EEC morphology was scored after 8-hour high fat meal feeding in 6 dpf GF and mono-associated animals. (H) EEC morphology score of GF and monoassociated zebrafish larvae following 8 hour high fat feeding. Data were pooled from 3 independent experiments, with each dot representing an individual animal. The EEC morphology score in *Acinetobacter* sp. ZOR0008 mono-associated animals was significantly lower than GF EECs (P<0.001). No significant difference was observed in other bacteria strain monoassociated group. One way ANOVA followed by Tukey’s post-test was used in H for statistical analysis.

We next tested if EEC silencing could be facilitated by representative members of the zebrafish microbiota, including those enriched by HF feeding. We selected a small panel of bacterial strains that were isolated previously from the zebrafish intestine (Stephens et al., 2016) and used them to monoassociate separate cohorts of GF *Tg(gata5:lifActin-EGFP)*; *Tg(-5kbneurod1:TagRFP)* zebrafish at 3dpf (Fig. 8I). These bacteria strains were from 9 different genera including *Acinetobacter* sp. ZOR0008, a member of the *Acinetobacter calcoaceticus*-*Acinetobacter baumannii* complex (Gerner-Smidt, Tjernberg, & Ursing, 1991) (Bouvet & Jeanjean, 1989). We did not observe significant differences in colonization efficiency among these bacteria strains that were inoculated into GF zebrafish (Fig. S12A-B). At 6 dpf, we performed HF feeding and examined the EEC morphology score. Strikingly, only *Acinetobacter* sp. ZOR0008 was sufficient to significantly reduce the EEC morphology score upon HF feeding (Fig. 8J) similar to conventionalized animals (Fig. 7A,B,I). These results indicate that the effects of microbiota on EEC silencing following HF feeding display strong bacterial species specificity, and suggest *Acinetobacter* bacteria enriched by HF feeding may mediate the effect of microbiota on HF sensing by EECs.

## Discussion

In this study, we established a new experimental system to directly investigate EEC activity *in vivo* using a zebrafish reporter of EEC calcium signaling. Combining genetics, diet and gnotobiotic manipulations allowed us to uncover a novel EEC adaptation mechanism through which high fat feeding induces rapid change of EEC morphology and reduced nutrient sensitivity. We called this novel adaptation “EEC silencing”. Our results show that EEC silencing following a high fat meal requires lipase activity and it is coupled to ER stress. Furthermore, HF meal induced EEC silencing is promoted by certain microbial species (e.g., *Acinetobacter* sp.). As discussed below, we propose a working model (Fig. S13) that nutrient over-stimulation from high fat feeding increases EECs hormone synthesis burden, overgrowth of the gut bacterial community including enrichment of Acinetobacter sp., which in turn activates EECs ER stress response pathways and thereby induces EECs silencing. This study demonstrates the utility of the zebrafish model to study *in vivo* interactions between diet, gut microbes, and EEC physiology. In the future, the mechanisms underlying EEC silencing could be targeted in rational manipulations of EEC adaptations to diet and microbiota which could be used to reduce incidence and severity of metabolic diseases.

### EEC physiology in zebrafish

Our studies here provide important new tools for studying EECs in the context of zebrafish intestinal epithelial development and physiology. Similar to mammals, fish EECs are thought to arise from intestinal stem cells through a series of signals that govern the differentiation process (Aghaallaei et al., 2016). Delta-Notch signaling appears to control the differentiation of stem cells into absorptive and secretory cell lineages in both zebrafish and mammalian models (Crosnier et al., 2005). Activation of Notch signaling can block the differentiation of EECs by inhibiting the expression of key EEC bHLH transcription factors (H. J. Li et al., 2011). In mammals, the bHLH transcription factor Neurod1 that has been shown to regulate EEC terminal differentiation (H. J. Li et al., 2011; Ray & Leiter, 2007). Our results indicate that Neurod1 is expressed by and important in EEC differentiation in zebrafish as it is in mammals. Moreover, this finding enabled us to use *neurod1* regulatory sequences to label and monitor zebrafish EECs.

The hallmark of EECs is their expression of hormones. In this study, using transgenic reporter lines and immunofluorescence staining approaches to examine a panel of gut hormones in zebrafish EECs, we found that zebrafish EECs express conserved hormones as mammalian EECs. Interestingly, a subset of EECs express proglucagon peptide which can be processed to hormones Glucagon like peptide 1 (GLP-1) and 2 (GLP-2) (Sandoval & D’Alessio, 2015). GLP-1, one of the incretin hormones, is released by EECs in response to oral glucose intake that can faclilate insulin secretion and reduce blood glucose (Drucker, Habener, & Holst, 2017). Multiple studies suggest that the expression of Sglt1 is important for EEC glucose sensing (Gorboulev et al., 2012; Reimann et al., 2008; Roder et al., 2014). EECs in Sglt1 knockout mice fail to secrete GLP-1 in response to glucose and galactose (Gorboulev et al., 2012). In our studies, we identified similar Sglt1 mediated glucose sensing machinery in zebrafish EECs. This together suggest that zebrafish EECs may exhibit conserved roles in regulating glucose metabolism.

Our data also establish that zebrafish EECs develop striking regional specificity in the hormones they express along the intestine (Fig. S1). For example, the CCK and PYY hormones that are important for regulating food digestion and energy homeostasis (Beglinger & Degen, 2006; Liddle, 1997; Raybould, 2007) were only expressed in the proximal intestine. In addition to hormonal regional specificity, we found that EEC calcium response toward nutrients also display regional specificity. For example, glucose and long chain/medium chain fatty acids only stimulate EECs in proximal intestine, a region in zebrafish where digestion and absorption of dietary fats primarily occurs (Carten et al., 2011). This hormonal and functional regional specificity suggests that distinct developmental and physiological programs govern EEC function along the intestinal tract, and that EECs in the proximal zebrafish intestine may play key roles in monitoring and adapting to dietary nutrient experience.

### EEC silencing

In this study, we discovered that high fat feeding can induce a series of functional and morphological changes in EECs we refer to as “EEC silencing”. EEC silencing includes (1) reduced EEC sensitivity to nutrient stimulation (e.g., fatty acids and glucose) and (2) conversion of EEC morphology from an open to a closed type. To our knowledge, EEC silencing has not been observed in previous studies of EEC in any vertebrate. This underscores the unique power of *in vivo* imaging in zebrafish to reveal new physiologic and metabolic processes. Our evidence suggests that EEC silencing is a stress response that EECs display following consumption of a high fat meal in the presence of specific microbes. However, EEC silencing may also serve to protect EECs against excessive stress following consumption of a high fat meal. In neurons for example, similar desensitization has been shown to protect nerve cells from excitatory neurotransmitter induced toxicity (Gainetdinov, Premont, Bohn, Lefkowitz, & Caron, 2004; Quick & Lester, 2002) and blocking desensitization of excitatory neuronal receptors induces rapid neuronal cell death (Walker et al., 2009). High dietary fat can also lead to excessive production of excitatory stimuli like long-chain fatty acids. We speculate that EEC silencing provides an adaptive mechanism for EECs to avoid excessive stimuli and protect against cell death.

The observation that EECs exhibited reduced sensitivity to oral glucose following high fat feeding is interesting and consistent with the finding in mice that high fat feeding reduces intestinal glucose sensing and glucose induces GLP-1 secretion *in vivo* (Bauer et al., 2018). *In vitr*o, small intestinal cultures from high fat fed mice also exhibit reduced secretory responsiveness to nutrient stimuli including glucose when comparing with intestine cultures from control mice (Richards et al., 2016) but underlying mechanisms remained unclear. These studies, together with our results, suggest that high fat feeding impairs EEC function. However, how high fat feeding reduces EEC glucose sensitivity is still unclear as we did not detect changes in the EEC glucose sensor *sglt1* expression in high fat fed dissected intestine (data not shown). One possibility is that high fat feeding affects EECs glucose sensing via altering Sglt1 activity (Ishikawa, Eguchi, & Ishida, 1997; Subramanian, Glitz, Kipp, Kinne, & Castaneda, 2009; Wright, Hirsch, Loo, & Zampighi, 1997). We also speculate that high fat feeding induced EEC morphological changes may contribute to EEC glucose insensitivity. Since Sglt1 is expressed on the brush border at the apical surface of the cell, as EECs switch from an open to closed type morphology they would lose their contact with the gut lumen and exposure to luminal glucose stimuli.

Our observation that EECs can change their morphology from an “open” to “closed” state upon high fat feeding was surprising. The majority of EECs in the intestinal tract are open with an apical extension and microvilli facing the intestinal lumen. In contrast, some EECs lie flat on the basement membrane and are “closed” to the gut lumen (Gribble & Reimann, 2016). The presence of open and closed EECs has been observed in both mammals and fish (Rombout, Lamers, & Hanstede, 1978). Previously, it was believed that the open and closed EECs were two differentiated EEC types that perhaps had different physiological functions (Gribble & Reimann, 2016). The open EECs were thought to sense and respond to luminal stimulation while although less clear the closed EECs were thought to respond to hormonal and neuronal stimulation from the basolateral side. However, our data reveal for the first time that EECs can convert from an open to a closed state. This indicates that EECs possess plasticity to actively prune their apical extension. The pruning of cellular process can be observed extensively in neurons. Studies from multiple organisms revealed that sensory neurons can eliminate their dendrites and axon during development and in response to injury through active pruning (Kanamori et al., 2013; Nikolaev, McLaughlin, O’Leary, & Tessier-Lavigne, 2009; Sagasti, Guido, Raible, & Schier, 2005; Williams, Kondo, Krzyzanowska, Hiromi, & Truman, 2006; Yu & Schuldiner, 2014). This process includes focal disruption of the microtubule cytoskeleton, followed by thinning of the disrupted region, severing and fragmentation and retraction in proximal stumps after severing events (Williams & Truman, 2005). In our system, the thinning and fragmentation in the EEC apical extension was also observed. It is well known that EECs possess many neuron-like features including neurotransmitters, neurofilaments, and synaptic proteins (Bohorquez et al., 2015). Whether EECs adopt the same mechanisms as neurons to prune their cellular processes in response to nutritional and microbial signals is interesting and requires future study.

### The effects of diet and microbes on EEC silencing

In this study, we have shown that both diet and microbes play important roles in inducing EEC silencing. Dietary manipulations and changes in gut microbiota have been shown to affect EEC cell number and GI hormone gene expression in mice and zebrafish (Arora et al., 2018; Rawls, Samuel, & Gordon, 2004; Richards et al., 2016; Troll et al., 2018). However, it remains unclear from previous studies how environmental factors like diet and gut microbiota affect EEC function. We found that while the presence of microbiota did not influence EEC nutrient sensing under basal conditions, microbiota played an essential role in mediating high fat diet induced EEC silencing as germ free EECs were resistant to high fat diet induced silencing. We speculate that EEC silencing may temporarily attenuate the host’s ability to accurately sense ingested nutrients and thereby control energy homeostasis. Our finding that gut microbiota play an essential role in high fat diet induced EEC silencing may provide a new mechanistic inroad for understanding the effects of gut microbiota in diet induced metabolic diseases including obesity and insulin resistance (Backhed, Manchester, Semenkovich, & Gordon, 2007; Rabot et al., 2010).

There are several nonexclusive ways by which specific gut microbiota members such as *Acinetobacter* sp. might affect EECs in the setting of a high fat diet. First, microbiota could affect EEC development to increase production of EEC subtypes that are relatively susceptible to diet-induced EEC silencing. Previous transcriptome analysis in the ileum of GLP-1 secreting EECs showed that microbiota colonization increased transcript levels of genes associated with synaptic cycling, ER stress response and cell polarity was reduced in germ free mice (Arora et al., 2018). This suggests that EECs in colonized animals may be more prone to diet-induced ER stress and morphological changes including those associated with EEC silencing.

Second, high fat meal conditions induce bacterial overgrowth and alter the selective pressures within the gut microbial community to allow for enrichment and depletion of specific bacterial taxa. Such changes in microbial density and community composition may then acutely affect EEC physiology. Indeed, we found that high fat feeding altered the relative abundance of several bacterial taxa in the zebrafish gut and media, including a 100-fold increase for members of the *Acinetobacter* genus. Strikingly, a representative *Acinetobacter* sp. was the only strain we identified that was sufficient to mediate high fat induced alterations in EEC morphology. We speculated that bacterial overgrowth may also result in increased presentation of microbe-associated molecular patterns which could then hyper-activate TLR or other microbe-sensing pathways that could lead to EEC functional changes. However, our data from *myd88* mutant zebrafish suggest that at least the Myd88-dependent microbial sensing pathways are not required for high fat induced EEC silencing. As described below, identification of the specific signals produced by *Acinetobacter* sp. and other bacteria that facilitate EEC silencing remain an important research goal.

Third, gut microbiota might affect EEC function through promoting lipid digestion and absorption. This is supported by our observations that blocking fat digestion and subsequent lipid absorption in enterocytes through orlistat treatment inhibited high fat diet induced EEC silencing. EEC function may be directly influenced by the products of lipolysis such as free fatty acids (Edfalk, Steneberg, & Edlund, 2008; Hirasawa et al., 2005; Katsuma et al., 2005). However, in our experiments, palmitate treatment was only sufficient to reproduce a portion of the EEC silencing phenotype (i.e. loss of nutrient sensitivity), suggesting that additional undefined signals from fat digestion in the intestine are required to fully induce EEC silencing. In the intestinal epithelium, EECs are surrounded by enterocytes and these two cell types exhibit complex bi-directional communication (Hein, Baker, Hsieh, Farr, & Adeli, 2013; Hsieh et al., 2009; Okawa et al., 2009; Shimotoyodome et al., 2009). Following ingestion of a complex high fat meal, free fatty acids and glycerol liberated from triglyceride digestion are taken up by enterocytes and assembled into lipid droplets and chylomicrons (Phan & Tso, 2001). The subsequent enlargement of enterocytes from lipid droplet accumulation may exert mechanical pressure on EECs that then induces EECs adaption through pruning of their apical protrusions. Besides mechanical pressure, lipoproteins and free fatty acids released from enterocytes may act on EECs basolaterally to alter their function (Chandra et al., 2013; Okawa et al., 2009; Shimotoyodome et al., 2009). Previous studies have shown that lipid digestion and absorption is impaired in germ free animals and enterocytes in germ free condition exhibit reduced lipid droplet accumulation (Martinez-Guryn et al., 2018; Semova et al., 2012). Therefore, reduced mechanical pressure or secondary signaling molecules from enterocytes in the germ free condition may lead to the resistance of EECs to high fat induced silencing. On the other hand, gut microbiota may promote EECs silencing by facilitating intestinal lipid digestion and absorption. *Acinetobacter* was the most highly enriched genus in the larval zebrafish intestine following high fat feeding in this study and was also enriched in adult zebrafish gut following a chronic high fat diet (Wong et al., 2015). Further, we identified a representative member of this genus that is sufficient to mediate EEC silencing under high fat diet conditions. However, the molecular mechanisms by which *Acinetobacter* spp. evoke this host response remains unknown. Studies suggest that *A. baumannii*, a closely related oportunitistic pathogen, can signal to host epithelial cells through secreted outer membrane vesicles (OMVs) and activation of downstream inflammatory pathways (Jha, Ghosh, Gautam, Malhotra, & Ray, 2017; Jin et al., 2011; Jun et al., 2013; March et al., 2010). In addition to OMVs, *Acinetobacter* strains are known to secrete phospholipase that can affect host cell membrane stability and interfere with host signaling (Lee et al., 2017; Songer, 1997). Members of the *Acinetobacter* genus are also known to possess potent oil degrading and lipolytic activities (Lal & Khanna, 1996; Snellman & Colwell, 2004). Moreover, species from *Acinetobacter* genus have the ability to produce emulsifiers which might enhance lipid digestion (Navon-Venezia et al., 1995; Toren, Navon-Venezia, Ron, & Rosenberg, 2001; Walzer, Rosenberg, & Ron, 2006). Interestingly, *Acinetobacter* spp. in the human gut are positively associated with plasma TG and total- and LDL-cholesterol (Graessler et al., 2013), and *Acinetobacter* spp. are also enriched in the crypts of the small intestine and colon in mammals (Mao, Zhang, Liu, & Zhu, 2015; Pedron et al., 2012; Saffarian et al., 2017). Therefore, it will be intertesting to determine whether *Acinetobacter* spp. also modulate EEC function in mammals under high fat diet conditions. Finally, considering the small scale of our monoassocation screen, we anticipate that additional members of the gut microbiota in zebrafish and other animals will be found to also affect EEC silencing and other aspects of EEC biology.

## MATERIALS AND METHODS

### Zebrafish strains and husbandry

All zebrafish experiments conformed to the US Public Health Service Policy on Humane Care and Use of Laboratory Animals, using protocol number A115-16-05 approved by the Institutional Animal Care and Use Committee of Duke University. Conventionally-reared adult zebrafish were reared and maintained on a recirculating aquaculture system using established methods (Murdoch et al., 2019). For experiments involving conventionally-raised zebrafish larvae, adults were bred naturally in system water and fertilized eggs were transferred to 100mm petri dishes containing ~25mL of egg water at approximately 6 hours post-fertilization. The resulting larvae were raised under a 14h light/10h dark cycle in an air incubator at 28°C at a density of 2 larvae/ml water. To ensure consistent microbiota colonization, 10mL filtered system water (5μm filter, SLSV025LS, Millipore) was added into 3 dpf zebrafish larva that were raised in 25mL egg water. All the experiments performed in this study ended at 6 dpf unless specifically indicated. The strains used in this study are listed in Table S1. All lines were maintained on a EKW background.

Gateway Tol2 cloning approach was used to generate *neurod1:lifActin-EGFP* and *neurod1:Gal4* plasmid (Kawakami, 2007; Kwan et al., 2007). The 5kb pDONR-neurod1 P5E promoter was previously reported (McGraw et al., 2012) and generously donated by Dr. Hillary McGraw. The PME-lifActin-EGFP (Riedl et al., 2008) and the PME-Gal4-vp16 plasmids (Kwan et al., 2007) were also previously reported. pDONR-neurod1 P5E and PME-lifActin-EGFP was cloned into pDestTol2pA2 through an LR Clonase reaction (ThermoFisher,11791). Similarly, pDONR-neurod1 P5E and PME-Gal4-vp16 was cloned into pDestTol2CG2 containing a *cmlc2:EGFP* marker. The final plasmid was sequenced and injected into the wild-type Ekkwill (EKW) zebrafish strain and the F2 generation of allele *Tg(neurod1:lifActin-EGFP)^rdu70^* and *Tg(neurod1:Gal4; cmlcl2:EGFP)^rdu71^* was used for this study.

The construct used to generate the *TgBAC(gata5:lifActin-EGFP)* line was made by inserting lifeact-GFP at the *gata5* ATG in the BAC clone DKEYP-73A2 using BAC recombineering as previously described (PMID: 12618378). The BAC was then linearized using I-SceI and injected to generate transgenic lines. Allele *TgBAC(gata5:lifActin-EGFP)^pd1007^* was selected for further analysis. The construct used to generate the *TgBAC(cd36-RFP)* lines was made by inserting link-RFP before the *cd36* stop codon in the BAC clone DKEY-27K7 using the same BAC recombineering as previously described (Navis et al PMID 23487313). Then, Tol2 sites were recombined into the BAC and the resulting construct was injected with transposase mRNA (cite Kawakami here) to generate the transgenic lines. Allele *TgBAC(cd36-RFP)^pd1203^* was selected for further analysis.

### Gnotobiotic zebrafish husbandry

For experiments involving gnotobiotic zebrafish, we used our established methods to generate germ-free zebrafish using natural breeding (Pham et al., 2008) with the following exception: Gnotobiotic Zebrafish Medium (GZM) with antibiotics (AB-GZM) was supplemented with 50 μg/ml gentamycin (Sigma, G1264). Germ free zebrafish eggs were maintained in cell culture flasks with GZM at a density of 1 larvae/ml. From 3 dpf to 6 dpf, 60% daily media change and ZM000 (ZM Ltd.) feeding were performed as described (Pham et al., 2008).

To generate conventionalized zebrafish, 15 mL filtered system water (5μm filter, SLSV025LS, Millipore, final concentration of system water ~30%) was inoculated to flasks containing germ-free zebrafish in GZM at 3 dpf when the zebrafish normally hatch from their protective chorions. The same feeding and media change protocol was followed as for germ free zebrafish. Microbial colonization density was determined via Colony Forming Unit (CFU) analysis. To analyze the effect of high fat feeding on intestinal bacteria colonization, dissected digestive tracts were dissected and pooled (5 guts/pool) into 1mL sterile phosphate buffered saline (PBS) which was then mechanically disassociated using a Tissue-Tearor (BioSpec Products, 985370). 100 µL of serially diluted solution was then spotted on a Tryptic soy agar (TSA) plate and cultured overnight at 30°C under aerobic conditions.

To generate mono-associated zebrafish, a single bacterial strain was inoculated into each flask containing 3dpf germ-free zebrafish. The respective bacterial stock was streaked on a TSA plate and cultured at 28°C overnight under aerobic conditions. A single colony was picked and cultured in 5mL Tryptic soy broth media shaking at 30°C for 16 hours under aerobic conditions. 250 µL bacterial culture was pelleted and washed 3 times with sterile GZM and inoculated into flasks containing germ-free zebrafish. OD600 and CFU measurements were performed in each mono-associated culture. The final innoculation density in GZM was 10^8^-10^9^ CFU/mL. The colonization efficiency was determined at 6 dpf by CFU analysis from dissected zebrafish intestines as described above.

### EEC response assay and image analysis

This assay was performed in *Tg(neurod1:Gcamp6f)* 6 dpf zebrafish larvae. Unanesthetized zebrafish larvae were gently moved into 35mm petri dishes that contained 500µL 3% methylcellulose. Excess water was removed with a 200µL pipettor. Zebrafish larvae were gently positioned horizontal to the bottom of the petri dish right side up carefully avoiding touching the abdominal region and moved onto an upright fluorescence microscope (Leica M205 FA microscope equipped with a Leica DFC 365FX camera). The zebrafish larvae were allowed to recover in that position for 2 minutes. One hundred µL of test agent was pipetted directly in front of the mouth region without making direct contact with the animal. Images were recorded every 10 seconds. For fatty acid stimulation, 30 frames (5mins) were recorded. For glucose stimulation, 60 frames (10mins) were recorded. The *Gcamp6f* fluorescence was recorded with the EGFP filter. The following stimulants were used in this study: palmitic acid/linoleate/dodecanoate (1.6mM), butyrate (2mM), glucose (500mM), fructose (500mM), galactose (500mM), cysteine (10mM). Since palmitic acid/linoleate/dodecanoate was not water soluble by itself, 1.6% BSA was used as a carrier to facilitate solubility. Solutions were filtered with 0.22µm filter.

Image processing and analysis was performed using FIJI software. The time-lapse fluorescent images of zebrafish EEC response to nutrient stimulation were first aligned to correct for experimental drift using the plugin “align slices in stack.” Normalized correlation coefficient matching method and bilinear interpolation method for subpixel translation was used for aligning slices (Tseng et al., 2012). The plugin “rolling ball background subtraction” with the rolling ball radius=10 pixels was used to remove the large spatial variation of background intensities. The Gcamp6f fluorescence intensity in the proximal intestinal region was then calculated for each time point. The ratio of maximum fluorescence (F_max_) and the initial fluorescence (F_0_) was used to measure EEC calcium responsiveness.

### High fat feeding

The high fat feeding regimen was performed in 6 dpf zebrafish larvae using methods previously described (Semova et al., 2012). ~25 zebrafish larvae were transferred into 6 well plates and 5mL egg water (for gnotobiotic studies, GZM was used). Replicates were performed in three wells for each treatment group in each experiment. Chicken eggs were obtained from a local grocery store from which 1mL chicken egg yolk was transferred into a 50mL tube containing 15mL egg water (for gnotobiotic studies, sterile GZM was used). Solutions were sonicated (Branson Sonifier, output control 5, Duty cycle 50%) to form a 6.25% egg yolk emulsion. 4 mL water from each well was removed and replenished with 4mL egg yolk. 4 mL egg water emulsion was used to replenish the control group. Zebrafish larvae were incubated at 28°C for the indicated time. The high fat meal was administered between 10am - 12pm to minimize circadian influences.

### Chemical treatment

To block Sglt1, phloridzin (0.15mM, Sigma P3449) was used to pretreat zebrafish for 3 hours prior to glucose stimulation, and 0.15mM phloridzin was co-administered with the glucose stimulant solution. To induce ER stress, thapsigargin (0.75µM, Sigma T9033) and brefeldin A (9µM, Sigma B6542) were added to egg water and zebrafish were treated for 10 hours prior to performing the EEC activity assay. To block high fat meal induced EEC silencing, sodium tauroursodeoxycholic acid (TUDCA; 0.5mM, T0266) or orlistat (0.1mM, Sigma O4139) were added to the high fat meal solution and zebrafish were treated for the indicated time.

### Quantitative RT-PCR

The quantitative real-time PCR was performed as described previously (Murdoch et al., 2019). In brief, 20 zebrafish larvae digestive tracts were dissected and pooled into 1mL TRIzol (ThermoFisher, 15596026). mRNA was then isolated with isopropanol precipitation and washed with 70% EtOH. 500ng mRNA was used for cDNA synthesis using the iScript kit (Bio-Rad, 1708891). Quantitative PCR was performed in triplicate 25 μl reactions using 2X SYBR Green SuperMix (PerfeCTa, Hi Rox, Quanta Biosciences, 95055) run on an ABI Step One Plus qPCR instrument using gene specific primers (Table S1). Data were analyzed with the ΔΔCt method. 18S was used as a housekeeping gene to normalize gene expression.

### 16S rRNA gene sequencing

Wild-type adult EKW zebrafish were bred and clutches of eggs from three distinct breeding pairs were collected, pooled, derived into GF conditions using our standard protocol (Pham et al., 2008), then split into three replicate flasks with 30ml GZM as described above. At 3 dpf 12.5ml 5μm-filtered system water was inoculated into each flask per our standard conventionalization method. ZM000 feeding and water changes were performed daily from 4 dpf to 5 dpf. At 6 dpf, zebrafish larvae from each flask were divided evenly into a control and a high fat fed group. High fat feeding was performed as described above for 6 hours. Then 1 ml water samples were collected from each flask and snap frozen on dry ice/EtOH bath. For intestinal samples, individual digestive tracts from 6dpf zebrafish were dissected and flash frozen (3-4 larvae/flask, 3 flasks/condition). All samples were stored in −80°C for subsequence DNA extraction.

The Duke Microbiome Shared Resource (MSR) extracted bacterial DNA from gut and water samples using a MagAttract PowerSoil DNA EP Kit (Qiagen, 27100-4-EP) as described previously (Murdoch et al., 2019). Sample DNA concentration was assessed using a Qubit dsDNA HS assay kit (ThermoFisher, Q32854) and a PerkinElmer Victor plate reader. Bacterial community composition in isolated DNA samples was characterized by amplification of the V4 variable region of the 16S rRNA gene by polymerase chain reaction using the forward primer 515 and reverse primer 806 following the Earth Microbiome Project protocol (http://www.earthmicrobiome.org/). These primers (515F and 806R) carry unique barcodes that allow for multiplexed sequencing. Equimolar 16S rRNA PCR products from all samples were quantified and pooled prior to sequencing. Sequencing was performed by the Duke Sequencing and Genomic Technologies shared resource on an Illumina MiSeq instrument configured for 150 base-pair paired-end sequencing runs. Sequence data are deposited at SRA under Bioproject accession number PRJNA532723.

Subsequent data analysis was conducted in QIIME2 (Caporaso et al., 2010)(https://peerj.com/preprints/27295/). Paired reads were demultiplexed with qiime demux emp-paired, and denoised with qiime dada2 denoise-paired (Callahan et al., 2016). Taxonomy was assigned with qiime feature-classifier classify-sklearn (Scikit-learn: Machine Learning in Python. Journal of Machine Learning Research), using a naive Bayesian classifier, trained against the 99% clustered 16S reference sequence set of SILVA, v. 1.19 (Quast et al., 2013). A basic statistical diversity analysis was performed, using qiime diversity core-metrics-phylogenetic, including alpha- and beta-diversity, as well as relative taxa abundances in sample groups. The determined relative taxa abundances were further analyzed with LEfSe (Linear discriminant analysis effect size) (Segata et al., 2011), to identify differential biomarkers in sample groups.

### Immunofluorescence Staining and Imaging

The whole mount immunofluorescence staining was performed as previously described (Ye et al., 2015). In brief, ice cold 2.5% formalin was used to fix zebrafish larvae overnight at 4°C. The samples were then washed with PT solution (PBS+0.75%Triton-100). The skin and remaining yolk was then removed using forceps under a dissecting microscope. The deyolked samples were then permeabilized with methanol for more than 2 hours at −20°C. The samples were then blocked with 4% BSA at room temperature for more than 1 hour. The primary antibody was diluted in PT solution and incubated at 4°C for more than 24 hours. Following primary antibody incubation, the samples were washed with PT solution and incubated overnight with secondary antibody with Hoechst 33342 for DNA staining. The imaging process was performed with a Zeiss 780 inverted confocal and Zeiss 710 inverted confocal microscopes with the 40× oil lenses. The following primary antibodies were used in this study: rabbit anti PYY (custom, aa4-21, 1:100 dilution) (Chandra, Hiniker, Kuo, Nussbaum, & Liddle, 2017), goat anti-CCK (Santa Cruz SC-21617, 1:100 dilution), rabbit anti-Sglt1 (Abcam ab14686, 1:100 dilution). The secondary antibodies used in this study are from Alexa Fluor Invitrogen. All the secondary antibodies were used at a dilution of 1:250.

To quantify EEC morphology score, chick anti-GFP (Aves GFP1010, 1:500 dilution) and rabbit anti-mcherry (TAKARA 632496, 1:250 dilution) antibodies were used in the fixed *Tg(gata5:lifActin-EGFP)*;*Tg(neurod1:TagRFP)* samples to perform immunofluorescence staining. The region following intestine bulb were imaged with a Zeiss 780 inverted confocal and Zeiss 710 inverted confocal microscopes with the 40× oil lenses. Images were processed with FIJI. The *gata5:lifActin-EGFP* only stains the apical brush border of the intestine. Total EECs number was assessed via counting RFP+ cell bodies. The number of EECs with intact apical protrusion was assessed via counting the number of RFP+ cells with attachment to GFP staining brush border. EEC morphology for each sample were quantified as ratio between EECs with intact apical protrusion and total EEC number.

For live imaging experiments, zebrafish larvae were anesthetized with Tricane and mounted in 1% low melting agarose in 35mm petri dishes. The live imaging was recorded with Zeiss 780 upright confocal with a 20× water lens.

### Statistical Analyses

The appropriate sample size for each experiment was suggested by preliminary experiments evaluating variance and effects. Using significance level of 0.05 and power of 80%, a biological replicate sample number 10 was suggested for EEC calcium response analysis and a biological replicate sample number 13 was suggested for EEC morphology analysis. For each experiment, wildtype or indicated transgenic zebrafish embryos were randomly allocated to test groups prior to treatment. In some EEC calcium response experiments, less than 10 biological replicate samples were used due to technical limitations associated with live sample imaging. In EEC morphology analysis, each experiment contained 8-15 biological replicates or individual fish samples. Individual data points, mean and standard deviation are plotted in each figure.

The raw data points in each figure are represented as solid dots. The data was analyzed using GraphPad Prism 7 software. For experiments comparing just two differentially treated populations, a Student’s t-test with equal variance assumptions was used. For experiments measuring a single variable with multiple treatment groups, a single factor ANOVA with post hoc means testing (Tukey) was utilized. Statistical evaluation for each figure was marked * P<0.05, ** P<0.01, *** P<0.001, **** P<0.0001 or ns (no significant difference, P>0.05). Statistical analyses for 16S rRNA gene sequencing data can be found in in the corresponding Methods section above.

## Acknowledgements

We thank Dr. Hillary McGraw for the 5kb pDONR-neurod1 P5E plasmid, Dr. Joachim Berger for the pMElifActin-EGFP plasmid, Dr. David Raible for the *Tg(neurod1:TagRFP)* transgenic fish and Dr. Clair Wyart for the *Tg(neurod1:Gcamp6f)* transgenic fish. We also thank the Duke Light Microscopy Core Facility for equipment access and technical support, the Duke Zebrafish Core Facility for assisting zebrafish husbandry and the Duke Microbiome Shared Resource for 16S rRNA gene sequencing. This work was supported by grants from the National Institutes of Health R01-DK093399 (to J.F.R. and R.A.L.), R01 DK109368 (to R.A.L.), and R01-DK081426 (to J.F.R.); the Department of Veterans Affairs I01BX002230 (to R.A.L.); and an Innovation Grant from the Pew Charitable Trusts (to J.F.R. and R.A.L.). L.Y. was supported by the Digestive Disease and Nutrition Training Program at Duke University (T32-DK007568).

## Supplemental Figures

**Supplemental Figure 1.**
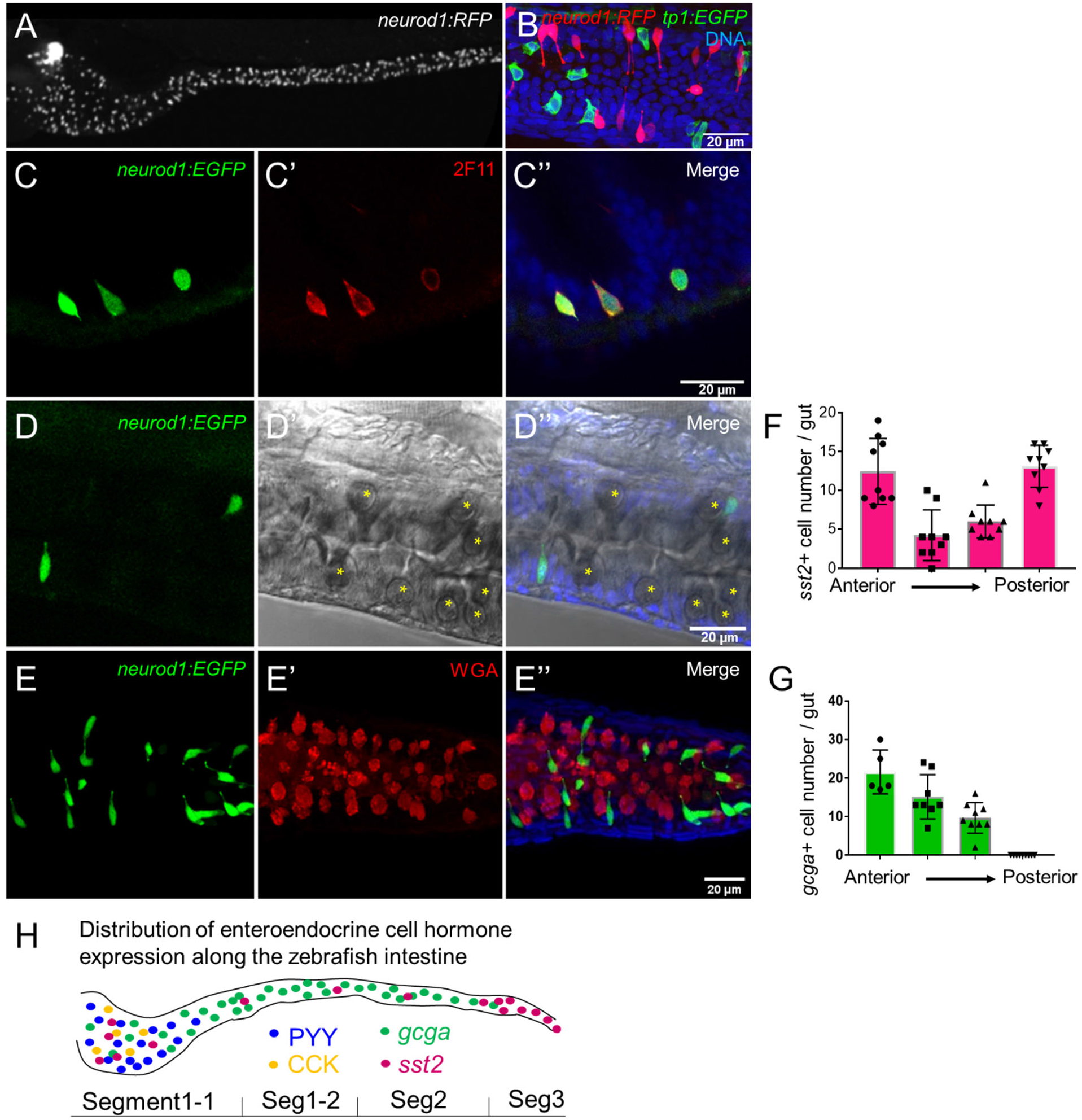
Characterization of zebrafish enteroendocrine cells. (A) Fluorescence images of *Tg(-5kbneurod1:TagRFP)* 6 dpf zebrafish gut. *Neurod1* is expressed in islet cells of the pancreas and enteroendocrine cells in the intestine. (B) Confocal projection of zebrafish EECs marked by *Tg(-5kbneurod1:TagRFP)*. Note that red *neurod1*+ EECs are not overlapping with green *tp1*+ cells. (C) Immunofluorescence staining of 6 dpf *TgBAC(neurod1:EGFP)* with the known intestinal secretory cell marker 2F11 (red). (D) Confocal plane of zebrafish intestine from *TgBAC(neurod1:EGFP)*. Goblet cells are identified by their specific cell shape in the white field (B”) and EGFP labeled EECs do not overlap with goblet cells. (E) Confocal projection of zebrafish EECs marked by *TgBAC(neurod1:EGFP)*. Mucus in Goblet cells is labeled with WGA lectin (red). *neurod1*+ EECs do not stain with WGA. (F) Quantification of somatostatin+ cells that are labeled by *Tg(sst2:RFP)* in the 6 dpf zebrafish intestine. (G) Quantification of glucagon+ cells that are labeled by *Tg(gcga:EGFP)* in the 6 dpf zebrafish intestine. (H) Schematic depiction of EEC hormone distribution along the intestinal segments of 6 dpf zebrafish larvae.

**Supplemental Figure 2.**
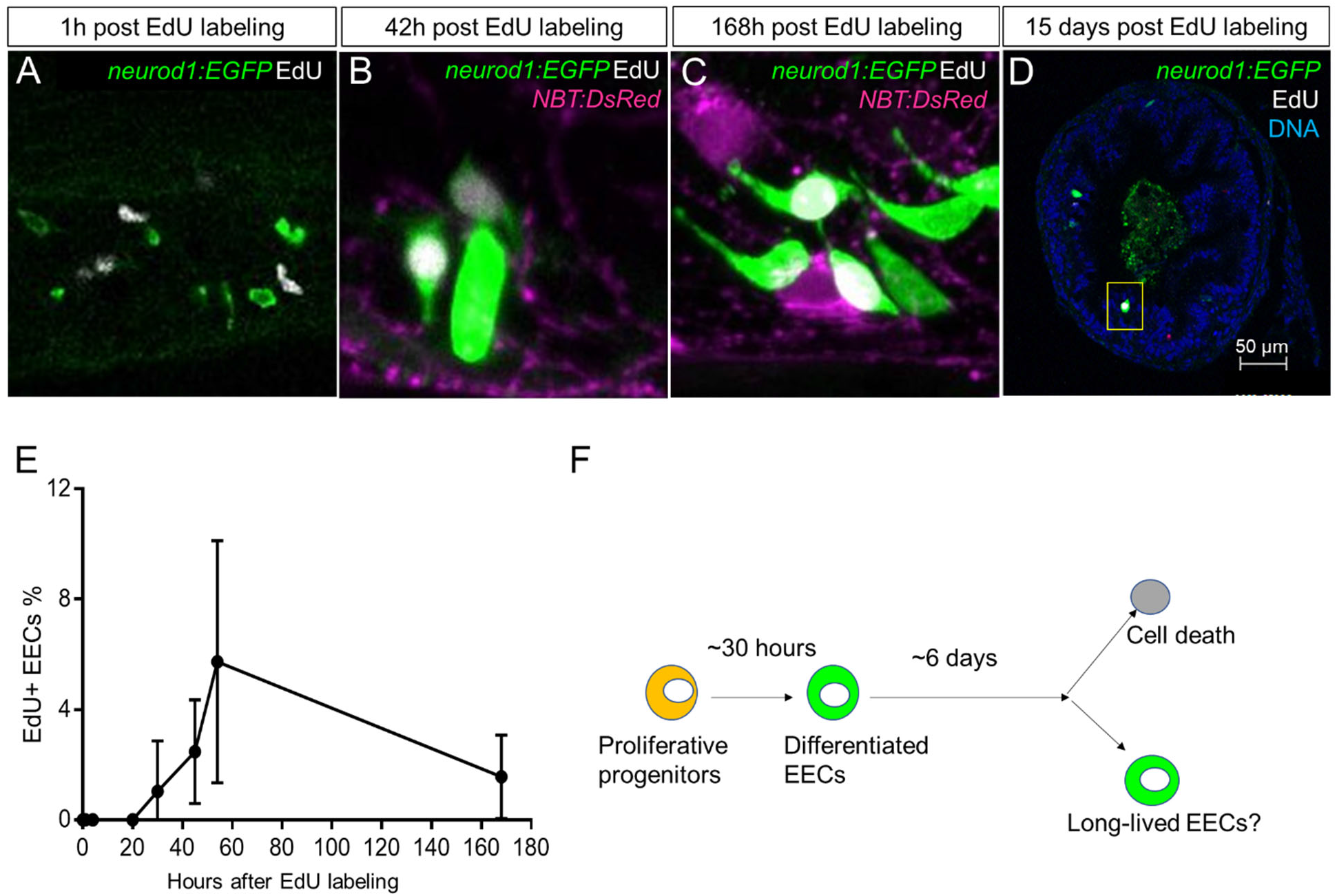
Analysis of EEC lifespan in zebrafish larvae using single dose EdU labeling. EdU was injected into the pericardiac sac region of 5 dpf *TgBAC(neurod1:EGFP)* zebrafish using described previously methods (Ye et al., 2015). Zebrafish were fixed at 1h, 4h, 20h, 30h, 45h, 54h, 7 days (168 hours) and 15 days post EdU injection. (A-D) Confocal images of EdU fluorescence staining in *TgBAC(neurod1:EGFP)* zebrafish intestine. (E) Quantification of the percentage of EdU+ EECs in zebrafish intestine following EdU tracing. t=0 (n=6), t=1h (n=8), t=4h (n=5), t=20h (n=6), t=30h (n=11), t=45h (n=9), t=54h (n=6), t=168h (n=). No EdU+EECs could be detected until 30h post EdU injection and some EdU+ EECs remained 15 days post EdU injection. (F) Schematic model of our hypothesis of the EEC lifespan.

**Supplemental Figure 3.**
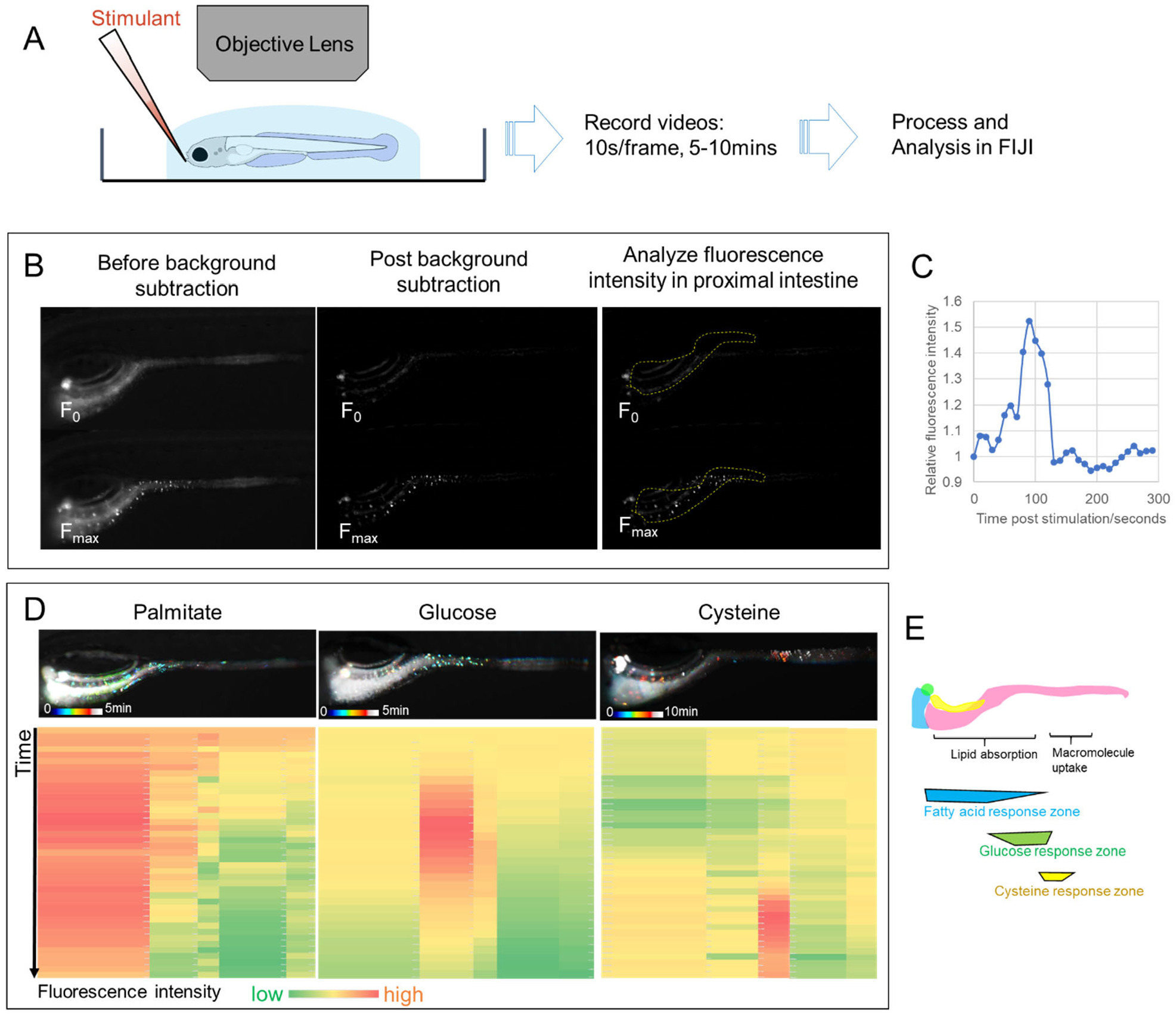
EEC activity assay. (A) Experimental flow of EEC activity assay using the *Tg(-5kbneurod1:Gcamp6f)* model. (B) Representative images of EEC calcium fluorescence analysis using FIJI template matching and background subtraction in *Tg(-5kbneurod1:Gcamp6f)* zebrafish stimulated with palmitate. (C) Relative fluorescence intensity in the proximal intestine in a series of video images from zebrafish in B. (D) Spatial-temporal resolution of the EEC response to palmitate, glucose and cysteine stimulation. (E) Representative images of the EEC nutrient response in a regional specific manner. Palmitate and glucose primarily activated EECs in the proximal intestine where most lipid and nutrient absorption occurs. Cysteine on the other hand activated EECs in the distal intestine where proteins are digested and amino acids absorbed by specialized intestinal epithelial cells in this region (Nakamura, Tazumi, Muro, Yasuhara, & Watanabe, 2004; Wang et al., 2010).

**Supplemental Figure 4.**
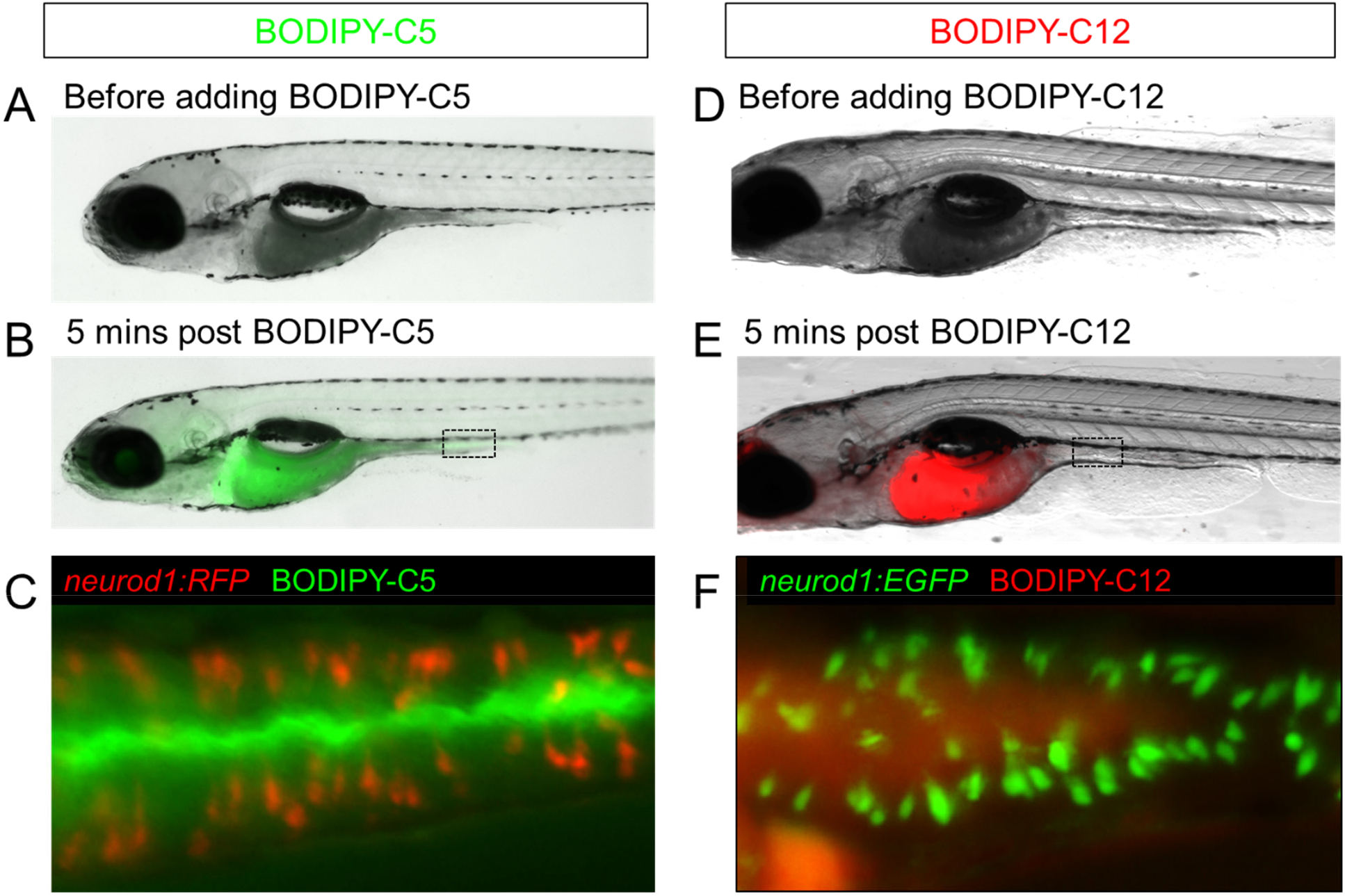
Feeding a high fat meal did not impair fatty acid intake. (A-C) Fatty acid was labeled with green fluorescence in BODIPY-C5 (Carten et al., 2011). BODIPY-C5 (in BSA complex) was delivered to zebrafish larvae that had been fed high fat (HF) meal for 6 hours, the same as the EEC activity assay. Within 5 minutes of delivery, green BODIPY-C5 was distributed throughout the entire zebrafish intestinal lumen. (D-F) Fatty acids were labeled with red fluorescence in BODIPY-C12 (Carten et al., 2011). BODIPY-C12 was delivered to zebrafish larvae that had been fed HF meal for 6 hours, the same as the EEC activity assay. Within 5 minutes of delivery, red BODIPY-C12 was distributed throughout the zebrafish intestinal lumen.

**Supplemental Figure 5.**
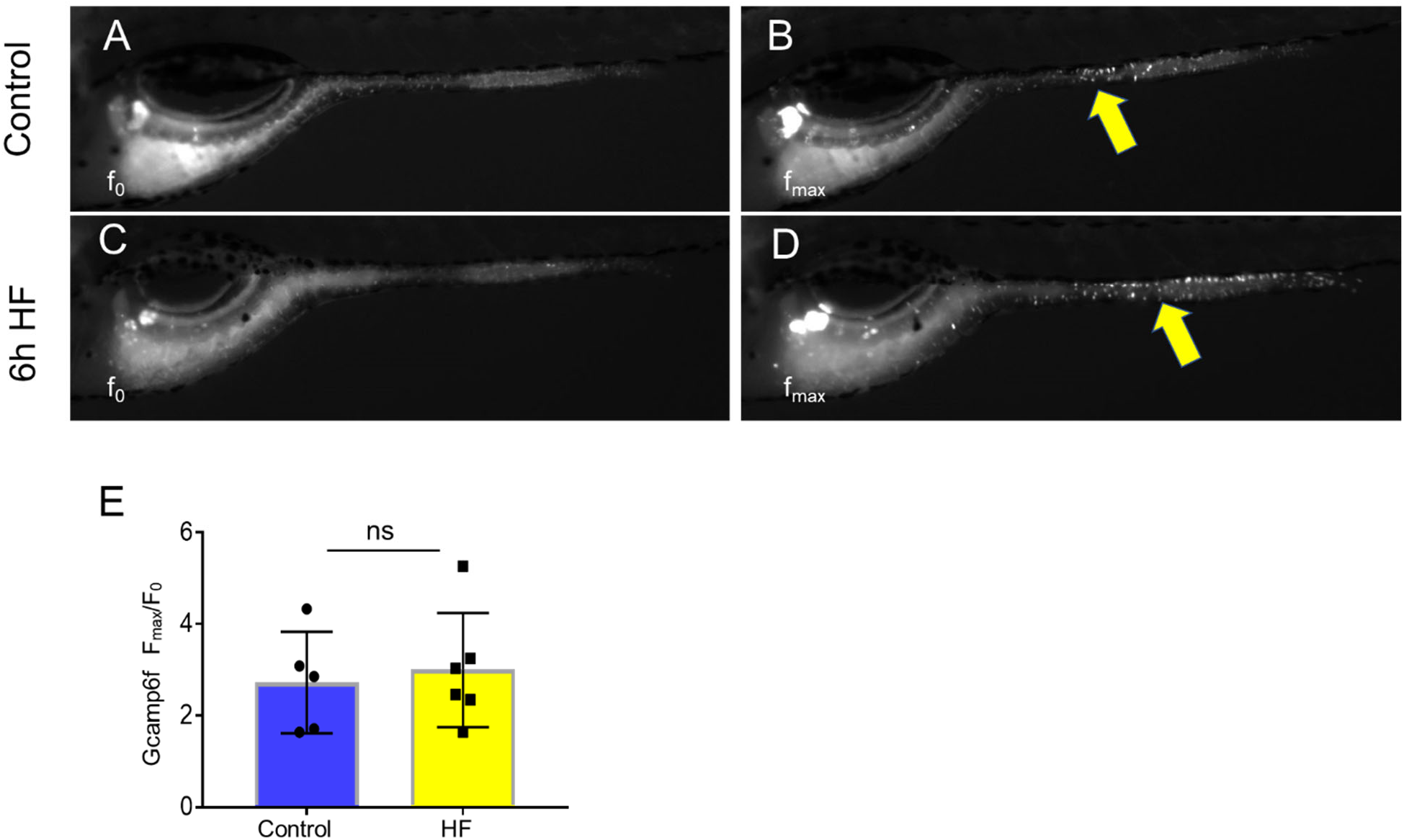
EECs remain responsive to cysteine following high fat feeding. (A-B) Representative images of the EEC response to cysteine in control *Tg(neurod1:Gcamp6f)* zebrafish larvae. Note the location of responsive EECs in the mid-intestinal region (yellow arrows) (C-D) Representative images of the EEC response to cysteine in 6h high fat meal fed *Tg(neurod1:Gcamp6f)* zebrafish larvae. (E) Quantification of the EEC response to cysteine in control and high fat fed zebrafish. Student t-test was used in E for statistical analysis.

**Supplemental Figure 6.**
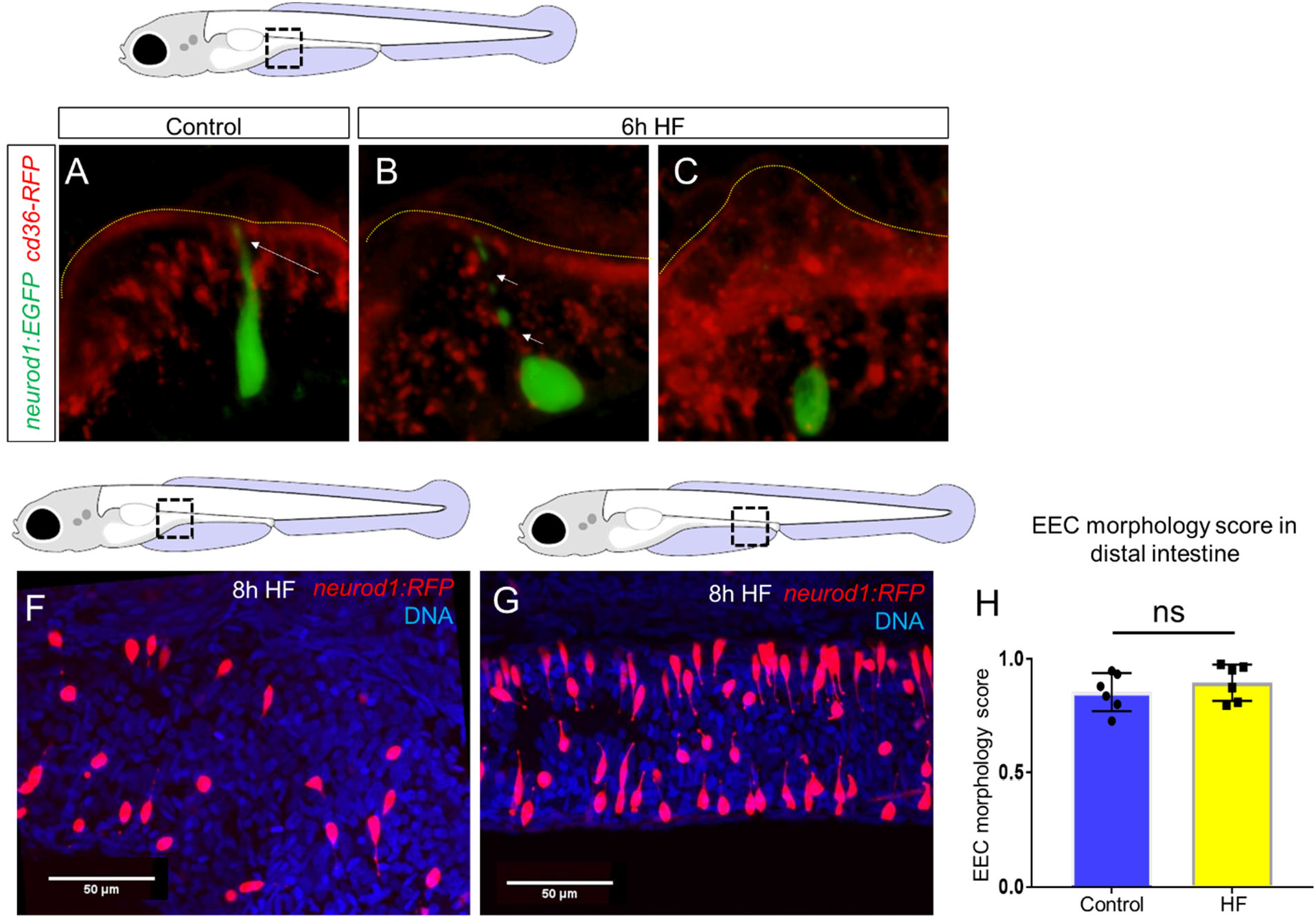
High fat feeding induced loss of the EEC apical protrusion in the proximal intestine but not in the distal intestine. (A) Confocal projection of a typical EEC of control *TgBAC(neurod1:EGFP); TgBAC(CD36-RFP)* zebrafish. The white arrow indicates the apical projection that extends to the intestinal lumen. (B) Confocal image of an EEC 6 hours post high fat (HF) meal feeding in *TgBAC(neurod1:EGFP); TgBAC(CD36-RFP)* zebrafish. The white arrows indicate the discontinuous fragmentation of an apical projection that can only be observed in HF fed EECs. (C) Confocal image of “closed” EECs after 6 hours post HF meal feeding in *TgBAC(neurod1:EGFP); TgBAC(CD36-RFP)* zebrafish. (F) Representative confocal image of EECs in the proximal intestine following 8 hours of high fat feeding. (G) Representative confocal image of EECs in the distal intestine following 8h HF feeding. (H) Quantification of EEC morphology in the distal intestine in control and 8h HF fed zebrafish. Student t-test was used in H for statistical analysis.

**Supplemental Figure 7.**
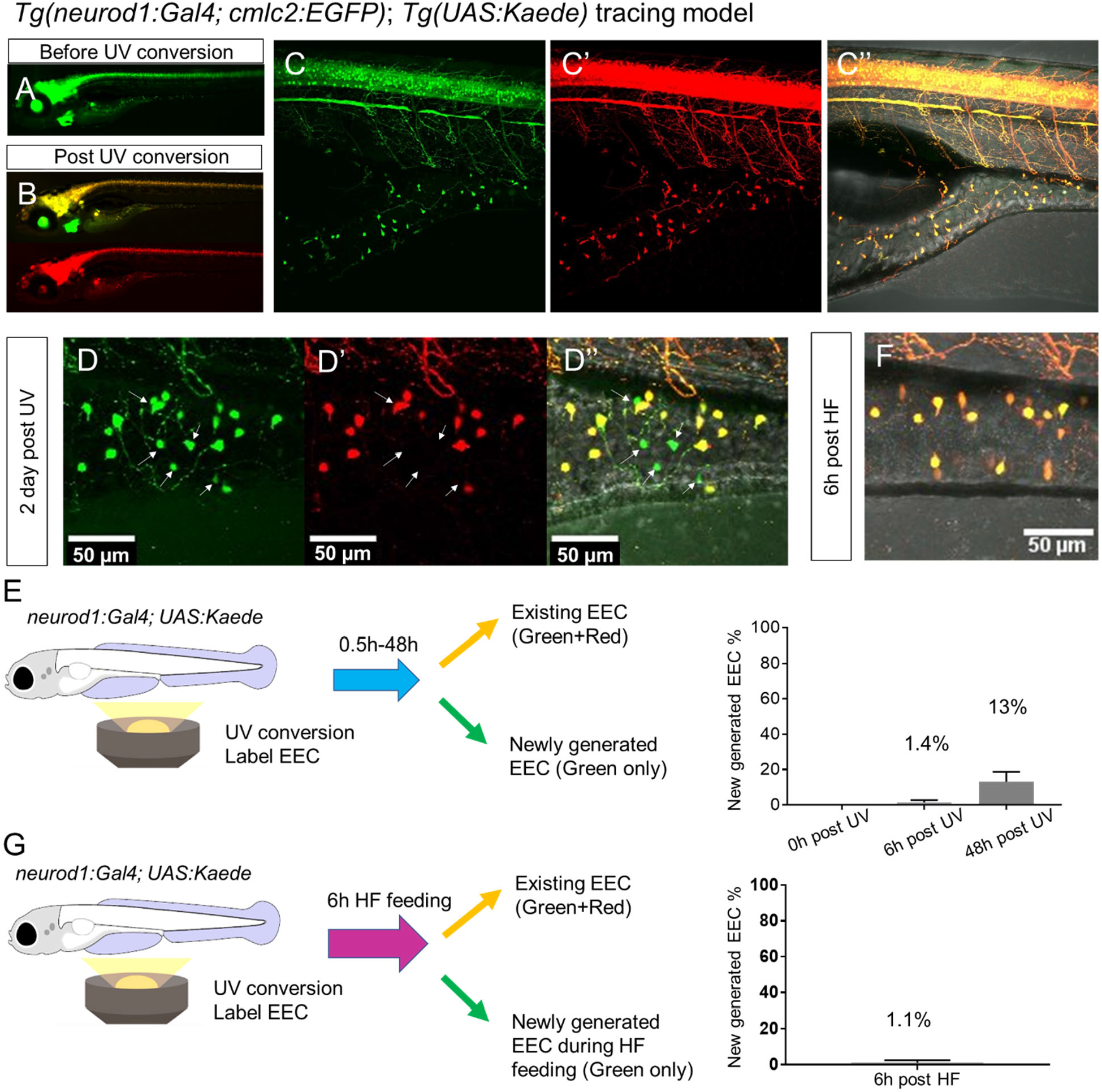
High fat feeding did not induce EEC neogenesis. (A-B) Epifluorescence image of *in vivo* EEC lineage tracing using *Tg(neurod1:Gal4;cmlc2:EGFP); Tg(UAS:Kaede)* (referred to as *neurod1-Kaede*). Before UV conversion, neurod1+ cells were labeled with green Kaede protein. Following UV exposure, green Kaede protein was converted into red Kaede protein and neurod1+ cells are labeled yellow. (C) Confocal image of live *neurod1-Kaede* zebrafish intestine 0.5h post UV conversion. All the EECs are labeled. (D) Confocal image of live *neurod1-Kaede* zebrafish intestine 2 days post UV conversion. Arrows indicate the EECs that were generated after UV conversion and exhibit green fluorescence only. (E) Confocal image of live *neurod1-Kaede* zebrafish intestine 6 hours post high fat (HF) meal. No green EECs were detected. (F) EEC neogenesis tracing at 0.5 hour, 6 hours and 2 days post UV conversion using the *neurod1-Kaede* system. (G) EEC neogenesis tracing at 6 hours post HF feeding.

**Supplemental Figure 8.**
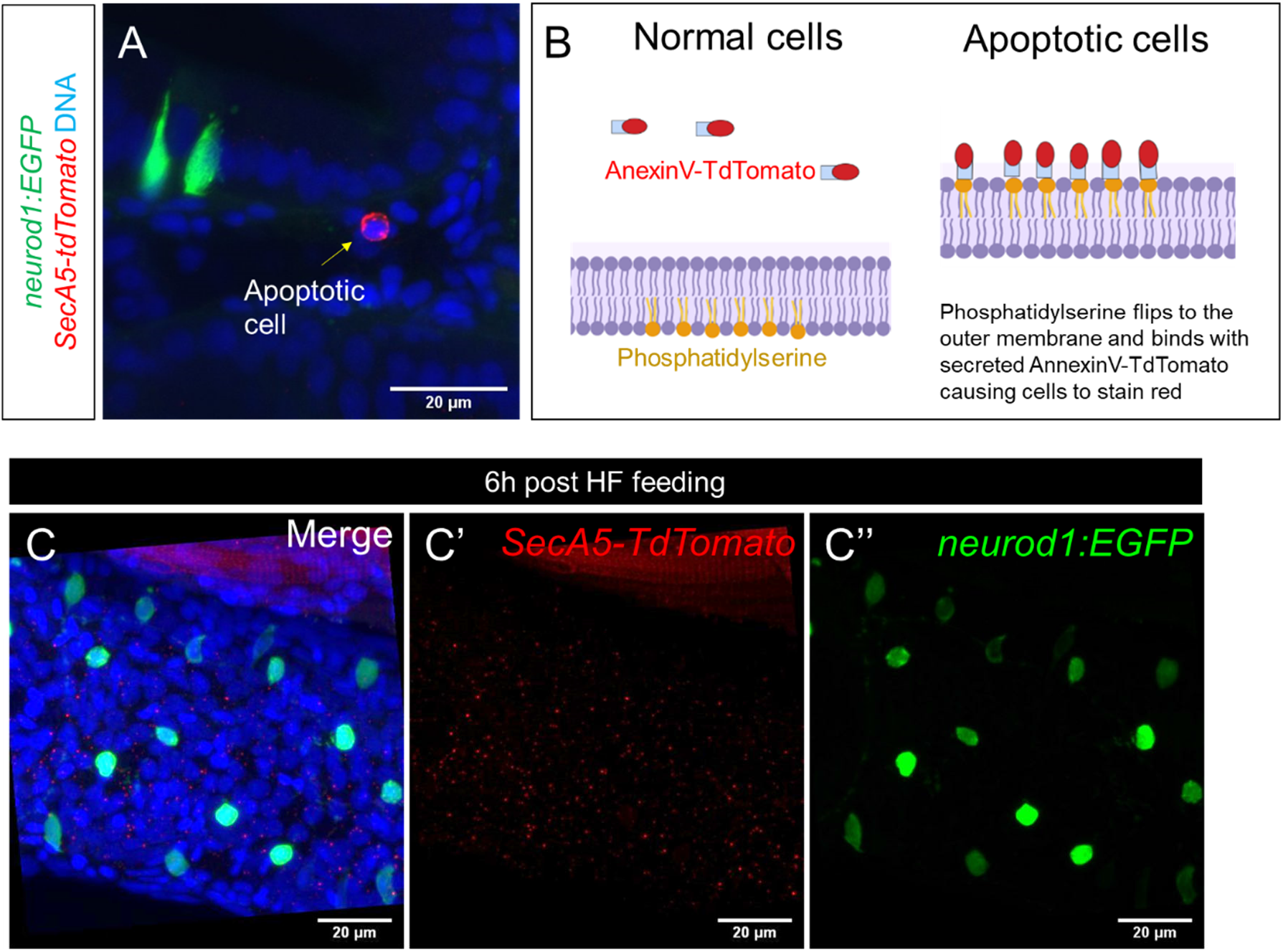
High fat feeding did not induce EEC apoptosis. (A) Confocal projection of 6 dpf *Tg(neurod1:EGFP); Tg(ubb:secA5-TdTomato)* intestine. The apoptotic cells were labeled by sesA5-tdTomato as red (yellow arrow). (B) Schematic view of labeling apoptotic cells using sesA5-tdTomato (secreted Annexin5-TdTomato). During apoptosis, phosphatidylserine flips to the outer cellular membrane. The secA5-TdTomato was then able to bind to the phosphatidylserine and labeled the apoptotic cells red. (C) Confocal image of *Tg(neurod1:EGFP); Tg(ubb:secA5-TdTomato)* zebrafish intestine following 6 hours of the high fat (HF) meal. In all the samples that were examined (n=10), no apoptotic EECs were observed.

**Supplemental Figure 9.**
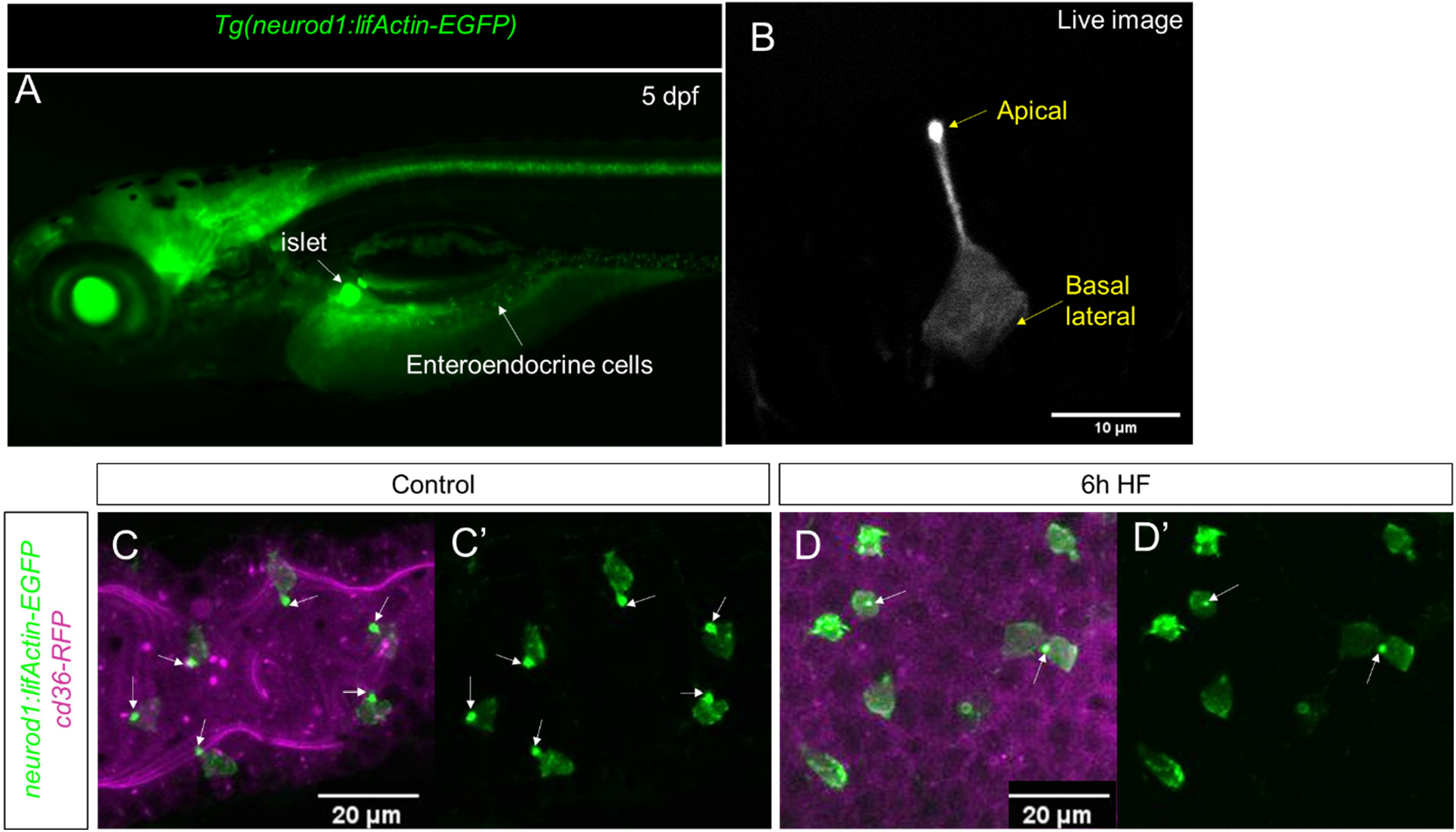
Characterization of *Tg(neurod1:lifActin-EGFP)*. (A) Epifluoresence image of 5 dpf *Tg(neurod1:lifActin-EGFP)*. The pancreatic islet and enteroendocrine cells that were labeled by lifActin-EGFP are designated by white arrows. (B) Confocal image of EEC labeled by*Tg(neurod1:lifActin-EGFP)* in live zebrafish mounted in 2% low melting agarose. The stronger lifActin-EGFP signal was detected in the apical of EEC protrusion. (C, C’) Confocal image of *Tg(neurod1:lifActin-EGFP); TgBAC(cd36-RFP)* zebrafish intestine. The EEC’s apical protrusion that labeled by a strong lifActin-EGFP signal is labeled with white arrows. (D, D’) Confocal images of *Tg(neurod1: lifActin-EGFP); TgBAC(cd36-RFP)* zebrafish intestine after 6h high fat (HF) meal feeding. The EECs’ apical protrusion labeled by strong lifActin-EGFP signal was reduced.

**Supplemental Figure 10.**
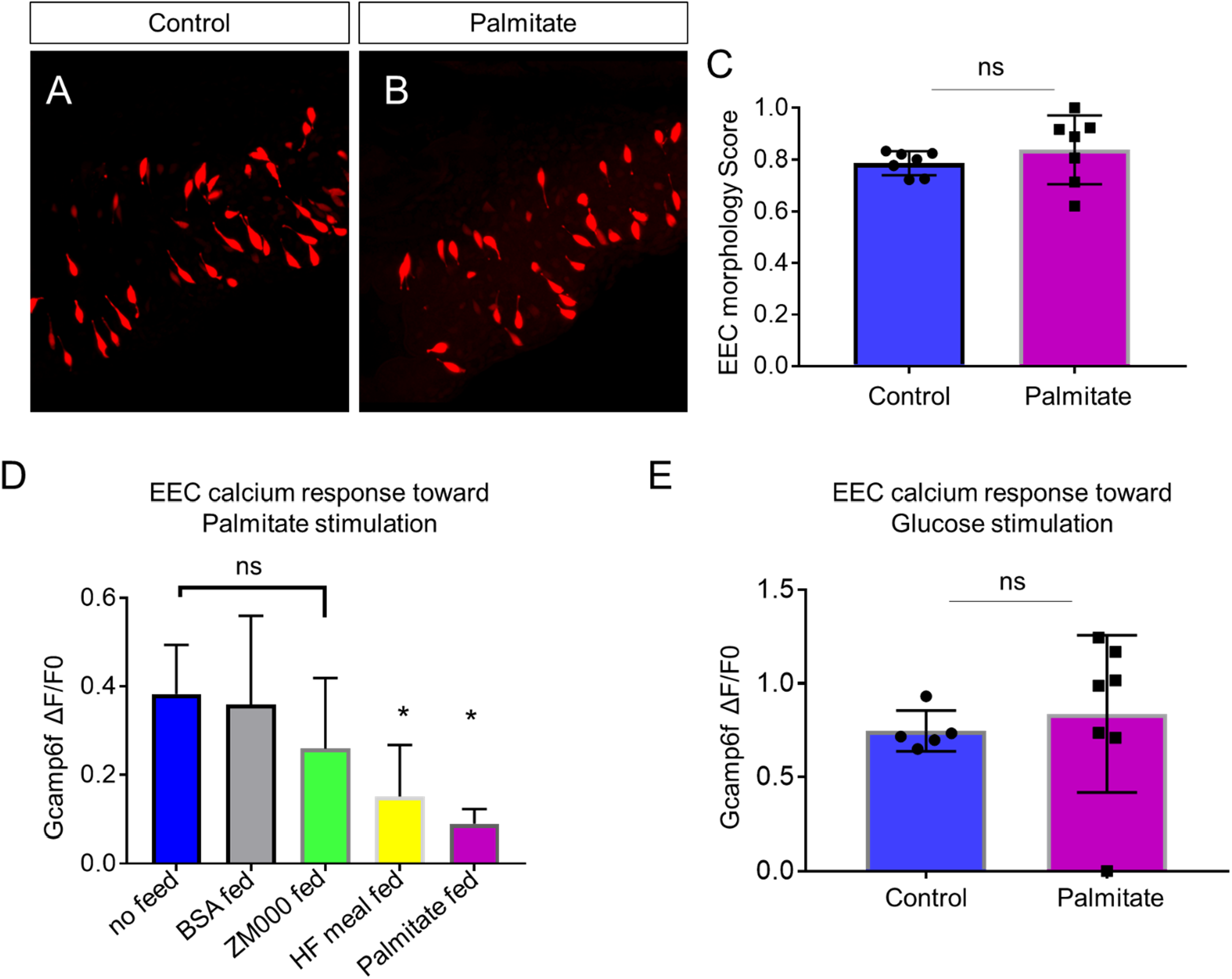
The effect of palmitate feeding on EEC morphology and function. (A) Confocal image of *Tg(neurod1:TagRFP)* zebrafish intestine in control and 6 hours post palmitate feeding zebrafish. (B) EEC morphology score in control and palmitate fed zebrafish embryos. (D) EECs’ response to palmitate following different dietary manipulations. 6 dpf *Tg(neurod1:Gcamp6f)* zebrafish were untreated (n=6) or fed for 6 hours with BSA (n=3), ZM000 (larvae zebrafish food) (n=4), HF meal (6.25% chicken egg yolk) (n=11), or palmitate (n=4). Only egg yolk and palmitate feeding reduced the EECs’ response to palmitate stimulation. (E) EECs’ response to glucose stimulation following 6 hours of palmitate feeding. Student t-test was used in C,E and one-way ANOVA with post-hoc Tukey test was used in D for statisc analysis. * P<0.05, ns P>0.05, not signficicantly different.

**Supplemental Figure 11.**
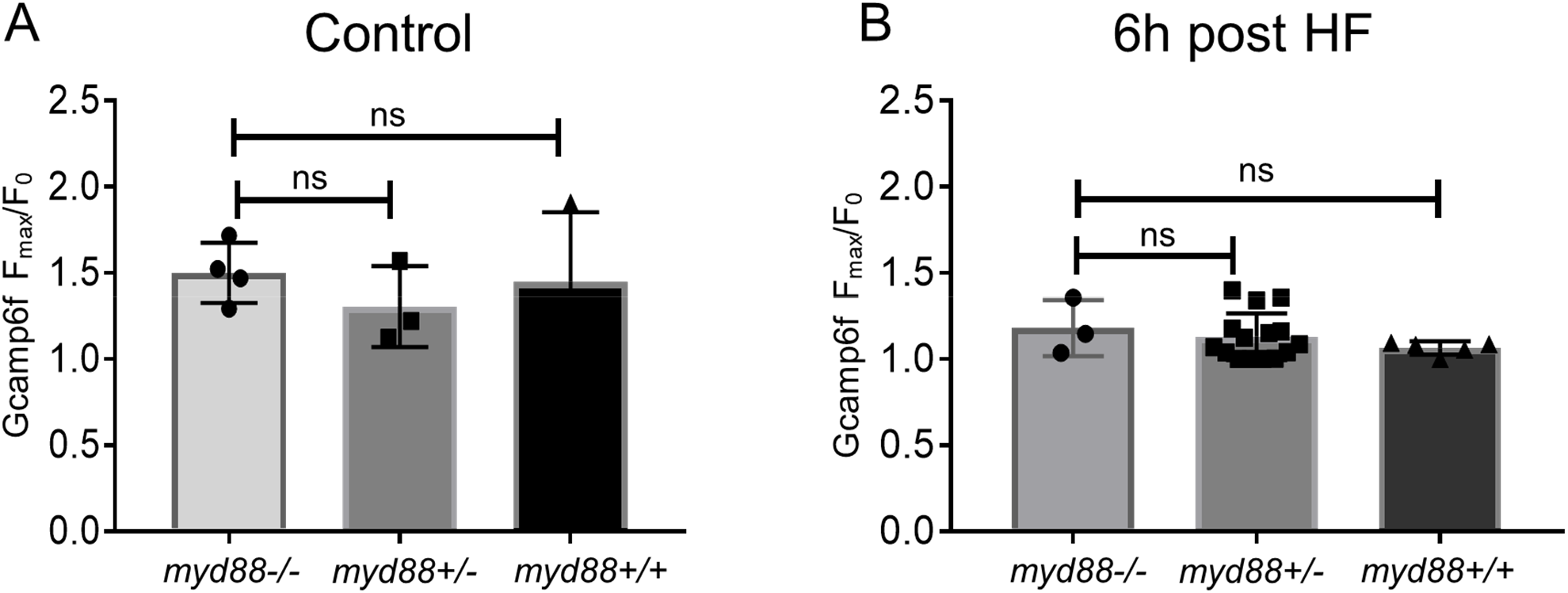
EEC sensitivity to palmitate stimulation is not altered in *myd88* mutant zebrafish. *Tg(neurod1:Gcamp6f)*; *myd88*+/− fish were crossed with *Tg(neurod1:Gcamp6f)*+ fish and sorted at 3 dpf. Response to palmitate stimulation was assessed afterwhich the genotypes of zebrafish were determined. (A) Quantification of the EEC response to palmitate stimulation in 6 dpf *Tg(neurod1:Gcamp6f)* zebrafish under control conditions. No differences were observed among different genotypes. (B) Quantification of EECs’ response to palmitate stimulation in 6 dpf *Tg(neurod1:Gcamp6f)* zebrafish 6 hours after high fat (HF) feeding. One-way ANOVA with post-hoc Tukey test was used in A, B for statistical analysis and no statistical differences were observed among different genotypes.

**Supplemental Figure 12.**
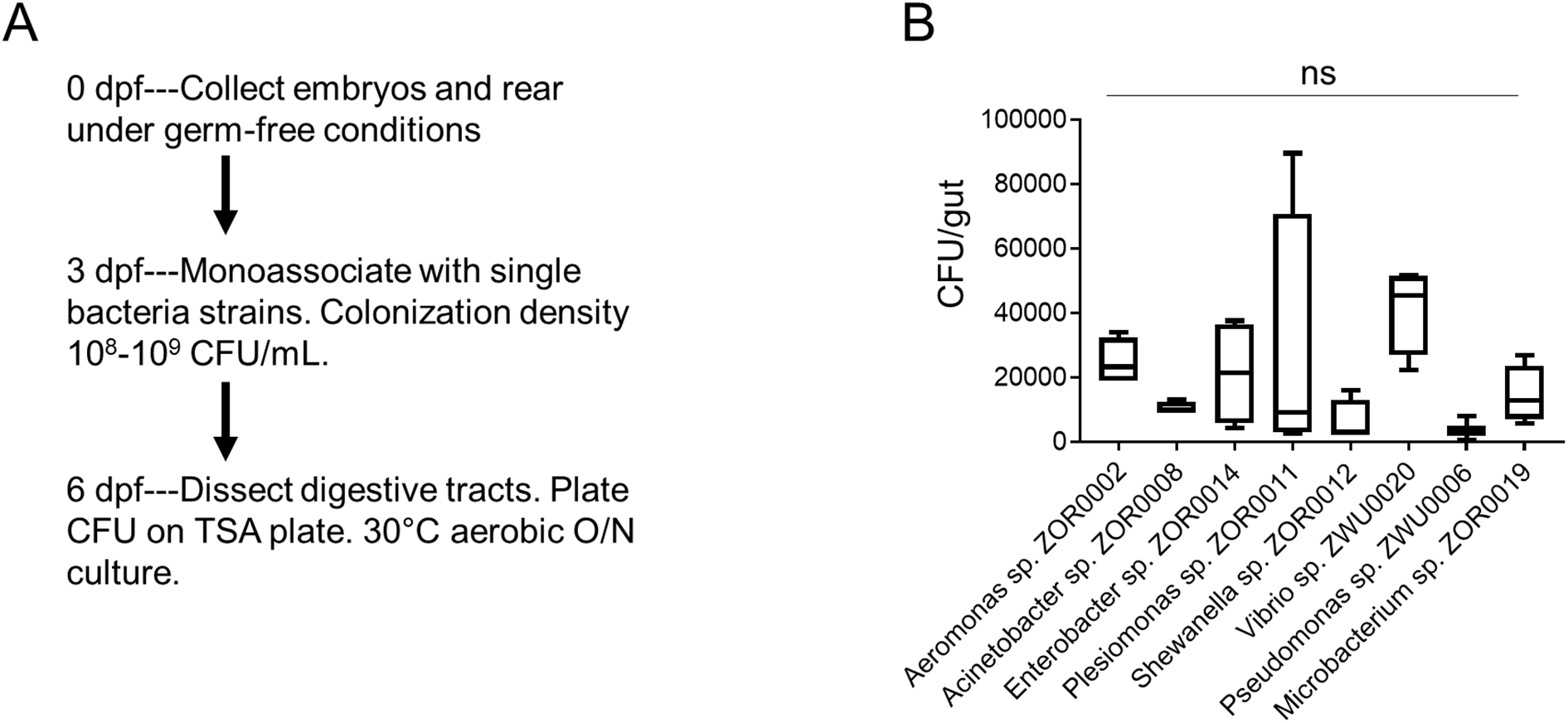
Colonization of bacteria strains used for monoassociation. (A) Schematics of the experimental design. The digestive tracts from 5 zebrafish larvae were dissected and pooled, and CFU analysis was performed to assess the colonization efficiency for bacteria strains that were used for consociation. (B) CFU quantification of zebrafish larvae samples that were monoassociated with different bacterial strains. One-way ANOVA with post-hoc Tukey test was used in B for statistic analysis and no statistical differences among groups were observed.

**Supplemental Figure 13.**
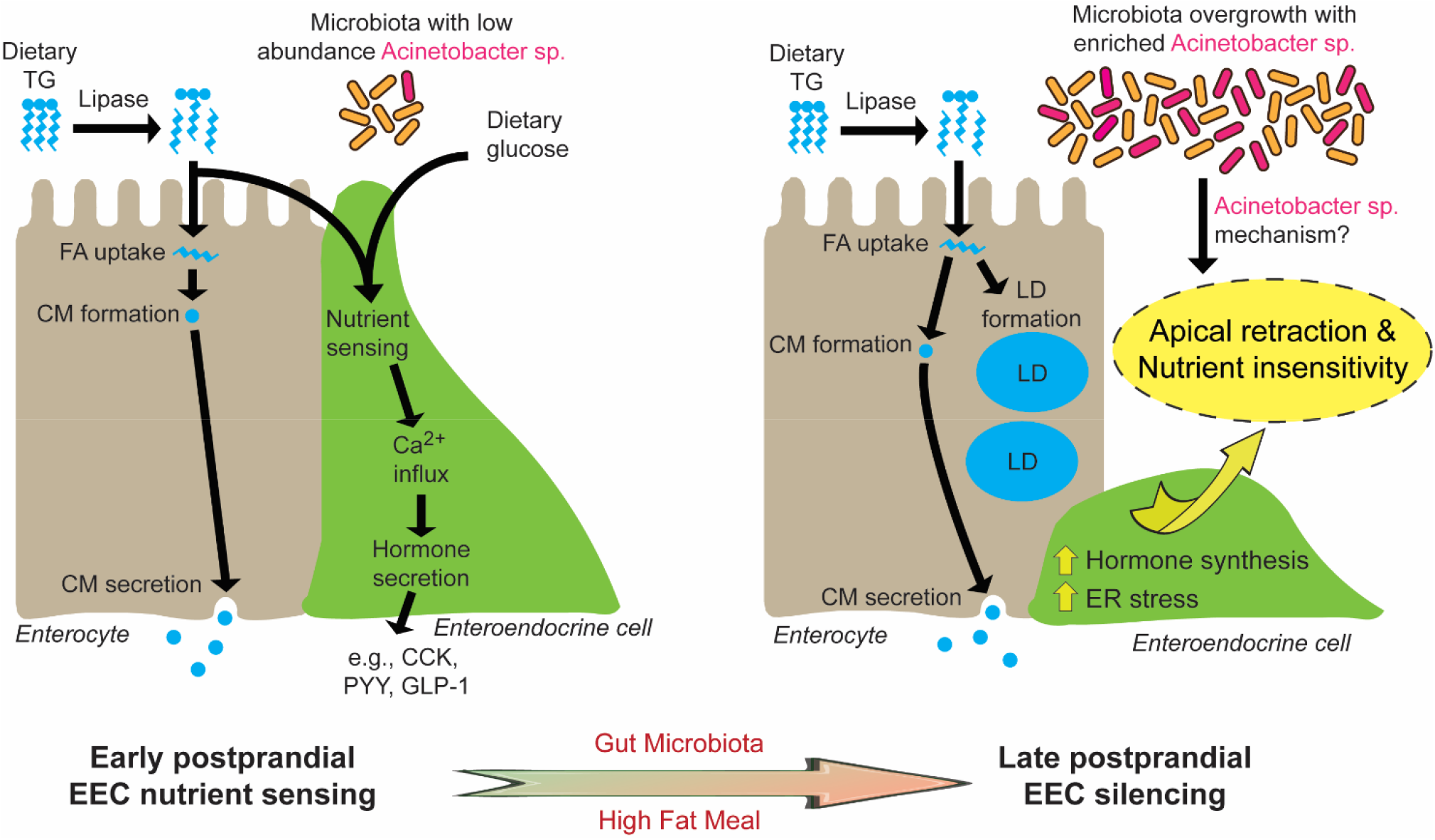
Proposed model for microbiota-dependent HF feeding-induced EEC silencing. At early postprandial stages after consumption of a HF meal, dietary triglyceride (TG) is hydrolyzed to monoglycerides and free fatty acids (FA) by lipases in the gut lumen. FA are taken up by enterocytes and re-esterified into TG which is packaged into chylomicrons (CM) for basolateral secretion. FA and dietary glucose stimulate EECs, increasing [Ca^2+^]_i_ and inducing secretion of hormones like CCK, PYY and GLP-1. During and after HF feeding, FA taken up by enterocytes are stored in cytosolic lipid droplets (LD) in addition to secreted CM. During later postprandial stages, nutrient over-stimulation from HF feeding increases EECs hormone synthesis burden, activates EECs ER stress response pathways and indues overgrowth of the gut bacterial community including enrichment of *Acinetobacter* sp.. These nutrient and microbial stimuli in turn induce EECs apical retraction and reduce EECs nutrient sensitivity at the late postprandial stage, a process we call “EEC silencing”.

**Supplemental video 1.** EEC calcium response to water, BSA, palmitate, glucose and cysteine administration. 6 dpf *Tg(neurod1:Gcamp6f)* zebrafish were used in these experiments and videos were recorded at 10s/frame for 5 or 10 minutes (cysteine).

**Supplemental video 2.** EEC calcium response to palmitate stimulation in control and 6 hour high fat fed *Tg(neurod1:Gcamp6f)* zebrafish larvae.

**Supplemental video 3.** EEC calcium response to glucose stimulation in control and 6 hours high fat fed *Tg(neurod1:Gcamp6f)* zebrafish larvae.

**Supplemental video 4.** Confocal Z stack images in *Tg(neurod1:TagRFP); TgBAC(gata5:lifActin-EGFP)* 6 dpf control zebrafish larvae. The apical border of the intestine was labeled by gata5:lifActin-EGFP. Note that the apical protrusion of EECs extends to the intestinal lumen.

**Supplemental video 5.** Confocal Z stack image in *Tg(neurod1:TagRFP); TgBAC(gata5:lifActin-EGFP)* 10 hours high fat fed zebrafish larvae. The majority of EECs have lost their apical protrusions that extend to the intestinal lumen.

**Supplemental video 6.** Time lapse video of intestine in control *Tg(neurod1:lifActin-EGFP)* zebrafish larvae. Videos were taken at 10s/frame for 16 minutes.

## References

Aghaallaei, N., Gruhl, F., Schaefer, C. Q., Wernet, T., Weinhardt, V., Centanin, L., … Wittbrodt, J. (2016). Identification, visualization and clonal analysis of intestinal stem cells in fish. Development, 143(19), 3470–3480. Retrieved from https://www.ncbi.nlm.nih.gov/pubmed/27578784. doi:10.1242/dev.134098

Arora, T., Akrami, R., Pais, R., Bergqvist, L., Johansson, B. R., Schwartz, T. W., … Backhed, F. (2018). Microbial regulation of the L cell transcriptome. Sci Rep, 8(1), 1207. Retrieved from https://www.ncbi.nlm.nih.gov/pubmed/29352262. doi:10.1038/s41598-017-18079-2

Austin, G. L., Ogden, L. G., & Hill, J. O. (2011). Trends in carbohydrate, fat, and protein intakes and association with energy intake in normal-weight, overweight, and obese individuals: 1971-2006. Am J Clin Nutr, 93(4), 836–843. Retrieved from https://www.ncbi.nlm.nih.gov/pubmed/21310830. doi:10.3945/ajcn.110.000141

Backhed, F., Manchester, J. K., Semenkovich, C. F., & Gordon, J. I. (2007). Mechanisms underlying the resistance to diet-induced obesity in germ-free mice. Proc Natl Acad Sci U S A, 104(3), 979–984. Retrieved from https://www.ncbi.nlm.nih.gov/pubmed/17210919. doi:10.1073/pnas.0605374104

Ballinger, A. (2000). Orlistat in the treatment of obesity. Expert Opin Pharmacother, 1(4), 841–847. Retrieved from https://www.ncbi.nlm.nih.gov/pubmed/11249520. doi:10.1517/14656566.1.4.841

Bauer, P. V., Duca, F. A., Waise, T. M. Z., Rasmussen, B. A., Abraham, M. A., Dranse, H. J., … Lam, T. K. T. (2018). Metformin Alters Upper Small Intestinal Microbiota that Impact a Glucose-SGLT1-Sensing Glucoregulatory Pathway. Cell Metab, 27(1), 101–117 e105. Retrieved from https://www.ncbi.nlm.nih.gov/pubmed/29056513. doi:10.1016/j.cmet.2017.09.019

Beglinger, C., & Degen, L. (2006). Gastrointestinal satiety signals in humans--physiologic roles for GLP-1 and PYY? Physiol Behav, 89(4), 460–464. Retrieved from https://www.ncbi.nlm.nih.gov/pubmed/16828127. doi:10.1016/j.physbeh.2006.05.048

Bohorquez, D. V., Shahid, R. A., Erdmann, A., Kreger, A. M., Wang, Y., Calakos, N., … Liddle, R. A. (2015). Neuroepithelial circuit formed by innervation of sensory enteroendocrine cells. J Clin Invest, 125(2), 782–786. Retrieved from https://www.ncbi.nlm.nih.gov/pubmed/25555217. doi:10.1172/JCI78361

Bouvet, P. J., & Jeanjean, S. (1989). Delineation of new proteolytic genomic species in the genus Acinetobacter. Res Microbiol, 140(4-5), 291–299. Retrieved from https://www.ncbi.nlm.nih.gov/pubmed/2799067.

Burns, A. R., Miller, E., Agarwal, M., Rolig, A. S., Milligan-Myhre, K., Seredick, S., … Bohannan, B. J. M. (2017). Interhost dispersal alters microbiome assembly and can overwhelm host innate immunity in an experimental zebrafish model. Proc Natl Acad Sci U S A, 114(42), 11181–11186. Retrieved from https://www.ncbi.nlm.nih.gov/pubmed/28973938. doi:10.1073/pnas.1702511114

Callahan, B. J., McMurdie, P. J., Rosen, M. J., Han, A. W., Johnson, A. J., & Holmes, S. P. (2016). DADA2: High-resolution sample inference from Illumina amplicon data. Nat Methods, 13(7), 581–583. Retrieved from https://www.ncbi.nlm.nih.gov/pubmed/27214047. doi:10.1038/nmeth.3869

Caporaso, J. G., Kuczynski, J., Stombaugh, J., Bittinger, K., Bushman, F. D., Costello, E. K., … Knight, R. (2010). QIIME allows analysis of high-throughput community sequencing data. Nat Methods, 7(5), 335–336. Retrieved from https://www.ncbi.nlm.nih.gov/pubmed/20383131. doi:10.1038/nmeth.f.303

Carten, J. D., Bradford, M. K., & Farber, S. A. (2011). Visualizing digestive organ morphology and function using differential fatty acid metabolism in live zebrafish. Dev Biol, 360(2), 276–285. Retrieved from https://www.ncbi.nlm.nih.gov/pubmed/21968100. doi:10.1016/j.ydbio.2011.09.010

Chandra, R., Hiniker, A., Kuo, Y. M., Nussbaum, R. L., & Liddle, R. A. (2017). alpha-Synuclein in gut endocrine cells and its implications for Parkinson’s disease. JCI Insight, 2(12). Retrieved from https://www.ncbi.nlm.nih.gov/pubmed/28614796. doi:10.1172/jci.insight.92295

Chandra, R., Wang, Y., Shahid, R. A., Vigna, S. R., Freedman, N. J., & Liddle, R. A. (2013). Immunoglobulin-like domain containing receptor 1 mediates fat-stimulated cholecystokinin secretion. J Clin Invest, 123(8), 3343–3352. Retrieved from https://www.ncbi.nlm.nih.gov/pubmed/23863714. doi:10.1172/JCI68587

Crosnier, C., Vargesson, N., Gschmeissner, S., Ariza-McNaughton, L., Morrison, A., & Lewis, J. (2005). Delta-Notch signalling controls commitment to a secretory fate in the zebrafish intestine. Development, 132(5), 1093–1104. Retrieved from https://www.ncbi.nlm.nih.gov/pubmed/15689380. doi:10.1242/dev.01644

Cuenco, J., Minnion, J., Tan, T., Scott, R., Germain, N., Ling, Y., … Bloom, S. (2017). Degradation Paradigm of the Gut Hormone, Pancreatic Polypeptide, by Hepatic and Renal Peptidases. Endocrinology, 158(6), 1755–1765. Retrieved from https://www.ncbi.nlm.nih.gov/pubmed/28323997. doi:10.1210/en.2016-1827

David, L. A., Maurice, C. F., Carmody, R. N., Gootenberg, D. B., Button, J. E., Wolfe, B. E., … Turnbaugh, P. J. (2014). Diet rapidly and reproducibly alters the human gut microbiome. Nature, 505(7484), 559–563. Retrieved from https://www.ncbi.nlm.nih.gov/pubmed/24336217. doi:10.1038/nature12820

Delzenne, N. M., Cani, P. D., & Neyrinck, A. M. (2007). Modulation of glucagon-like peptide 1 and energy metabolism by inulin and oligofructose: experimental data. J Nutr, 137(11 Suppl), 2547S–2551S. Retrieved from https://www.ncbi.nlm.nih.gov/pubmed/17951500. doi:10.1093/jn/137.11.2547S

Donaldson, J. G., Cassel, D., Kahn, R. A., & Klausner, R. D. (1992). ADP-ribosylation factor, a small GTP-binding protein, is required for binding of the coatomer protein beta-COP to Golgi membranes. Proc Natl Acad Sci U S A, 89(14), 6408–6412. Retrieved from https://www.ncbi.nlm.nih.gov/pubmed/1631136.

Druce, M. R., Minnion, J. S., Field, B. C., Patel, S. R., Shillito, J. C., Tilby, M., … Bloom, S. R. (2009). Investigation of structure-activity relationships of Oxyntomodulin (Oxm) using Oxm analogs. Endocrinology, 150(4), 1712–1722. Retrieved from https://www.ncbi.nlm.nih.gov/pubmed/19074579. doi:10.1210/en.2008-0828

Drucker, D. J., Habener, J. F., & Holst, J. J. (2017). Discovery, characterization, and clinical development of the glucagon-like peptides. J Clin Invest, 127(12), 4217–4227. Retrieved from https://www.ncbi.nlm.nih.gov/pubmed/29202475. doi:10.1172/JCI97233

Edfalk, S., Steneberg, P., & Edlund, H. (2008). Gpr40 is expressed in enteroendocrine cells and mediates free fatty acid stimulation of incretin secretion. Diabetes, 57(9), 2280–2287. Retrieved from https://www.ncbi.nlm.nih.gov/pubmed/18519800. doi:10.2337/db08-0307

Faith, D. P., & Baker, A. M. (2007). Phylogenetic diversity (PD) and biodiversity conservation: some bioinformatics challenges. Evol Bioinform Online, 2, 121–128. Retrieved from https://www.ncbi.nlm.nih.gov/pubmed/19455206.

Farber, S. A., Pack, M., Ho, S. Y., Johnson, I. D., Wagner, D. S., Dosch, R., … Halpern, M. E. (2001). Genetic analysis of digestive physiology using fluorescent phospholipid reporters. Science, 292(5520), 1385–1388. Retrieved from https://www.ncbi.nlm.nih.gov/pubmed/11359013. doi:10.1126/science.1060418

Furness, J. B., Rivera, L. R., Cho, H. J., Bravo, D. M., & Callaghan, B. (2013). The gut as a sensory organ. Nat Rev Gastroenterol Hepatol, 10(12), 729–740. Retrieved from https://www.ncbi.nlm.nih.gov/pubmed/24061204. doi:10.1038/nrgastro.2013.180

Gainetdinov, R. R., Premont, R. T., Bohn, L. M., Lefkowitz, R. J., & Caron, M. G. (2004). Desensitization of G protein-coupled receptors and neuronal functions. Annu Rev Neurosci, 27, 107–144. Retrieved from https://www.ncbi.nlm.nih.gov/pubmed/15217328. doi:10.1146/annurev.neuro.27.070203.144206

Gerner-Smidt, P., Tjernberg, I., & Ursing, J. (1991). Reliability of phenotypic tests for identification of Acinetobacter species. J Clin Microbiol, 29(2), 277–282. Retrieved from https://www.ncbi.nlm.nih.gov/pubmed/2007635.

Goldspink, D. A., Reimann, F., & Gribble, F. M. (2018). Models and Tools for Studying Enteroendocrine Cells. Endocrinology, 159(12), 3874–3884. Retrieved from https://www.ncbi.nlm.nih.gov/pubmed/30239642. doi:10.1210/en.2018-00672

Graessler, J., Qin, Y., Zhong, H., Zhang, J., Licinio, J., Wong, M. L., … Bornstein, S. R. (2013). Metagenomic sequencing of the human gut microbiome before and after bariatric surgery in obese patients with type 2 diabetes: correlation with inflammatory and metabolic parameters. Pharmacogenomics J, 13(6), 514–522. Retrieved from https://www.ncbi.nlm.nih.gov/pubmed/23032991. doi:10.1038/tpj.2012.43

Green, C. J., & Hodson, L. (2014). The influence of dietary fat on liver fat accumulation. Nutrients, 6(11), 5018–5033. Retrieved from https://www.ncbi.nlm.nih.gov/pubmed/25389901. doi:10.3390/nu6115018

Gribble, F. M., & Reimann, F. (2016). Enteroendocrine Cells: Chemosensors in the Intestinal Epithelium. Annu Rev Physiol, 78, 277–299. Retrieved from https://www.ncbi.nlm.nih.gov/pubmed/26442437. doi:10.1146/annurev-physiol-021115-105439

Hara, T., Hirasawa, A., Ichimura, A., Kimura, I., & Tsujimoto, G. (2011). Free fatty acid receptors FFAR1 and GPR120 as novel therapeutic targets for metabolic disorders. J Pharm Sci, 100(9), 3594–3601. Retrieved from https://www.ncbi.nlm.nih.gov/pubmed/21618241. doi:10.1002/jps.22639

Hein, G. J., Baker, C., Hsieh, J., Farr, S., & Adeli, K. (2013). GLP-1 and GLP-2 as yin and yang of intestinal lipoprotein production: evidence for predominance of GLP-2-stimulated postprandial lipemia in normal and insulin-resistant states. Diabetes, 62(2), 373–381. Retrieved from https://www.ncbi.nlm.nih.gov/pubmed/23028139. doi:10.2337/db12-0202

Hetz, C. (2012). The unfolded protein response: controlling cell fate decisions under ER stress and beyond. Nat Rev Mol Cell Biol, 13(2), 89–102. Retrieved from https://www.ncbi.nlm.nih.gov/pubmed/22251901. doi:10.1038/nrm3270

Hildebrandt, M. A., Hoffmann, C., Sherrill-Mix, S. A., Keilbaugh, S. A., Hamady, M., Chen, Y. Y., … Wu, G. D. (2009). High-fat diet determines the composition of the murine gut microbiome independently of obesity. Gastroenterology, 137(5), 1716–1724 e1711-1712. Retrieved from https://www.ncbi.nlm.nih.gov/pubmed/19706296. doi:10.1053/j.gastro.2009.08.042

Hill, J. O., Hauptman, J., Anderson, J. W., Fujioka, K., O’Neil, P. M., Smith, D. K., … Aronne, L. J. (1999). Orlistat, a lipase inhibitor, for weight maintenance after conventional dieting: a 1-y study. Am J Clin Nutr, 69(6), 1108–1116. Retrieved from https://www.ncbi.nlm.nih.gov/pubmed/10357727. doi:10.1093/ajcn/69.6.1108

Hirasawa, A., Tsumaya, K., Awaji, T., Katsuma, S., Adachi, T., Yamada, M., … Tsujimoto, G. (2005). Free fatty acids regulate gut incretin glucagon-like peptide-1 secretion through GPR120. Nat Med, 11(1), 90–94. Retrieved from https://www.ncbi.nlm.nih.gov/pubmed/15619630. doi:10.1038/nm1168

Hofer, D., Asan, E., & Drenckhahn, D. (1999). Chemosensory Perception in the Gut. News Physiol Sci, 14, 18–23. Retrieved from https://www.ncbi.nlm.nih.gov/pubmed/11390812.

Hsieh, J., Longuet, C., Maida, A., Bahrami, J., Xu, E., Baker, C. L., … Adeli, K. (2009). Glucagon-like peptide-2 increases intestinal lipid absorption and chylomicron production via CD36. Gastroenterology, 137(3), 997–1005, 1005 e1001-1004. Retrieved from https://www.ncbi.nlm.nih.gov/pubmed/19482026. doi:10.1053/j.gastro.2009.05.051

Hu, S., Wang, L., Yang, D., Li, L., Togo, J., Wu, Y., … Speakman, J. R. (2018). Dietary Fat, but Not Protein or Carbohydrate, Regulates Energy Intake and Causes Adiposity in Mice. Cell Metab, 28(3), 415–431 e414. Retrieved from https://www.ncbi.nlm.nih.gov/pubmed/30017356. doi:10.1016/j.cmet.2018.06.010

Ishikawa, Y., Eguchi, T., & Ishida, H. (1997). Mechanism of beta-adrenergic agonist-induced transmural transport of glucose in rat small intestine. Regulation of phosphorylation of SGLT1 controls the function. Biochim Biophys Acta, 1357(3), 306–318. Retrieved from https://www.ncbi.nlm.nih.gov/pubmed/9268055.

Jha, C., Ghosh, S., Gautam, V., Malhotra, P., & Ray, P. (2017). In vitro study of virulence potential of Acinetobacter baumannii outer membrane vesicles. Microb Pathog, 111, 218–224. Retrieved from https://www.ncbi.nlm.nih.gov/pubmed/28870696. doi:10.1016/j.micpath.2017.08.048

Jin, J. S., Kwon, S. O., Moon, D. C., Gurung, M., Lee, J. H., Kim, S. I., & Lee, J. C. (2011). Acinetobacter baumannii secretes cytotoxic outer membrane protein A via outer membrane vesicles. PLoS One, 6(2), e17027. Retrieved from https://www.ncbi.nlm.nih.gov/pubmed/21386968. doi:10.1371/journal.pone.0017027

Jun, S. H., Lee, J. H., Kim, B. R., Kim, S. I., Park, T. I., Lee, J. C., & Lee, Y. C. (2013). Acinetobacter baumannii outer membrane vesicles elicit a potent innate immune response via membrane proteins. PLoS One, 8(8), e71751. Retrieved from https://www.ncbi.nlm.nih.gov/pubmed/23977136. doi:10.1371/journal.pone.0071751

Kaelberer, M. M., Buchanan, K. L., Klein, M. E., Barth, B. B., Montoya, M. M., Shen, X., & Bohorquez, D. V. (2018). A gut-brain neural circuit for nutrient sensory transduction. Science, 361(6408). Retrieved from https://www.ncbi.nlm.nih.gov/pubmed/30237325. doi:10.1126/science.aat5236

Kahn, S. E., Hull, R. L., & Utzschneider, K. M. (2006). Mechanisms linking obesity to insulin resistance and type 2 diabetes. Nature, 444(7121), 840–846. Retrieved from https://www.ncbi.nlm.nih.gov/pubmed/17167471. doi:10.1038/nature05482

Kanamori, T., Kanai, M. I., Dairyo, Y., Yasunaga, K., Morikawa, R. K., & Emoto, K. (2013). Compartmentalized calcium transients trigger dendrite pruning in Drosophila sensory neurons. Science, 340(6139), 1475–1478. Retrieved from https://www.ncbi.nlm.nih.gov/pubmed/23722427. doi:10.1126/science.1234879

Kanther, M., Sun, X., Muhlbauer, M., Mackey, L. C., Flynn, E. J., 3rd, Bagnat, M., … Rawls, J. F. (2011). Microbial colonization induces dynamic temporal and spatial patterns of NF-kappaB activation in the zebrafish digestive tract. Gastroenterology, 141(1), 197–207. Retrieved from https://www.ncbi.nlm.nih.gov/pubmed/21439961. doi:10.1053/j.gastro.2011.03.042

Kanwal, Z., Wiegertjes, G. F., Veneman, W. J., Meijer, A. H., & Spaink, H. P. (2014). Comparative studies of Toll-like receptor signalling using zebrafish. Dev Comp Immunol, 46(1), 35–52. Retrieved from https://www.ncbi.nlm.nih.gov/pubmed/24560981. doi:10.1016/j.dci.2014.02.003

Katsuma, S., Hatae, N., Yano, T., Ruike, Y., Kimura, M., Hirasawa, A., & Tsujimoto, G. (2005). Free fatty acids inhibit serum deprivation-induced apoptosis through GPR120 in a murine enteroendocrine cell line STC-1. J Biol Chem, 280(20), 19507–19515. Retrieved from https://www.ncbi.nlm.nih.gov/pubmed/15774482. doi:10.1074/jbc.M412385200

Kawakami, K. (2007). Tol2: a versatile gene transfer vector in vertebrates. Genome Biol, 8 *Suppl 1*, S7. Retrieved from https://www.ncbi.nlm.nih.gov/pubmed/18047699. doi:10.1186/gb-2007-8-s1-s7

Kawasaki, T., & Kawai, T. (2014). Toll-like receptor signaling pathways. Front Immunol, 5, 461. Retrieved from https://www.ncbi.nlm.nih.gov/pubmed/25309543. doi:10.3389/fimmu.2014.00461

Kay, R. J., Boissy, R. J., Russnak, R. H., & Candido, E. P. (1986). Efficient transcription of a Caenorhabditis elegans heat shock gene pair in mouse fibroblasts is dependent on multiple promoter elements which can function bidirectionally. Mol Cell Biol, 6(9), 3134–3143. Retrieved from https://www.ncbi.nlm.nih.gov/pubmed/3023964.

Kieffer, T. J., McIntosh, C. H., & Pederson, R. A. (1995). Degradation of glucose-dependent insulinotropic polypeptide and truncated glucagon-like peptide 1 in vitro and in vivo by dipeptidyl peptidase IV. Endocrinology, 136(8), 3585–3596. Retrieved from https://www.ncbi.nlm.nih.gov/pubmed/7628397. doi:10.1210/endo.136.8.7628397

Kim, S., Joe, Y., Kim, H. J., Kim, Y. S., Jeong, S. O., Pae, H. O., … Chung, H. T. (2015). Endoplasmic reticulum stress-induced IRE1alpha activation mediates cross-talk of GSK-3beta and XBP-1 to regulate inflammatory cytokine production. J Immunol, 194(9), 4498–4506. Retrieved from https://www.ncbi.nlm.nih.gov/pubmed/25821218. doi:10.4049/jimmunol.1401399

Klausner, R. D., Donaldson, J. G., & Lippincott-Schwartz, J. (1992). Brefeldin A: insights into the control of membrane traffic and organelle structure. J Cell Biol, 116(5), 1071–1080. Retrieved from https://www.ncbi.nlm.nih.gov/pubmed/1740466.

Kwan, K. M., Fujimoto, E., Grabher, C., Mangum, B. D., Hardy, M. E., Campbell, D. S., … Chien, C. B. (2007). The Tol2kit: a multisite gateway-based construction kit for Tol2 transposon transgenesis constructs. Dev Dyn, 236(11), 3088–3099. Retrieved from https://www.ncbi.nlm.nih.gov/pubmed/17937395. doi:10.1002/dvdy.21343

Lal, B., & Khanna, S. (1996). Degradation of crude oil by Acinetobacter calcoaceticus and Alcaligenes odorans. J Appl Bacteriol, 81(4), 355–362. Retrieved from https://www.ncbi.nlm.nih.gov/pubmed/8896350.

Latorre, R., Sternini, C., De Giorgio, R., & Greenwood-Van Meerveld, B. (2016). Enteroendocrine cells: a review of their role in brain-gut communication. Neurogastroenterol Motil, 28(5), 620–630. Retrieved from https://www.ncbi.nlm.nih.gov/pubmed/26691223. doi:10.1111/nmo.12754

Lauffer, L. M., Iakoubov, R., & Brubaker, P. L. (2009). GPR119 is essential for oleoylethanolamide-induced glucagon-like peptide-1 secretion from the intestinal enteroendocrine L-cell. Diabetes, 58(5), 1058–1066. Retrieved from https://www.ncbi.nlm.nih.gov/pubmed/19208912. doi:10.2337/db08-1237

Lee, C. R., Lee, J. H., Park, M., Park, K. S., Bae, I. K., Kim, Y. B., … Lee, S. H. (2017). Biology of Acinetobacter baumannii: Pathogenesis, Antibiotic Resistance Mechanisms, and Prospective Treatment Options. Front Cell Infect Microbiol, 7, 55. Retrieved from https://www.ncbi.nlm.nih.gov/pubmed/28348979. doi:10.3389/fcimb.2017.00055

Li, H. J., Kapoor, A., Giel-Moloney, M., Rindi, G., & Leiter, A. B. (2012). Notch signaling differentially regulates the cell fate of early endocrine precursor cells and their maturing descendants in the mouse pancreas and intestine. Dev Biol, 371(2), 156–169. Retrieved from https://www.ncbi.nlm.nih.gov/pubmed/22964416. doi:10.1016/j.ydbio.2012.08.023

Li, H. J., Ray, S. K., Singh, N. K., Johnston, B., & Leiter, A. B. (2011). Basic helix-loop-helix transcription factors and enteroendocrine cell differentiation. Diabetes Obes Metab, 13 *Suppl 1*, 5–12. Retrieved from https://www.ncbi.nlm.nih.gov/pubmed/21824251. doi:10.1111/j.1463-1326.2011.01438.x

Li, J., Chen, Z., Gao, L. Y., Colorni, A., Ucko, M., Fang, S., & Du, S. J. (2015). A transgenic zebrafish model for monitoring xbp1 splicing and endoplasmic reticulum stress in vivo. Mech Dev, 137, 33–44. Retrieved from https://www.ncbi.nlm.nih.gov/pubmed/25892297. doi:10.1016/j.mod.2015.04.001

Li, Z., Wen, C., Peng, J., Korzh, V., & Gong, Z. (2009). Generation of living color transgenic zebrafish to trace somatostatin-expressing cells and endocrine pancreas organization. Differentiation, 77(2), 128–134. Retrieved from https://www.ncbi.nlm.nih.gov/pubmed/19281772. doi:10.1016/j.diff.2008.09.014

Lickwar, C. R., Camp, J. G., Weiser, M., Cocchiaro, J. L., Kingsley, D. M., Furey, T. S., … Rawls, J. F. (2017). Genomic dissection of conserved transcriptional regulation in intestinal epithelial cells. PLoS Biol, 15(8), e2002054. Retrieved from https://www.ncbi.nlm.nih.gov/pubmed/28850571. doi:10.1371/journal.pbio.2002054

Liddle, R. A. (1997). Cholecystokinin cells. Annu Rev Physiol, 59, 221–242. Retrieved from https://www.ncbi.nlm.nih.gov/pubmed/9074762. doi:10.1146/annurev.physiol.59.1.221

Ludwig, D. S., Willett, W. C., Volek, J. S., & Neuhouser, M. L. (2018). Dietary fat: From foe to friend? Science, 362(6416), 764–770. Retrieved from https://www.ncbi.nlm.nih.gov/pubmed/30442800. doi:10.1126/science.aau2096

Mao, S., Zhang, M., Liu, J., & Zhu, W. (2015). Characterising the bacterial microbiota across the gastrointestinal tracts of dairy cattle: membership and potential function. Sci Rep, 5, 16116. Retrieved from https://www.ncbi.nlm.nih.gov/pubmed/26527325. doi:10.1038/srep16116

March, C., Regueiro, V., Llobet, E., Moranta, D., Morey, P., Garmendia, J., & Bengoechea, J. A. (2010). Dissection of host cell signal transduction during Acinetobacter baumannii-triggered inflammatory response. PLoS One, 5(4), e10033. Retrieved from https://www.ncbi.nlm.nih.gov/pubmed/20383325. doi:10.1371/journal.pone.0010033

Martinez-Guryn, K., Hubert, N., Frazier, K., Urlass, S., Musch, M. W., Ojeda, P., … Chang, E. B. (2018). Small Intestine Microbiota Regulate Host Digestive and Absorptive Adaptive Responses to Dietary Lipids. Cell Host Microbe, 23(4), 458–469 e455. Retrieved from https://www.ncbi.nlm.nih.gov/pubmed/29649441. doi:10.1016/j.chom.2018.03.011

McGraw, H. F., Snelson, C. D., Prendergast, A., Suli, A., & Raible, D. W. (2012). Postembryonic neuronal addition in zebrafish dorsal root ganglia is regulated by Notch signaling. Neural Dev, 7, 23. Retrieved from https://www.ncbi.nlm.nih.gov/pubmed/22738203. doi:10.1186/1749-8104-7-23

Moran-Ramos, S., Tovar, A. R., & Torres, N. (2012). Diet: friend or foe of enteroendocrine cells--how it interacts with enteroendocrine cells. Adv Nutr, 3(1), 8–20. Retrieved from https://www.ncbi.nlm.nih.gov/pubmed/22332097. doi:10.3945/an.111.000976

Murdoch, C. C., Espenschied, S. T., Matty, M. A., Mueller, O., Tobin, D. M., & Rawls, J. E (2019). Intestinal Serum amyloid A suppresses systemic neutrophil activation and bactericidal activity in response to microbiota colonization. PLoS Pathog, 15(3), e1007381. Retrieved from https://www.ncbi.nlm.nih.gov/pubmed/30845179. doi:10.1371/journal.ppat.1007381

Murphy, E. F., Cotter, P. D., Healy, S., Marques, T. M., O’Sullivan, O., Fouhy, F., … Shanahan, F. (2010). Composition and energy harvesting capacity of the gut microbiota: relationship to diet, obesity and time in mouse models. Gut, 59(12), 1635–1642. Retrieved from https://www.ncbi.nlm.nih.gov/pubmed/20926643. doi:10.1136/gut.2010.215665

Nakamura, O., Tazumi, Y., Muro, T., Yasuhara, Y., & Watanabe, T. (2004). Active uptake and transport of protein by the intestinal epithelial cells in embryo of viviparous fish, Neoditrema ransonneti (Perciformes: Embiotocidae). J Exp Zool A Comp Exp Biol, 301(1), 38–48. Retrieved from https://www.ncbi.nlm.nih.gov/pubmed/14695687. doi:10.1002/jez.a.20005

Navon-Venezia, S., Zosim, Z., Gottlieb, A., Legmann, R., Carmeli, S., Ron, E. Z., & Rosenberg, E. (1995). Alasan, a new bioemulsifier from Acinetobacter radioresistens. Appl Environ Microbiol, 61(9), 3240–3244. Retrieved from https://www.ncbi.nlm.nih.gov/pubmed/7574633.

Ng, A. N., de Jong-Curtain, T. A., Mawdsley, D. J., White, S. J., Shin, J., Appel, B., … Heath, J. K. (2005). Formation of the digestive system in zebrafish: III. Intestinal epithelium morphogenesis. Dev Biol, 286(1), 114–135. Retrieved from https://www.ncbi.nlm.nih.gov/pubmed/16125164. doi:10.1016/j.ydbio.2005.07.013

Nikolaev, A., McLaughlin, T., O’Leary, D. D., & Tessier-Lavigne, M. (2009). APP binds DR6 to trigger axon pruning and neuron death via distinct caspases. Nature, 457(7232), 981–989. Retrieved from https://www.ncbi.nlm.nih.gov/pubmed/19225519. doi:10.1038/nature07767

Oakes, N. D., Cooney, G. J., Camilleri, S., Chisholm, D. J., & Kraegen, E. W. (1997). Mechanisms of liver and muscle insulin resistance induced by chronic high-fat feeding. Diabetes, 46(11), 1768–1774. Retrieved from https://www.ncbi.nlm.nih.gov/pubmed/9356024.

Okawa, M., Fujii, K., Ohbuchi, K., Okumoto, M., Aragane, K., Sato, H., … Yoshimoto, R. (2009). Role of MGAT2 and DGAT1 in the release of gut peptides after triglyceride ingestion. Biochem Biophys Res Commun, 390(3), 377–381. Retrieved from https://www.ncbi.nlm.nih.gov/pubmed/19732742. doi:10.1016/j.bbrc.2009.08.167

Pahl, H. L., & Baeuerle, P. A. (1997). The ER-overload response: activation of NF-kappa B. Trends Biochem Sci, 22(2), 63–67. Retrieved from https://www.ncbi.nlm.nih.gov/pubmed/9048485.

Palti, Y. (2011). Toll-like receptors in bony fish: from genomics to function. Dev Comp Immunol, 35(12), 1263–1272. Retrieved from https://www.ncbi.nlm.nih.gov/pubmed/21414346. doi:10.1016/j.dci.2011.03.006

Panchal, S. K., Poudyal, H., Iyer, A., Nazer, R., Alam, A., Diwan, V., … Brown, L. (2011). High-carbohydrate high-fat diet-induced metabolic syndrome and cardiovascular remodeling in rats. J Cardiovasc Pharmacol, 57(1), 51–64. Retrieved from https://www.ncbi.nlm.nih.gov/pubmed/20966763. doi:10.1097/FJC.0b013e3181feb90a

Parsons, M. J., Pisharath, H., Yusuff, S., Moore, J. C., Siekmann, A. F., Lawson, N., & Leach, S. D. (2009). Notch-responsive cells initiate the secondary transition in larval zebrafish pancreas. Mech Dev, 126(10), 898–912. Retrieved from https://www.ncbi.nlm.nih.gov/pubmed/19595765. doi:10.1016/j.mod.2009.07.002

Pedron, T., Mulet, C., Dauga, C., Frangeul, L., Chervaux, C., Grompone, G., & Sansonetti, P. J. (2012). A crypt-specific core microbiota resides in the mouse colon. MBio, 3(3). Retrieved from https://www.ncbi.nlm.nih.gov/pubmed/22617141. doi:10.1128/mBio.00116-12

Pham, L. N., Kanther, M., Semova, I., & Rawls, J. F. (2008). Methods for generating and colonizing gnotobiotic zebrafish. Nat Protoc, 3(12), 1862–1875. Retrieved from https://www.ncbi.nlm.nih.gov/pubmed/19008873. doi:10.1038/nprot.2008.186

Phan, C. T., & Tso, P. (2001). Intestinal lipid absorption and transport. Front Biosci, 6, D299–319. Retrieved from https://www.ncbi.nlm.nih.gov/pubmed/11229876.

Poureslami, R., Raes, K., Huyghebaert, G., Batal, A. B., & De Smet, S. (2012). Egg yolk fatty acid profile in relation to dietary fatty acid concentrations. J Sci Food Agric, 92(2), 366–372. Retrieved from https://www.ncbi.nlm.nih.gov/pubmed/21815168. doi:10.1002/jsfa.4587

Quast, C., Pruesse, E., Yilmaz, P., Gerken, J., Schweer, T., Yarza, P., … Glockner, F. O. (2013). The SILVA ribosomal RNA gene database project: improved data processing and web-based tools. Nucleic Acids Res, 41(Database issue), D590–596. Retrieved from https://www.ncbi.nlm.nih.gov/pubmed/23193283. doi:10.1093/nar/gks1219

Quick, M. W., & Lester, R. A. (2002). Desensitization of neuronal nicotinic receptors. J Neurobiol, 53(4), 457–478. Retrieved from https://www.ncbi.nlm.nih.gov/pubmed/12436413. doi:10.1002/neu.10109

Quinlivan, V. H., & Farber, S. A. (2017). Lipid Uptake, Metabolism, and Transport in the Larval Zebrafish. Front Endocrinol (Lausanne), 8, 319. Retrieved from https://www.ncbi.nlm.nih.gov/pubmed/29209275. doi:10.3389/fendo.2017.00319

Rabot, S., Membrez, M., Bruneau, A., Gerard, P., Harach, T., Moser, M., … Chou, C. J. (2010). Germ-free C57BL/6J mice are resistant to high-fat-diet-induced insulin resistance and have altered cholesterol metabolism. FASEB J, 24(12), 4948–4959. Retrieved from https://www.ncbi.nlm.nih.gov/pubmed/20724524. doi:10.1096/fj.10-164921

Rawls, J. F., Mahowald, M. A., Goodman, A. L., Trent, C. M., & Gordon, J. I. (2007). In vivo imaging and genetic analysis link bacterial motility and symbiosis in the zebrafish gut. Proc Natl Acad Sci U S A, 104(18), 7622–7627. Retrieved from https://www.ncbi.nlm.nih.gov/pubmed/17456593. doi:10.1073/pnas.0702386104

Rawls, J. F., Samuel, B. S., & Gordon, J. I. (2004). Gnotobiotic zebrafish reveal evolutionarily conserved responses to the gut microbiota. Proc Natl Acad Sci U S A, 101(13), 4596–4601. Retrieved from https://www.ncbi.nlm.nih.gov/pubmed/15070763. doi:10.1073/pnas.0400706101

Ray, S. K., & Leiter, A. B. (2007). The basic helix-loop-helix transcription factor NeuroD1 facilitates interaction of Sp1 with the secretin gene enhancer. Mol Cell Biol, 27(22), 7839–7847. Retrieved from https://www.ncbi.nlm.nih.gov/pubmed/17875929. doi:10.1128/MCB.00438-07

Ray, S. K., Li, H. J., Metzger, E., Schule, R., & Leiter, A. B. (2014). CtBP and associated LSD1 are required for transcriptional activation by NeuroD1 in gastrointestinal endocrine cells. Mol Cell Biol, 34(12), 2308–2317. Retrieved from https://www.ncbi.nlm.nih.gov/pubmed/24732800. doi:10.1128/MCB.01600-13

Raybould, H. E. (2007). Mechanisms of CCK signaling from gut to brain. Curr Opin Pharmacol, 7(6), 570–574. Retrieved from https://www.ncbi.nlm.nih.gov/pubmed/17954038. doi:10.1016/j.coph.2007.09.006

Reimann, F., Habib, A. M., Tolhurst, G., Parker, H. E., Rogers, G. J., & Gribble, F. M. (2008). Glucose sensing in L cells: a primary cell study. Cell Metab, 8(6), 532–539. Retrieved from https://www.ncbi.nlm.nih.gov/pubmed/19041768. doi:10.1016/j.cmet.2008.11.002

Richards, P., Pais, R., Habib, A. M., Brighton, C. A., Yeo, G. S., Reimann, F., & Gribble, F. M. (2016). High fat diet impairs the function of glucagon-like peptide-1 producing L-cells. Peptides, 77, 21–27. Retrieved from https://www.ncbi.nlm.nih.gov/pubmed/26145551. doi:10.1016/j.peptides.2015.06.006

Ridaura, V. K., Faith, J. J., Rey, F. E., Cheng, J., Duncan, A. E., Kau, A. L., … Gordon, J. I. (2013). Gut microbiota from twins discordant for obesity modulate metabolism in mice. Science, 341(6150), 1241214. Retrieved from https://www.ncbi.nlm.nih.gov/pubmed/24009397. doi:10.1126/science.1241214

Riedl, J., Crevenna, A. H., Kessenbrock, K., Yu, J. H., Neukirchen, D., Bista, M., … Wedlich-Soldner, R. (2008). Lifeact: a versatile marker to visualize F-actin. Nat Methods, 5(7), 605–607. Retrieved from https://www.ncbi.nlm.nih.gov/pubmed/18536722. doi:10.1038/nmeth.1220

Rombout, J. H., Lamers, C. H., & Hanstede, J. G. (1978). Enteroendocrine APUD cells in the digestive tract of larval Barbus conchonius (Teleostei, Cyprinidae). J Embryol Exp Morphol, 47, 121–135. Retrieved from https://www.ncbi.nlm.nih.gov/pubmed/31409.

Rupprecht, P., Prendergast, A., Wyart, C., & Friedrich, R. W. (2016). Remote z-scanning with a macroscopic voice coil motor for fast 3D multiphoton laser scanning microscopy. Biomed Opt Express, 7(5), 1656–1671. Retrieved from https://www.ncbi.nlm.nih.gov/pubmed/27231612. doi:10.1364/BOE.7.001656

Saffarian, A., Touchon, M., Mulet, C., Tournebize, R., Passet, V., Brisse, S., … Pedron, T. (2017). Comparative genomic analysis of Acinetobacter strains isolated from murine colonic crypts. BMC Genomics, 18(1), 525. Retrieved from https://www.ncbi.nlm.nih.gov/pubmed/28697749. doi:10.1186/s12864-017-3925-x

Sagasti, A., Guido, M. R., Raible, D. W., & Schier, A. F. (2005). Repulsive interactions shape the morphologies and functional arrangement of zebrafish peripheral sensory arbors. Curr Biol, 15(9), 804–814. Retrieved from https://www.ncbi.nlm.nih.gov/pubmed/15886097. doi:10.1016/j.cub.2005.03.048

Samali, A., Fitzgerald, U., Deegan, S., & Gupta, S. (2010). Methods for monitoring endoplasmic reticulum stress and the unfolded protein response. Int J Cell Biol, 2010, 830307. Retrieved from https://www.ncbi.nlm.nih.gov/pubmed/20169136. doi:10.1155/2010/830307

Sandoval, D. A., & D’Alessio, D. A. (2015). Physiology of proglucagon peptides: role of glucagon and GLP-1 in health and disease. Physiol Rev, 95(2), 513–548. Retrieved from https://www.ncbi.nlm.nih.gov/pubmed/25834231. doi:10.1152/physrev.00013.2014

Scott T., Espenschied, M. R. C., Molly A. Matty, Olaf Mueller, Matthew R. Redinbod, David M. Tobina, and John F. Rawls. (2019). Epithelial delamination is protective during pharmaceutical-induced enteropathy. In Revision.

Segata, N., Izard, J., Waldron, L., Gevers, D., Miropolsky, L., Garrett, W. S., & Huttenhower, C. (2011). Metagenomic biomarker discovery and explanation. Genome Biol, 12(6), R60. Retrieved from https://www.ncbi.nlm.nih.gov/pubmed/21702898. doi:10.1186/gb-2011-12-6-r60

Semova, I., Carten, J. D., Stombaugh, J., Mackey, L. C., Knight, R., Farber, S. A., & Rawls, J. F. (2012). Microbiota regulate intestinal absorption and metabolism of fatty acids in the zebrafish. Cell Host Microbe, 12(3), 277–288. Retrieved from https://www.ncbi.nlm.nih.gov/pubmed/22980325. doi:10.1016/j.chom.2012.08.003

Shimotoyodome, A., Fukuoka, D., Suzuki, J., Fujii, Y., Mizuno, T., Meguro, S., … Hase, T. (2009). Coingestion of acylglycerols differentially affects glucose-induced insulin secretion via glucose-dependent insulinotropic polypeptide in C57BL/6J mice. Endocrinology, 150(5), 2118–2126. Retrieved from https://www.ncbi.nlm.nih.gov/pubmed/19179446. doi:10.1210/en.2008-1162

Snellman, E. A., & Colwell, R. R. (2004). Acinetobacter lipases: molecular biology, biochemical properties and biotechnological potential. J Ind Microbiol Biotechnol, 31(9), 391–400. Retrieved from https://www.ncbi.nlm.nih.gov/pubmed/15378387. doi:10.1007/s10295-004-0167-0

Song, P., Onishi, A., Koepsell, H., & Vallon, V. (2016). Sodium glucose cotransporter SGLT1 as a therapeutic target in diabetes mellitus. Expert Opin Ther Targets, 20(9), 1109–1125. Retrieved from https://www.ncbi.nlm.nih.gov/pubmed/26998950. doi:10.1517/14728222.2016.1168808

Songer, J. G. (1997). Bacterial phospholipases and their role in virulence. Trends Microbiol, 5(4), 156–161. Retrieved from https://www.ncbi.nlm.nih.gov/pubmed/9141190. doi:10.1016/S0966-842X(97)01005-6

Stephens, W. Z., Burns, A. R., Stagaman, K., Wong, S., Rawls, J. F., Guillemin, K., & Bohannan, B. J. (2016). The composition of the zebrafish intestinal microbial community varies across development. ISME J, 10(3), 644–654. Retrieved from https://www.ncbi.nlm.nih.gov/pubmed/26339860. doi:10.1038/ismej.2015.140

Sternini, C., Anselmi, L., & Rozengurt, E. (2008). Enteroendocrine cells: a site of ‘taste’ in gastrointestinal chemosensing. Curr Opin Endocrinol Diabetes Obes, 15(1), 73–78. Retrieved from https://www.ncbi.nlm.nih.gov/pubmed/18185066. doi:10.1097/MED.0b013e3282f43a73

Subramanian, S., Glitz, P., Kipp, H., Kinne, R. K., & Castaneda, F. (2009). Protein kinase-A affects sorting and conformation of the sodium-dependent glucose co-transporter SGLT1. J Cell Biochem, 106(3), 444–452. Retrieved from https://www.ncbi.nlm.nih.gov/pubmed/19115253. doi:10.1002/jcb.22025

Toren, A., Navon-Venezia, S., Ron, E. Z., & Rosenberg, E. (2001). Emulsifying activities of purified Alasan proteins from Acinetobacter radioresistens KA53. Appl Environ Microbiol, 67(3), 1102–1106. Retrieved from https://www.ncbi.nlm.nih.gov/pubmed/11229898. doi:10.1128/AEM.67.3.1102-1106.2001

Trapani, J. G., Obholzer, N., Mo, W., Brockerhoff, S. E., & Nicolson, T. (2009). Synaptojanin1 is required for temporal fidelity of synaptic transmission in hair cells. PLoS Genet, 5(5), e1000480. Retrieved from https://www.ncbi.nlm.nih.gov/pubmed/19424431. doi:10.1371/journal.pgen.1000480

Troll, J. V., Hamilton, M. K., Abel, M. L., Ganz, J., Bates, J. M., Stephens, W. Z., … Guillemin, K. (2018). Microbiota promote secretory cell determination in the intestinal epithelium by modulating host Notch signaling. Development, 145(4). Retrieved from https://www.ncbi.nlm.nih.gov/pubmed/29475973. doi:10.1242/dev.155317

Tseng, Q., Duchemin-Pelletier, E., Deshiere, A., Balland, M., Guillou, H., Filhol, O., & Thery, M. (2012). Spatial organization of the extracellular matrix regulates cell-cell junction positioning. Proc Natl Acad Sci U S A, 109(5), 1506–1511. Retrieved from https://www.ncbi.nlm.nih.gov/pubmed/22307605. doi:10.1073/pnas.1106377109

Turnbaugh, P. J., Backhed, F., Fulton, L., & Gordon, J. I. (2008). Diet-induced obesity is linked to marked but reversible alterations in the mouse distal gut microbiome. Cell Host Microbe, 3(4), 213–223. Retrieved from https://www.ncbi.nlm.nih.gov/pubmed/18407065. doi:10.1016/j.chom.2008.02.015

Turnbaugh, P. J., Ley, R. E., Mahowald, M. A., Magrini, V., Mardis, E. R., & Gordon, J. I. (2006). An obesity-associated gut microbiome with increased capacity for energy harvest. Nature, 444(7122), 1027–1031. Retrieved from https://www.ncbi.nlm.nih.gov/pubmed/17183312. doi:10.1038/nature05414

Uppala, J. K., Gani, A. R., & Ramaiah, K. V. A. (2017). Chemical chaperone, TUDCA unlike PBA, mitigates protein aggregation efficiently and resists ER and non-ER stress induced HepG2 cell death. Sci Rep, 7(1), 3831. Retrieved from https://www.ncbi.nlm.nih.gov/pubmed/28630443. doi:10.1038/s41598-017-03940-1

Vang, S., Longley, K., Steer, C. J., & Low, W. C. (2014). The Unexpected Uses of Urso-and Tauroursodeoxycholic Acid in the Treatment of Non-liver Diseases. Glob Adv Health Med, 3(3), 58–69. Retrieved from https://www.ncbi.nlm.nih.gov/pubmed/24891994. doi:10.7453/gahmj.2014.017

Walker, C. S., Jensen, S., Ellison, M., Matta, J. A., Lee, W. Y., Imperial, J. S., … Maricq, A. V. (2009). A novel Conus snail polypeptide causes excitotoxicity by blocking desensitization of AMPA receptors. Curr Biol, 19(11), 900–908. Retrieved from https://www.ncbi.nlm.nih.gov/pubmed/19481459. doi:10.1016/j.cub.2009.05.017

Wallace, K. N., Akhter, S., Smith, E. M., Lorent, K., & Pack, M. (2005). Intestinal growth and differentiation in zebrafish. Mech Dev, 122(2), 157–173. Retrieved from https://www.ncbi.nlm.nih.gov/pubmed/15652704. doi:10.1016/j.mod.2004.10.009

Wallace, K. N., & Pack, M. (2003). Unique and conserved aspects of gut development in zebrafish. Dev Biol, 255(1), 12–29. Retrieved from https://www.ncbi.nlm.nih.gov/pubmed/12618131.

Walzer, G., Rosenberg, E., & Ron, E. Z. (2006). The Acinetobacter outer membrane protein A (OmpA) is a secreted emulsifier. Environ Microbiol, 8(6), 1026–1032. Retrieved from https://www.ncbi.nlm.nih.gov/pubmed/16689723. doi:10.1111/j.1462-2920.2006.00994.x

Wang, Z., Du, J., Lam, S. H., Mathavan, S., Matsudaira, P., & Gong, Z. (2010). Morphological and molecular evidence for functional organization along the rostrocaudal axis of the adult zebrafish intestine. BMC Genomics, 11, 392. Retrieved from https://www.ncbi.nlm.nih.gov/pubmed/20565988. doi:10.1186/1471-2164-11-392

Williams, D. W., Kondo, S., Krzyzanowska, A., Hiromi, Y., & Truman, J. W. (2006). Local caspase activity directs engulfment of dendrites during pruning. Nat Neurosci, 9(10), 1234–1236. Retrieved from https://www.ncbi.nlm.nih.gov/pubmed/16980964. doi:10.1038/nn1774

Williams, D. W., & Truman, J. W. (2005). Cellular mechanisms of dendrite pruning in Drosophila: insights from in vivo time-lapse of remodeling dendritic arborizing sensory neurons. Development, 132(16), 3631–3642. Retrieved from https://www.ncbi.nlm.nih.gov/pubmed/16033801. doi:10.1242/dev.01928

Wong, S., Stephens, W. Z., Burns, A. R., Stagaman, K., David, L. A., Bohannan, B. J., … Rawls, J. F. (2015). Ontogenetic Differences in Dietary Fat Influence Microbiota Assembly in the Zebrafish Gut. MBio, 6(5), e00687–00615. Retrieved from https://www.ncbi.nlm.nih.gov/pubmed/26419876. doi:10.1128/mBio.00687-15

Wright, E. M., Hirsch, J. R., Loo, D. D., & Zampighi, G. A. (1997). Regulation of Na+/glucose cotransporters. J Exp Biol, 200(Pt 2), 287–293. Retrieved from https://www.ncbi.nlm.nih.gov/pubmed/9050236.

Xu, C., Bailly-Maitre, B., & Reed, J. C. (2005). Endoplasmic reticulum stress: cell life and death decisions. J Clin Invest, 115(10), 2656–2664. Retrieved from https://www.ncbi.nlm.nih.gov/pubmed/16200199. doi:10.1172/JCI26373

Ye, L., Robertson, M. A., Hesselson, D., Stainier, D. Y., & Anderson, R. M. (2015). Glucagon is essential for alpha cell transdifferentiation and beta cell neogenesis. Development, 142(8), 1407–1417. Retrieved from https://www.ncbi.nlm.nih.gov/pubmed/25852199. doi:10.1242/dev.117911

Yoshida, H., Matsui, T., Yamamoto, A., Okada, T., & Mori, K. (2001). XBP1 mRNA is induced by ATF6 and spliced by IRE1 in response to ER stress to produce a highly active transcription factor. Cell, 107(7), 881–891. Retrieved from https://www.ncbi.nlm.nih.gov/pubmed/11779464.

Yu, F., & Schuldiner, O. (2014). Axon and dendrite pruning in Drosophila. Curr Opin Neurobiol, 27, 192–198. Retrieved from https://www.ncbi.nlm.nih.gov/pubmed/24793180. doi:10.1016/j.conb.2014.04.005

